# Calling Cards: a customizable platform to longitudinally record protein-DNA interactions over time in cells and tissues

**DOI:** 10.1101/2023.06.07.544098

**Authors:** Allen Yen, Chase Mateusiak, Simona Sarafinovska, Mariam A. Gachechiladze, Juanru Guo, Xuhua Chen, Arnav Moudgil, Alexander J. Cammack, Jessica Hoisington-Lopez, MariaLynn Crosby, Michael R. Brent, Robi D. Mitra, Joseph D. Dougherty

**Affiliations:** Department of Genetics, Washington University School of Medicine, Saint Louis, MO 63110; Department of Psychiatry, Washington University School of Medicine, Saint Louis, MO 63110; Department of Neurology, Washington University School of Medicine, Saint Louis, MO 63110; Edison Family Center for Genome Sciences and Systems Biology, Washington University School of Medicine, Saint Louis, MO 63110; Department of Computer Science and Engineering, Washington University, Saint Louis, MO 63130; Intellectual and Developmental Disabilities Research Center, Washington University School of Medicine, Saint Louis, MO 63110

**Keywords:** Epigenetics, transcription factor binding, enhancer usage, gene regulation, Calling Cards

## Abstract

Calling Cards is a platform technology to record a cumulative history of transient protein-DNA interactions in the genome of genetically targeted cell types. The record of these interactions is recovered by next generation sequencing. Compared to other genomic assays, whose readout provides a snapshot at the time of harvest, Calling Cards enables correlation of historical molecular states to eventual outcomes or phenotypes. To achieve this, Calling Cards uses the piggyBac transposase to insert self-reporting transposon (SRT) “Calling Cards” into the genome, leaving permanent marks at interaction sites. Calling Cards can be deployed in a variety of *in vitro* and *in vivo* biological systems to study gene regulatory networks involved in development, aging, and disease. Out of the box, it assesses enhancer usage but can be adapted to profile specific transcription factor binding with custom transcription factor (TF)-piggyBac fusion proteins. The Calling Cards workflow has five main stages: delivery of Calling Card reagents, sample preparation, library preparation, sequencing, and data analysis. Here, we first present a comprehensive guide for experimental design, reagent selection, and optional customization of the platform to study additional TFs. Then, we provide an updated protocol for the five steps, using reagents that improve throughput and decrease costs, including an overview of a newly deployed computational pipeline. This protocol is designed for users with basic molecular biology experience to process samples into sequencing libraries in 1-2 days. Familiarity with bioinformatic analysis and command line tools is required to set up the pipeline in a high-performance computing environment and to conduct downstream analyses.

Basic Protocol 1: Preparation and delivery of Calling Cards reagents
Basic Protocol 2: Sample preparation
Basic Protocol 3: Sequencing library preparation
Basic Protocol 4: Library pooling and sequencing
Basic Protocol 5: Data analysis

## INTRODUCTION

Transcription factors (TFs) and DNA regulatory elements interact to drive proper spatial and temporal patterns of gene expression. Transcriptional dysregulation caused by mutations in TFs or regulatory elements can result in disease (reviewed in (Lee and Young, 2013; Chatterjee and Ahituv, 2017)). In addition, transcriptional and epigenetic changes are often studied to understand disease processes, even when the disease is driven by other causes. Next generation sequencing has enabled genome-wide analysis of protein-DNA interactions by chromatin immunoprecipitation followed by sequencing (ChIP-seq), however, ChIP-seq requires high quality antibodies, relatively large amounts of starting material, and only provides a snapshot of states at the time of harvest. Thus, without performing a time course experiment, one cannot attribute historical molecular events with eventual cell states. Furthermore, the need for large amounts of starting material precludes the widespread use of these approaches in specific cell types in complex tissues. Newer immunotethering approaches such as CUT&RUN and CUT&TAG enable experiments with less input material and lower sequencing depths, but the dependency on antibodies remains and is not easily adaptable to query targeted cell populations. To address these limitations, we developed Calling Cards, a customizable platform that records protein-DNA interactions over time using a genetically encoded system. Calling Cards can be adapted and deployed in cell lines and tissues across different biological contexts, without the use of antibodies, and in specific, genetically targeted cell types.

Calling Cards relies upon two key components: a TF-piggyBac transposase fusion and a self-reporting transposon (SRT), which is a piggyBac transposon that contains a tdTomato reporter. When tethered to a TF, the piggyBac transposase inserts SRTs into the genome near TF binding sites, leaving permanent “Calling Cards”, which can then be recovered by sequencing and mapped with base pair resolution (**Figure 1A**). Use of the hyperactive piggyBac (hyPB) increased the overall number of transposition events while maintaining a similar insertion pattern (Yusa et al., 2011; Moudgil et al., 2020b). This accumulation of Calling Card insertions provides a cumulative recording of TF binding over the assayed time period. Examples of Calling Cards data are shown in **Figure 1B-F**. A TF of interest can be assayed through cloning TF-transposase fusion proteins. As an alternative to using TF-fusions, the naive piggyBac transposase can be leveraged for its natural affinity for BRD4, a TF that recognizes acetylated lysine residues and found to be highly enriched in super enhancers (Yoshida et al., 2017), and shown to be important in driving transcription of genes that define cell identity (Wang et al., 2008, 2012). Thus, unfused piggyBac can be used to record BRD4-bound enhancer usage.

**Figure 1:**
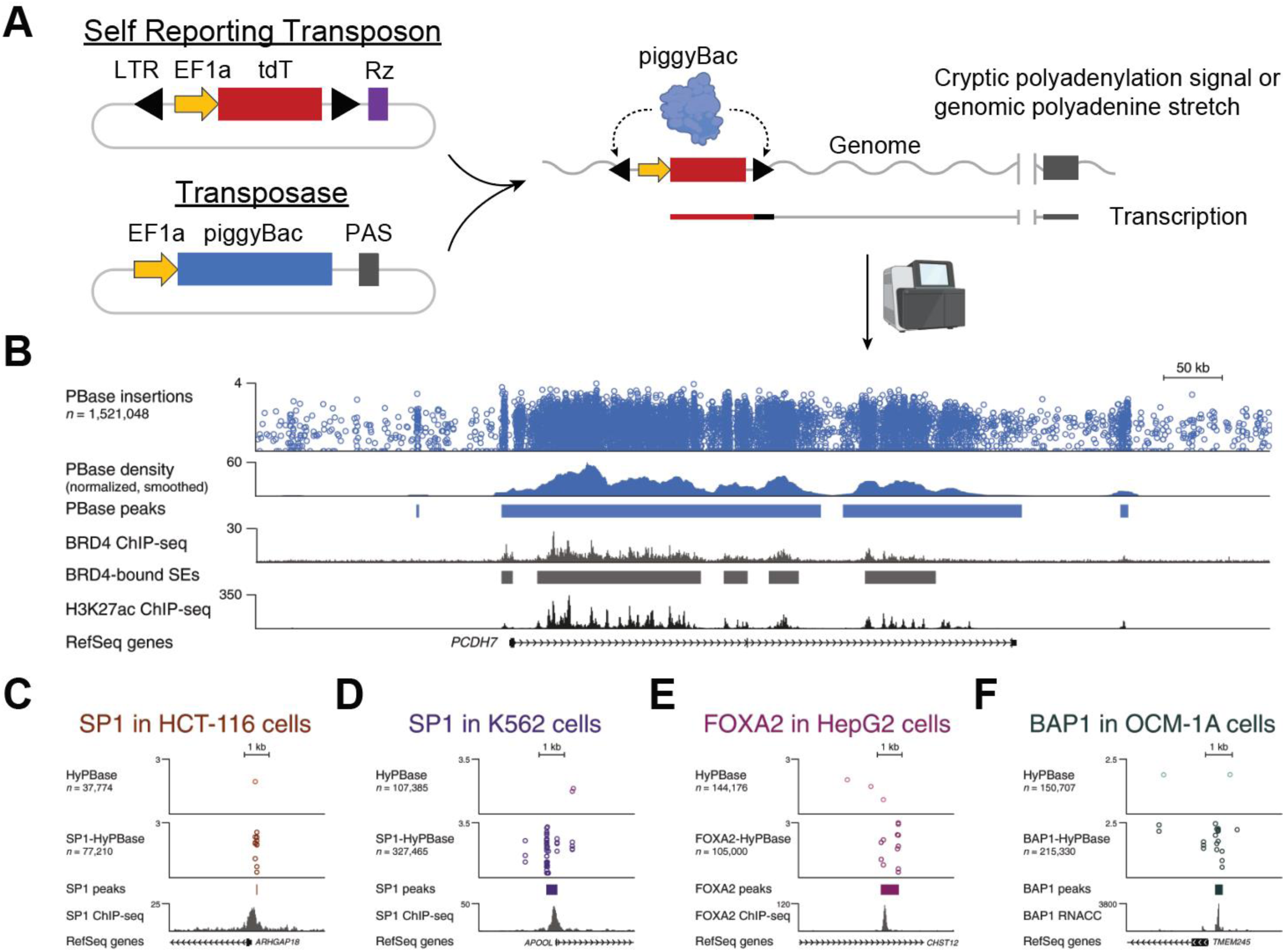
Example tracks showing recording of BRD4-bound super enhancers and TF binding sites using Calling Cards. **(A)** Diagram of the self reporting transposon (SRT) and piggyBac transposase constructs. When expressed in cells, the piggyBac transposase inserts the SRT into the genome at sites of protein-DNA interaction leaving a permanent mark, or Calling Card. The location of Calling Cards insertions can be recovered through RNA sequencing. **(B)** The top track shows the genomic locations of SRT insertions in cells transfected with Calling Cards at the PCDH7 locus. The normalized density of Calling Cards correlates with BRD4 and H3K27ac ChIP-seq peaks. **(C-F)** Fusion of hyPB with a variety of TFs works to redirect Calling Cards across different cell lines. This figure is adapted from (Moudgil et al., 2020b).

While Calling Cards can be thought of as an alternative to CUT&RUN or ChIP-seq, its recording feature also enables additional kinds of experimental questions. First, it can cumulatively record protein-DNA interactions over time, providing an integrated snapshot that could replace time series data. Second, it can be used to correlate early enhancer usage to later cell fate decisions. Some examples include: how can a seemingly homogenous cluster of pluripotent stem cells give rise to many different cell types? How can genetically identical organisms have distinct biological responses to the same stimulus? Current standard genomic technologies and assays are inherently destructive, since in order to analyze the molecular state, the harvested cells are destroyed. By recording protein-DNA interactions over time, historical molecular events such as transcription factor binding, or historical epigenetic states can be linked to current cell states. Linking recorded molecular states to eventual outcomes could be broadly applicable to many areas of research, including but not limited to developmental biology, aging, and gene-environment interactions.

In this protocol, we provide a resource to guide researchers, especially those new to genomic assays, to design and execute a Calling Cards experiment. We have created a streamlined workflow (**Figure 2**) to simplify the selection of reagents needed to perform the desired experiment (**Figure 3**), discuss methods to validate the constructs, and provide recommendations for reagent delivery methods with suggested controls. Specifically, Basic Protocol 1 describes how to create a plasmid pool of barcoded SRTs that can be used for *in vitro* transfections or AAV packaging for *in vivo* transductions. It also describes intracerebroventricular injections, a relatively simple and robust method to reliably deliver AAVs directly into the cerebral lateral ventricles and CNS in early postnatal mice. Next, Basic Protocol 2 outlines the steps to harvest RNA from Calling Cards-containing samples. Basic Protocol 3 describes the entire sequencing library preparation protocol that includes first strand synthesis, amplification of Calling Card transcripts, bead cleanup, tagmentation, indexing, and final bead cleanup. It also contains an optional “library density” qPCR assay that assess the relative abundance of Calling Card transcripts in an RNA sample. This can be performed early on to determine if samples should or should not be carried through the entire protocol. This step can minimize unnecessary labor and usage of reagents. Additionally, there are numerous quality control (QC) steps built into the protocol to monitor progress through the procedure. Basic Protocol 4 details library pooling strategies and recommended parameters for next generation sequencing on various Illumina platforms. Lastly, Basic Protocol 5 provides a high-level guide on using the nextflow Calling Cards bioinformatic pipeline to prepare the raw sequencing data into a format that can be used for downstream analysis.

**Figure 2:**
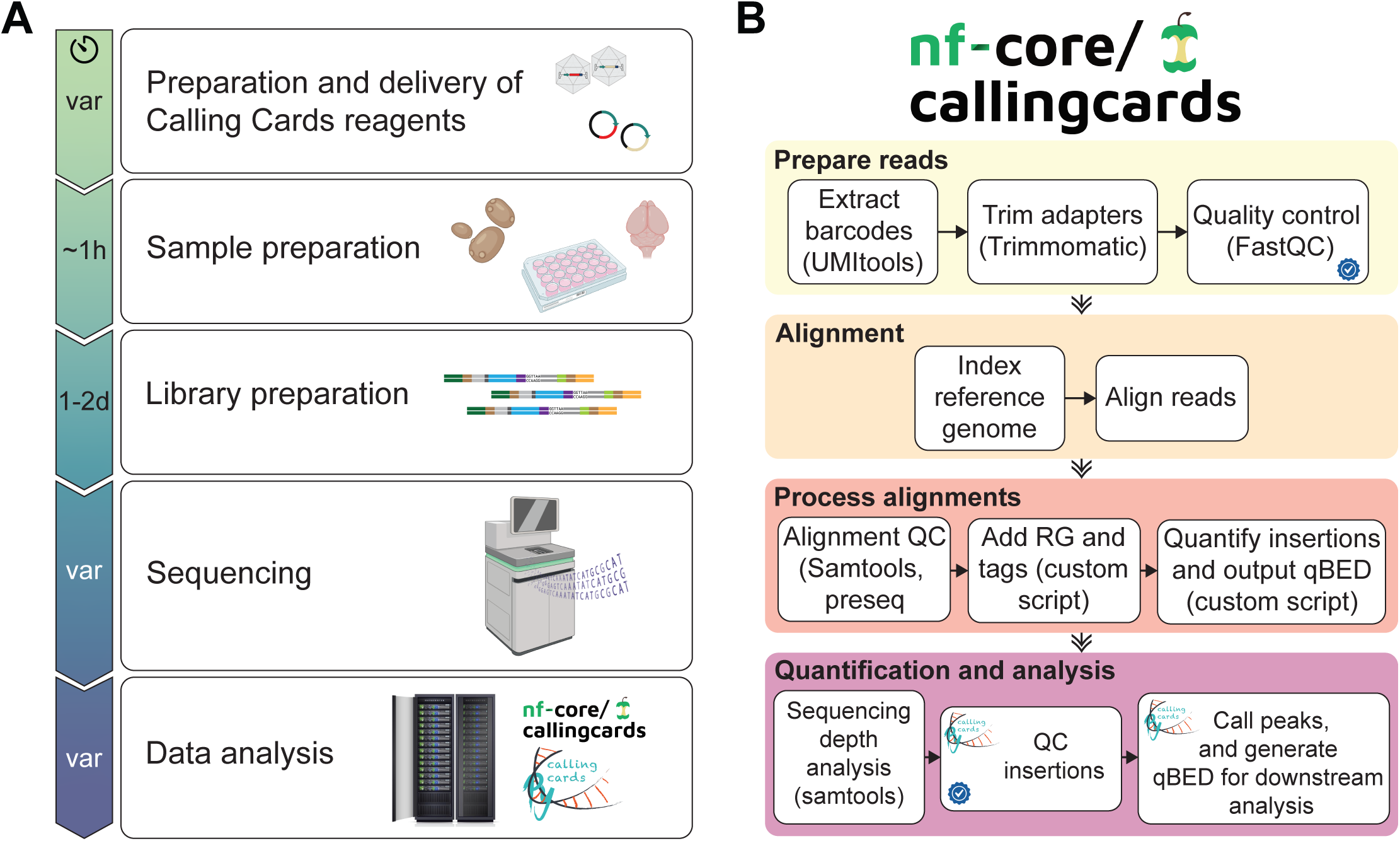
General workflows of a Calling Cards experiment. **(A)** The wet lab protocol is split into five main stages: 1) the viral or plasmid Calling Cards reagents are prepared and delivered into the target cells/tissue; 2) The sample is harvested; 3) the sequencing libraries are prepared; 4) the libraries are sequenced on Illumina NGS platforms; and 5) the generated FASTQ files are processed through the Calling Cards Nextflow pipeline and other downstream computational softwares. **(B)** The computational pipeline is distributed as a self-contained package that will process FASTQ files to Calling Card qBED files. The pipeline is divided into four main chunks: 1) the reads are prepared by extracting sample barcodes, trimming Illumina adapters, and standard quality control; 2) the reads are aligned to a reference genome; 3) the alignments undergo standard quality control and sample barcodes are added to headers of each read then collated into a qBED file; 4) the output files can be used for downstream analysis such as differential peak analysis and motif enrichment analysis. The blue check mark represents steps where QC metrics will be written to a file in the output and analysis directory.

**Figure 3:**
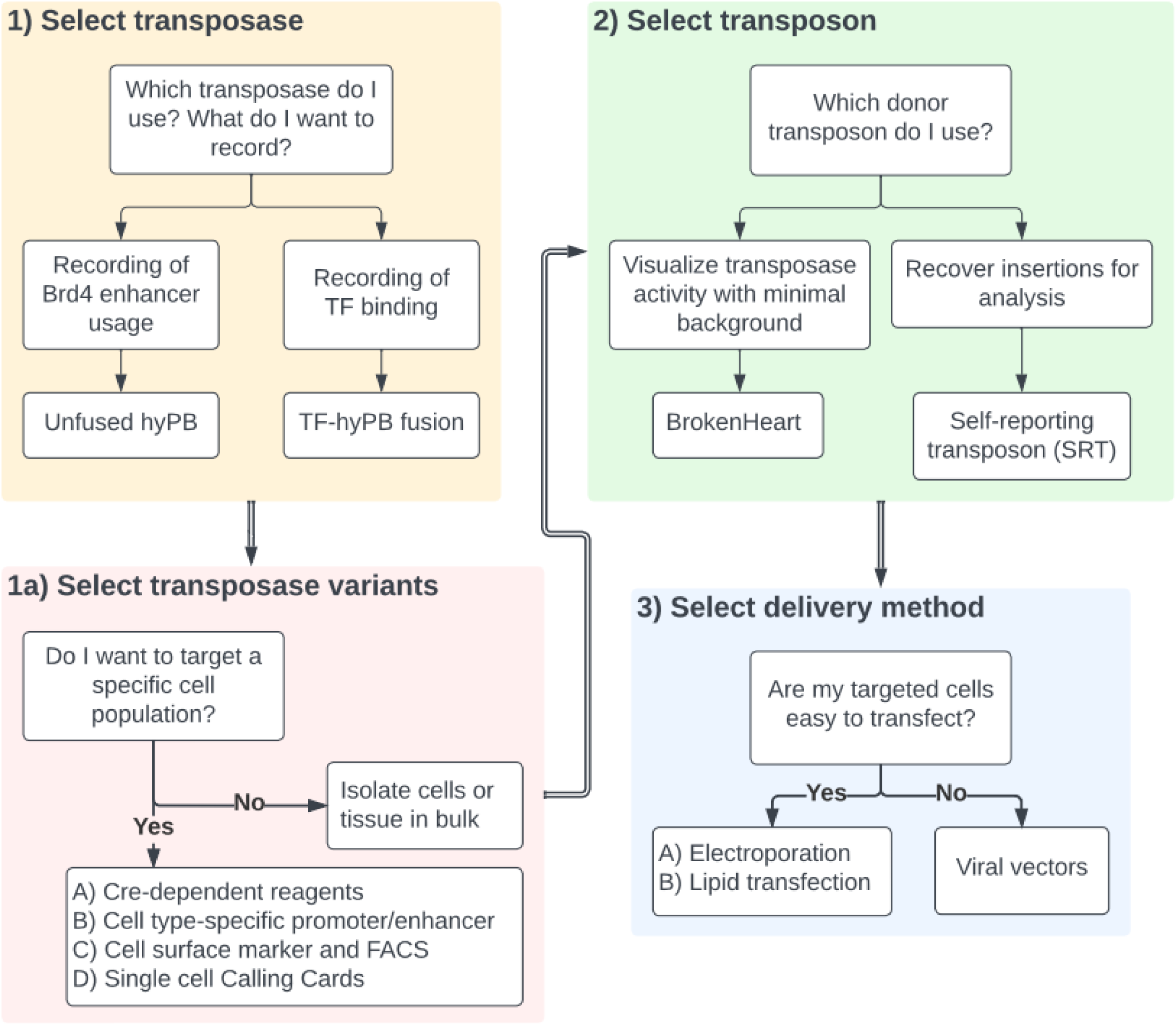
Decision tree for selecting Calling Card reagents for desired readouts. There are various transposase and donor transposon variants depending on the biological question and goal. The first decision is to decide between using an unfused or TF-fused transposase (step 1). If Calling Card recording is desired in a genetically defined cell population, “FrontFlip-hyPB” Cre-dependent transposase options are available (step 1a.A). Alternatively, a cell type-specific promoter can be used to drive expression of hyPB (e.g. Nestin-hyPB to target neural progenitors) (step 1a.B). A constitutive hyPB can be used for ubiquitous expression followed by enrichment of target cell population by FACS (step 1a.C). The final option is to conduct a single cell Calling Cards experiment (step 1a.D; see (Moudgil et al., 2020b) for details). The decision of donor transposon is made in step 2, followed by delivery method in step 3.

An illustrative vignette demonstrating how Calling Cards can be used to identify cumulative mouse brain region-specific BRD4-bound enhancer usage across postnatal neurodevelopment and maturation is bundled with this protocol. This same data is provided to the users for benchmark usage when setting up and testing the computational pipeline.

Finally, this protocol does not include the molecular steps and computational analysis for single-cell (e.g, 10x Genomics) Calling Cards, as these are distinct enough to require separate protocols (Moudgil et al., 2020b). However, the reagents and delivery steps are the same as described here.

## STRATEGIC PLANNING

### Design of custom TF-hyPB fusion proteins *(optional)*

Calling Cards relies on the presence of two components within a cell: the piggyBac transposase and the donor transposon. Using Calling Cards with unfused wild-type hyPB can identify BRD4-bound super enhancers. By creating a TF-hyPB fusion, one can redirect the insertion of Calling Cards to specific TF binding sites. We have previously used this for several TFs in yeast (e.g., Gal4, Gal80, Ste12, Bas1, Pho2, Gcn4, and Pho4) (Wang et al., 2007) and vertebrate systems *in vitro (*e.g., SP1, FOXA2, BAP1, ASCL1, MYOD1, NEUROD2, and NGN1) (Moudgil et al., 2020b; Cammack et al., 2020; Lalli et al., 2022; Yen et al., 2018) and *in vivo* (e.g., SP1) (Cammack et al., 2020). The general workflow to design, create, and validate a TF-hyPB fusion is described in **Figure 4**.

**Figure 4.**
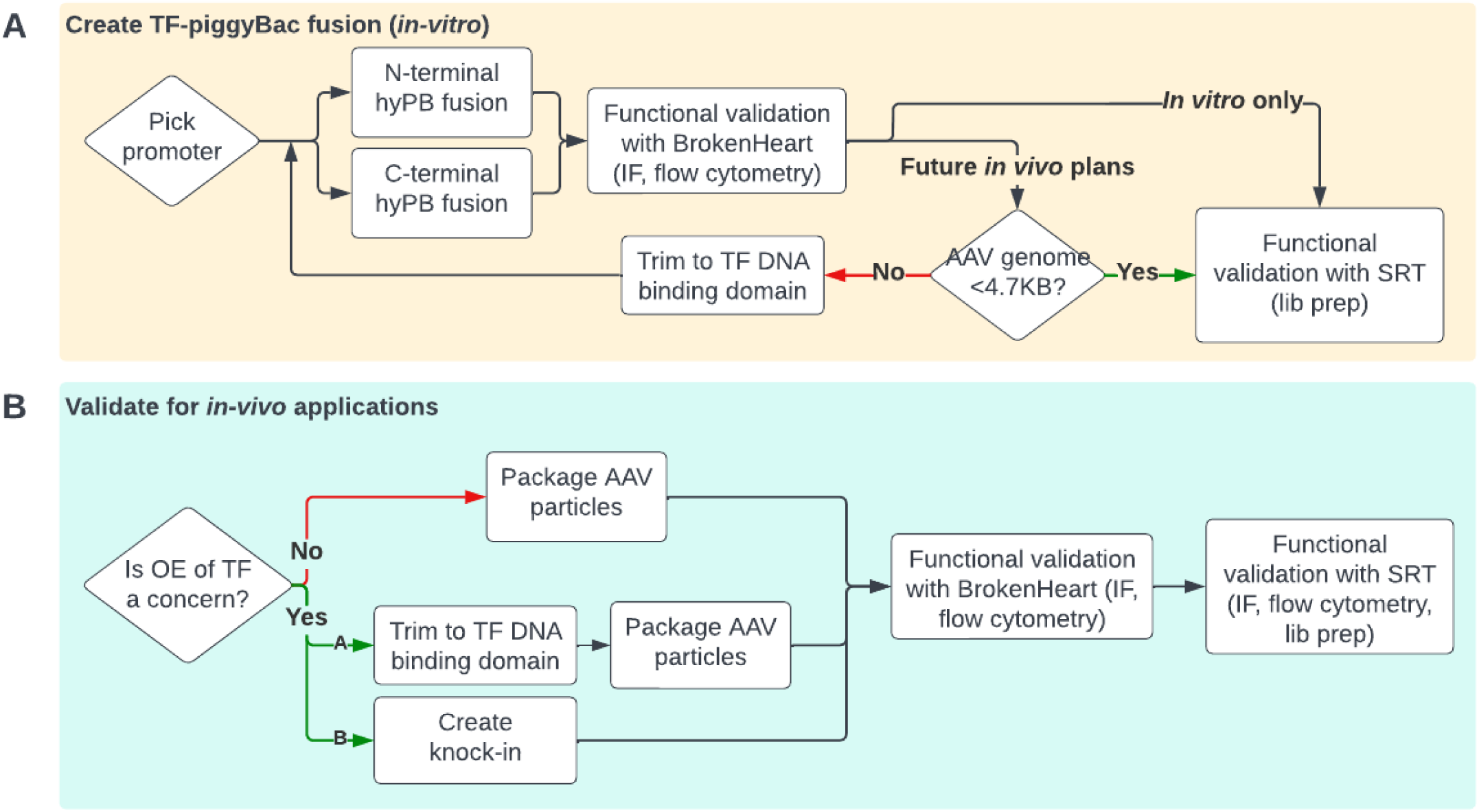
General workflows for creating TF-piggyBac fusions. **(A)** Steps to create a TF-hyPB fusion construct. Functional validation with immunofluorescence or flow cytometry is recommended to be performed using the BrokenHeart donor transposon due to its complete absence of fluorescence background (Supplemental Figure 2A), compared to the minimal background of the SRT. Final functional validation is performed using the SRT to generate libraries for downstream analysis. **(B)** Additional considerations for in vivo applications.

The first step is to select a suitable promoter for your targeted application. Ubiquitous promoters such as EF1a (Wang et al., 2017), CMV early enhancer-chicken ꞵ-actin (CAG) (Alexopoulou et al., 2008), and phosphoglycerate kinase 1 (PGK) (McBurney et al., 1991) have been successfully used to drive expression of the transposase. Next, clone both an N- and C-terminal TF-hyPB fusion construct. As TFs have diverse binding modes, having both versions will allow empirical determination of which fusion has the most efficient transposase activity, yet maintains specificity for the TF-targeted motif. With the completed expression construct, it is important to sequence the entire plasmid to ensure that all elements are intact, especially the repetitive AAV ITRs which are prone to deletions. Low cost, commercial long read sequencing (e.g. Plasmidsaurus) can be a useful resource to sequence the entire plasmid without primer walking, to determine if multiple plasmids species are within the submitted sample, and to resolve repetitive regions such as AAV ITRs that are often difficult for traditional Sanger sequencing.

Once the sequence is confirmed, the TF-hyPB fusion can be functionally validated by transfection with BrokenHeart *in vitro* into an easily transfectable cell line such as HEK293 or mouse neuroblastoma N2A cells. BrokenHeart is a tdTomato reporter that is interrupted with a donor transposon (**Supplemental Figure 1A**). Upon transposase activity and removal of transposon, the coding sequence is rescued and tdTomato will be expressed (**Supplemental Figure 1B**). By co-transfecting TF-hyPB fusion and Brokenheart, one would expect tdTomato expression in the cells only if the TF-hyPB fusion is functional. N- and C-terminal fusions can be quantitatively compared by microscopy or FACS to determine the relative activity compared to an unfused hyPB. In general, we see that fusions to any TF reduce transposase activity, yet this reduced activity is still sufficient to mediate transposition. This suggests that transposon levels, rather than transposase activity, is typically the rate limiting process (Wu et al., 2006; Nakazawa et al., 2009), and this is consistent with our observations of various donor/transposon ratios *in vitro* (**Supplemental Figure 2**)

If future *in vivo* experiments with AAVs are desired, it will be important to ensure that the length of the AAV transfer genome (ITR to ITR) is less than ∼4.7kB, the maximum cargo size of AAV particles for efficient viral packaging (Wu et al., 2010). If the sequence is larger, steps to trim away bases that are not critical for TF DNA binding will be necessary and functional validation should be redone to confirm TF-hyPB fusion activity. Lentiviral vectors with larger packaging capacities have also been used to deliver TF-hyPB fusions *in vivo* with limited success, likely due to the more restricted spread of lentivirus in the brain. Finally, after BrokenHeart validation, functional validation recovering insertion sites with SRTs will reveal if TF-hyPB fusion directs Calling Card insertions at expected TF binding sites to confirm that fusion of hyPB does not alter the specificity or TF binding properties. If available, TF ChIP-seq data from the same cell lines can provide a benchmark to validate that the TF-hyPB fusion is functioning as expected. If not, detection of the TF’s canonical motif using tools such as HOMER (Heinz et al., 2010) or the MEME suite (Bailey et al., 2015) can also indicate on-target activity. Likewise, comparison to profiles from unfused hyPB can confirm redirection of binding. After the construct passes all QC and functional validation steps described above, then it can be packaged into AAV particles and injected *in vivo* into the target tissue or experimental system of choice.

A potential concern when expressing the Calling Card reagents into cells through transfection or transduction is that the TF-hyPB fusion protein, and inadvertently the TF itself, is expressed above endogenous levels, and could thus alter gene expression. If overexpression is a concern, there are multiple approaches that can be used. The first (**Figure 4B**), is to trim to the minimal TF DNA binding domain, as mentioned for reducing size. Removal of other effector domains may render the TF-hyPB fusion sufficient to bind DNA but not interact with co-factors, thus minimizing effects of overexpression. Another approach is to create a knock-in hyPB as a fusion to the endogenous TF locus of a cell or mouse line. For all approaches, functional validation with BrokenHeart and SRT should be carried out at an appropriate time point, allowing enough time for the AAV to mature and Calling Card reagents to express at high levels. In brain tissue, AAVs are often allowed to mature 10+ days, although we have been able to recover sufficient insertions as early as 2 days after injection *in vivo* (**Figure 5A-C**). AAV maturation can be tracked over time with tdTomato RT-qPCR, which correlates with insertion density (**Figure 5C-E).**

**Figure 5.**
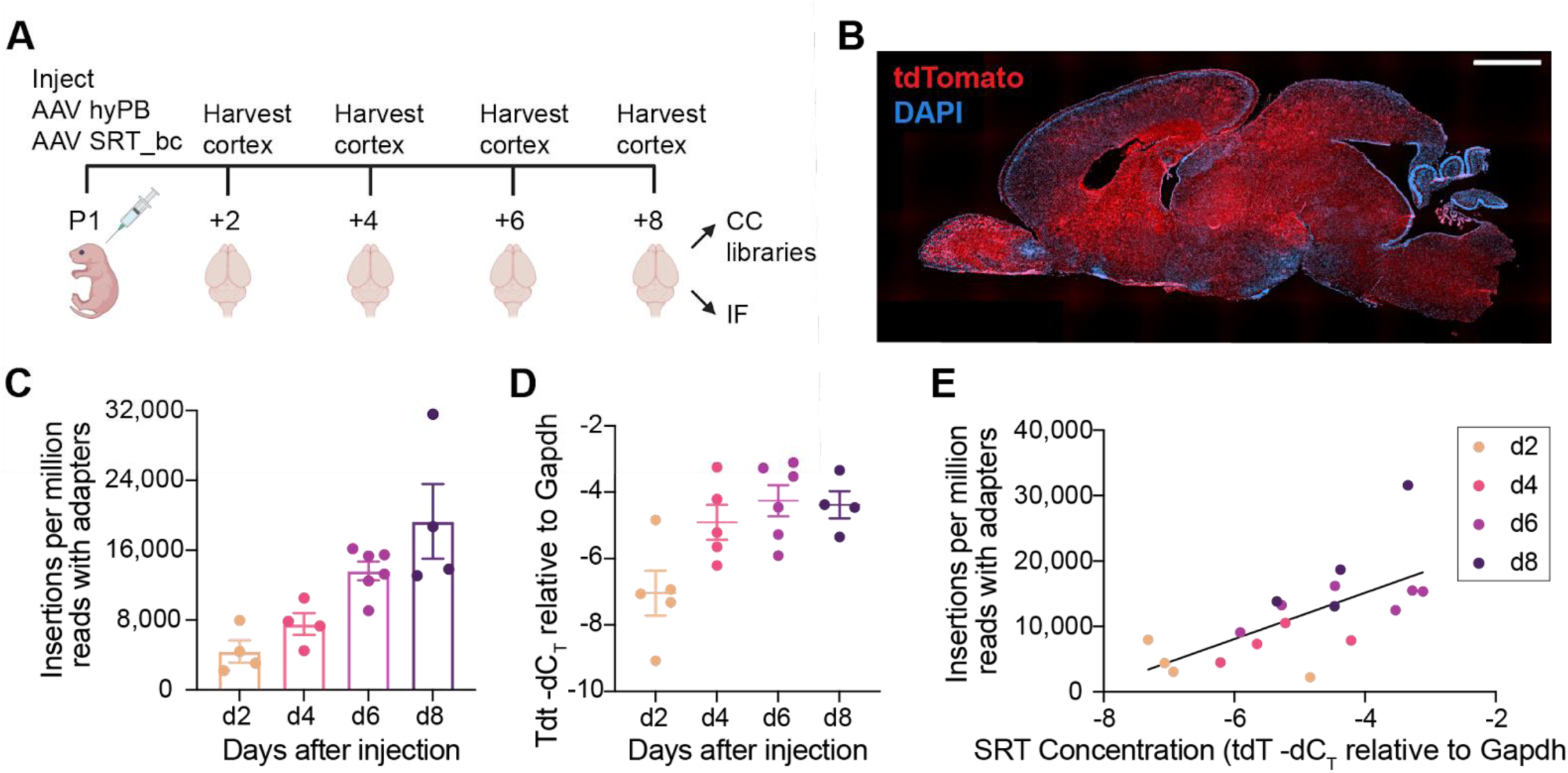
Timeline of Calling Cards activity in the mouse brain after AAV delivery. **(A)** Schematic of AAV Calling Cards time course experimental design. **(B)** Sagittal section of a brain harvested 2 days after neonatal intracerebroventricular injection with Calling Cards reagents, showing widespread expression of SRT-derived tdTomato. Scale bar 1000 μM. **(C)** Insertion counts recovered at each time point, normalized to read depth. n = 4-6 hemi-cortices. **(D)** SRT concentration, measured by RT-qPCR as tdTomato -dCT relative to Gapdh, over time. **(E)** Insertion counts recovered as a function of SRT concentration. Simple linear regression, R^2^ = 0.4447, p = 0.0025.

### Considerations for primer selection and ordering for sequencing libraries

In the final steps of library preparation, a dual indexing strategy is used to multiplex and increase the number of samples that can be sequenced per run to decrease costs. Here, we refer to “indexes” as sequences within Illumina adapters that are sequenced independently of the standard R1/R2 sequences, while “barcodes” are within the R1/R2 reads. In ideal conditions, unique dual indexes would be used during the indexing PCR to mitigate index hopping, which occurs when an index that is reused across multiple samples within the pool is incorrectly assigned to sample reads within a run (Kircher et al., 2012). This typically occurs when free adapters from the library preparation steps are carried over into the multiplexing step and hybridize to the wrong i5 or i7 index sequence. However, since the OM-PB primer (see **Supplemental Figure 3**), which contains the Illumina P5 adapter, Index2/i5, TruSeq Read1, 3bp primer barcode, and partial piggyBac LTR is ∼100bp long, it is cost prohibitive to synthesize an OM-PB primer with unique Index2/i5s for each sample. Instead, we have found it most effective to assign unique Index1/i7 indexes to each sample and have a small collection of up to 8 OM-PB primers with the same Index2/i5 yet varying the 3bp primer barcode with a hamming distance of 2 (GCA, ATC, CTA, ACG, CGT, TGC, GAT, and TAG). If more OM-PB primers are needed, additional combinations can be created by switching the Index2/i5. Note that these indexes and primer barcodes are used to distinguish samples on the sequencer. These are separate entities from SRT barcodes, which are used to identify unique insertions in the same locus *within* each sample.

While there are many strategies to allocate indexes and barcodes, one potential approach is to designate an Index2/i5 to an experiment, primer barcodes to different animals within that experiment (biological replicates), and Index1/i7 as different tissue samples or technical replicates for each animal, if multiple samples are prepared.

We have found Calling Cards to be reproducible across biological replicates as shown by the high correlation of normalized insertions per million (**Supplemental Figure 4**). Thus, for a given experiment, we typically create at least three replicate libraries per animal, with at least 3 animals per experimental group. When sequenced to saturation, this is typically sufficient to collectively recover at least 500k unique insertions per experimental group, which is a threshold we have found to be reliable for peak calling. The additional benefit of uniquely indexed samples is that deeper sequencing of specific samples can be re-pooled and sequenced without worry of index/barcode clashing.

### Expertise needed to implement the protocol

Basic molecular biology skills are required to successfully perform this protocol. If performing *in vivo* experiments, basic animal handling and husbandry skills are also necessary. Also, the use of viral vectors requires proper safety training and laboratory approval according to institutional guidelines. The sequencing of libraries requires the use of Illumina high throughput sequencing instruments that are typically found within genomics core facilities or commercial fee-for-service sequencing companies. For data analysis, a Linux based high-performance computing environment or small dedicated server is necessary for computationally intensive tasks. If one is not available, pay-as-you-go cloud computing platforms such as Amazon Web Services (AWS), Google Cloud, or Microsoft Azure can be used. Proper setup of the computational environment benefits from familiarity with nextflow and container runtimes such as Charliecloud, Singularity, or Docker. Basic to moderate skills with computational and bioinformatic analysis using command line tools, packages, and job schedulers is required. If not available within the lab, this computational expertise, similar to what would be required for RNA-seq, ChIP-seq or ATAC-seq workflows, may be available through local genomics cores that provide sequencing services.

### Limitations

While Calling Cards enables the recording of protein-DNA interactions over time, the readout of the method provides a cumulative history of enhancer usage or TF binding and is unable to resolve the temporal order of insertions (e.g. which insertions occurred first vs. those that occurred at the end of the recording period). To obtain some time information, one could harvest samples in a time course and resolve unique Calling Card insertion peaks by computationally subtracting common regions. This approach was used to map SP1 binding and expression of early and late genes in the developing mouse cortex (Cammack et al., 2020).

Another limitation is that current Calling Card reagents record continuously from delivery of the viral reagent (hyPB and/or SRT) until sacrifice of the animal. Recording only during specified time points would require development of drug inducible transposases, which would open up additional experimental opportunities.

The protocol described here is based on starting from a total RNA sample derived from potentially many cells (a “bulk” sample), thus the Calling Cards data represent the average signal from all the different cell types expressing the transposase. If the sample is heterogeneous like the brain, the resulting data should be interpreted taking this into account: the bulk Calling Cards data represents the average insertions across all the cell types expressing transposase. However, Calling Cards is compatible with droplet-based microfluidic platforms such as 10x Genomics to identify enhancer usage or TF binding with single cell/nucleus resolution. This innovation circumvents the need for SRT barcodes, as the cell-barcodes inherent to the single cell platforms can serve this purpose. The protocol adaptations for these single-cell Calling Cards are covered in (Moudgil et al., 2020b). Of note, the Nextera Mate Pair Sample Prep Kit (Illumina FC-132-1001) used for single cell Calling Cards library preps has been discontinued by the manufacturer. An in-house protocol is being developed and will be published when completed, though interested readers can reach out to us sooner. An alternate non-single cell approach to measure Calling Cards from specific cell types is to use Cre-dependent reagents (Cammack et al., 2020).

Another limitation, based on observations from single cell Calling Cards data, is that the number of insertions per cell is relatively low (<100). While this decreases the potential deleterious effects of transposon insertions in key regulatory elements, since any given cell has few insertions, the recovery of Calling Card insertions from rare cell types in sufficient numbers poses a challenge. The number of biological replicates needed to achieve a 500k unique insertion threshold may be a limitation and should be considered during experimental design.

The number of Calling Cards insertions is also dependent upon the delivery and expression of the SRT and piggyBac transposase. The copy number of SRTs is important in determining the success of a Calling Card experiment. When transfecting cells with plasmids, one can begin with a 1:1 plasmid cocktail of SRT:transposase and can further optimize by increasing the ratio and amount of SRT (**Supplemental Figure 2**). The same applies to the AAV Calling Cards reagents as sufficient time is needed for the AAV to mature, express the transgenes, and functionally hop into the genome. Our preliminary data demonstrates functional hopping as early as 2 days after injection (**Figure 5**), though insertions increase with time as expected.

The protocol presented here represents an optimized library preparation protocol (**Figure 6**) to enable robust recovery of Calling Cards insertions even when in lower abundance (e.g., from a relatively sparse cell type labeled by a Cre line), however, if extremely rare cell types are targeted using either a transgenic or molecular approach, FACS enrichment of TdTomato positive cells/nuclei prior to RNA extraction may be necessary to enrich for cells with Calling Card insertions.

**Figure 6.**
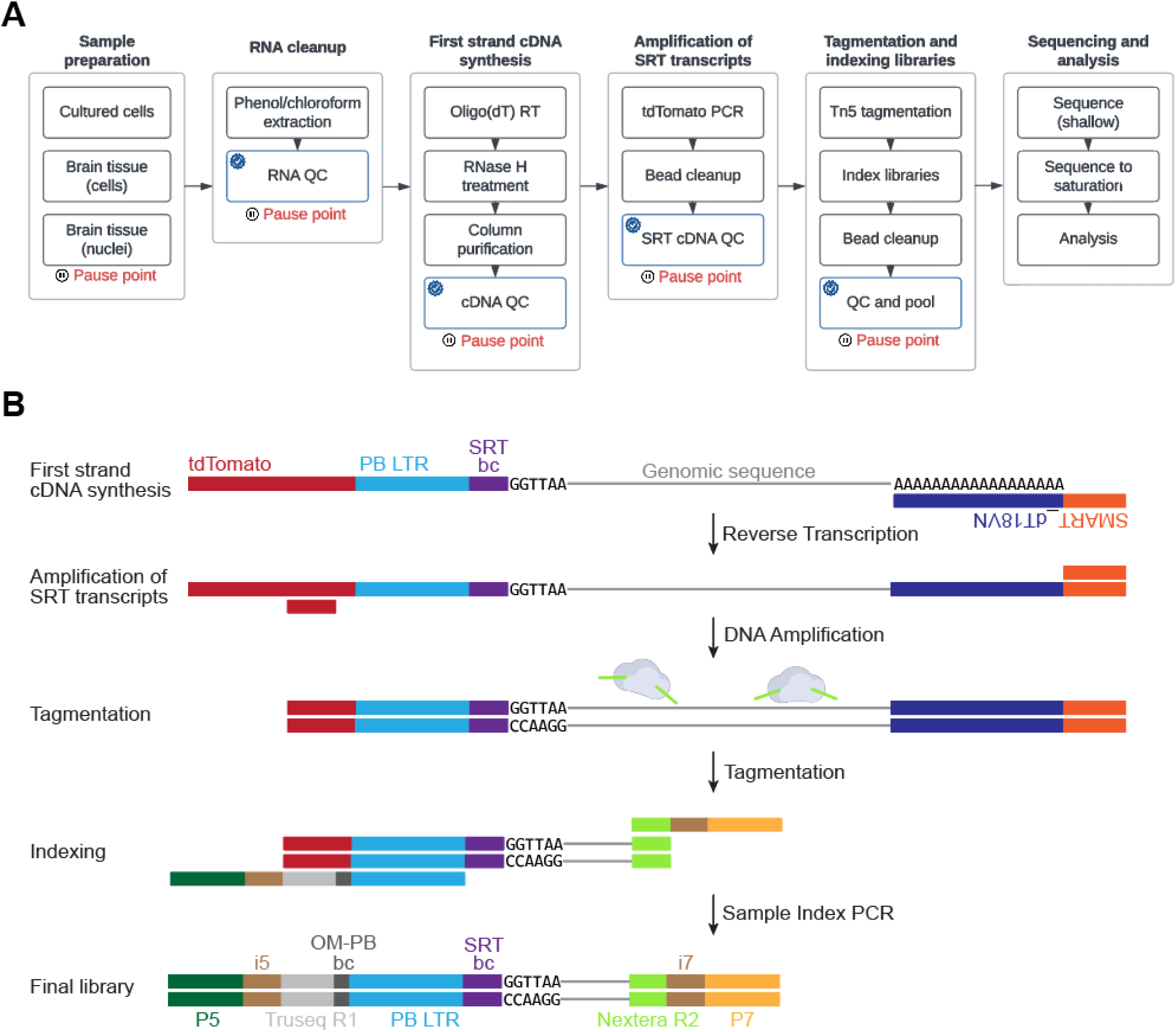
The experimental workflow for bulk Calling Card library preparation. **(A)** The sequencing library preparation protocol is broken down into several main sections. Recommended quality control (QC) checkpoints are noted by the blue checkmark. Appropriate pause points are shown in red. **(B)** A cartoon depicting how the libraries are prepared and the final library structure that is loaded onto the sequencer.

## BASIC PROTOCOL 1: PREPARATION AND DELIVERY OF CALLING CARDS REAGENTS

This two-track protocol describes the procedure to transfect plasmids into *in vitro* cultures or inject AAV reagents into the developing mouse brain. Prior to beginning this protocol, high quality endotoxin-free Calling Cards plasmids or purified AAVs are required. Plasmid DNA can be prepared using many available commercial maxiprep kits. If an academic core or commercial virus packaging service is not available to generate AAVs, the procedure is outlined and described here (Challis et al., 2019). This protocol describes neonatal intracerebroventricular injections as an example, but AAVs can be injected into the target tissue of choice.

*NOTE: All animal studies were approved by and were performed in accordance with the guidelines of the Animal Care and Use Committee of Washington University in Saint Louis, School of Medicine and conform to NIH guidelines of the care and use of laboratory animals.*

*NOTE: Viral vectors are biohazardous materials and investigators must be trained according to governmental and institutional regulations and standard operating procedures*.

### Materials

#### Plasmid preparation

Plasmids, supplied as bacterial stabs (Addgene; see **Table 1** for complete list
Lysogeny Broth, Miller (Beckman, Dickinson, and Company 244610)
Carbenicillin disodium salt (Sigma-Aldrich C1389)
Kanamycin sulfate (Sigma-Aldrich K1377)
NEB Stable Competent E. coli (High Efficiency) (NEB C3040H, or equivalent)
ZymoPURE II Plasmid Maxiprep Kit (Zymo D4203) or EndoFree Plasmid Maxi Kit (Qiagen 12362), or equivalent
Baffled Erlenmeyer flasks (Sigma-Aldrich CLS44441L-6EA, or equivalent)
Nalgene PPCO Centrifuge Bottles (ThermoFisher 3141-0250, or equivalent)
Sorvall LYNX 6000 Centrifuge (ThermoFisher 75006590, or equivalent)
Fiberlite F14-6 x 250y Fixed-Angle Rotor (ThermoFisher 096-062075, or equivalent)
Thermomixer R (Eppendorf 05-400-205, or equivalent)
MaxQ 8000 Incubated Shaker (ThermoFisher SHKE8000, or equivalent)
Precision Econotherm Incubator (ThermoFisher 51221126, or equivalent)
Vortex-Genie 2 Mixer, Variable speed (Scientific Industries SI0236, or equivalent)
Mini Microcentrifuge (Midsci MF12, or equivalent)
NanoDrop Spectrophotometer
Qubit 3.0 Fluorometer (ThermoFisher Q33216)
Endosafe nexgen-PTS (Charles River PTS150K)
Endosafe Compendial LAL Cartridges (Charles River PTS2001)

#### *In vitro* plasmid transfections

DMEM (ThermoFisher 11965092)
Fetal Bovine Serum - Premium (Atlanta Biologicals S11150, or equivalent)
Trypsin-EDTA (0.25%) (ThermoFisher 25200056)
Polyethylenimine hydrochloride MAX (MW 40,000) (Polysciences 24765)
OptiMEM (ThermoFisher 31985070)

#### Neonatal intracerebroventricular injections

Purified AAV particles
0.5% sodium hypochlorite (10% Clorox) solution
Gas Tight syringe with small removable needle, 50ul (Hamilton 80930)
Custom syringe needles (33G, 0.5in, point style 4, 12 deg bevel angle) (Hamilton 7803-15)
Custom syringe needle guard (**see Supplemental Figure 3**)
Laboratory support stand (Grainger 23YW89, or equivalent)
Multipurpose clamp (Grainger 404R44, or equivalent)
Heating pad (Sunbeam 731-500, or equivalent)
Ice bucket with lid (Thomas Scientific 20A00F928, or equivalent)
Beaker (Cole-Parmer EW-34502-42, or equivalent)

### Calling Cards plasmid preparation (timing: variable)

Once you receive the bacterial stabs from Addgene, it is recommended to create glycerol stocks for each of the plasmids for long-term storage at −80°C. A complete guide can be found on their website (https://www.addgene.org/protocols/create-glycerol-stock/).

**Table 1.**
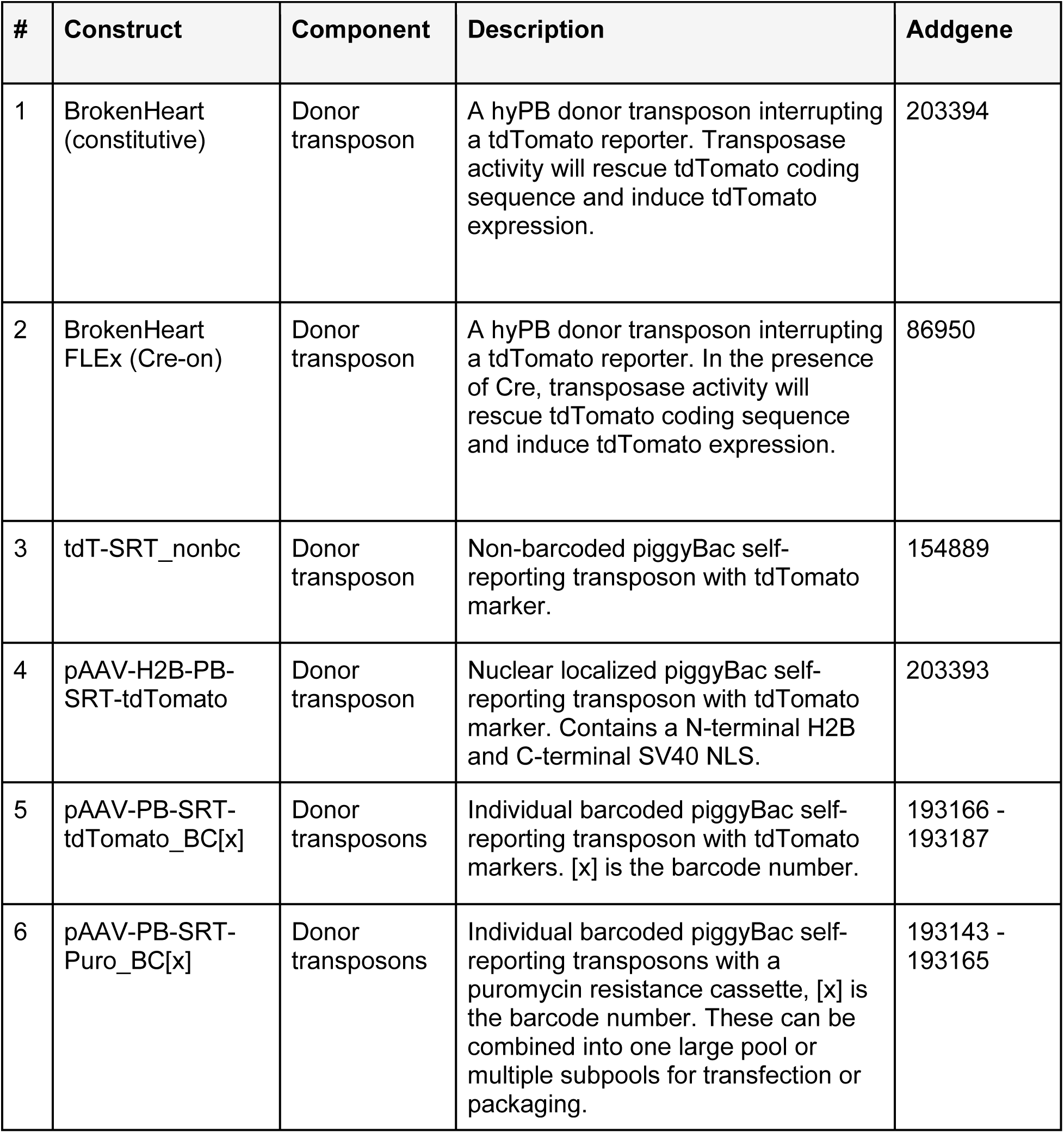
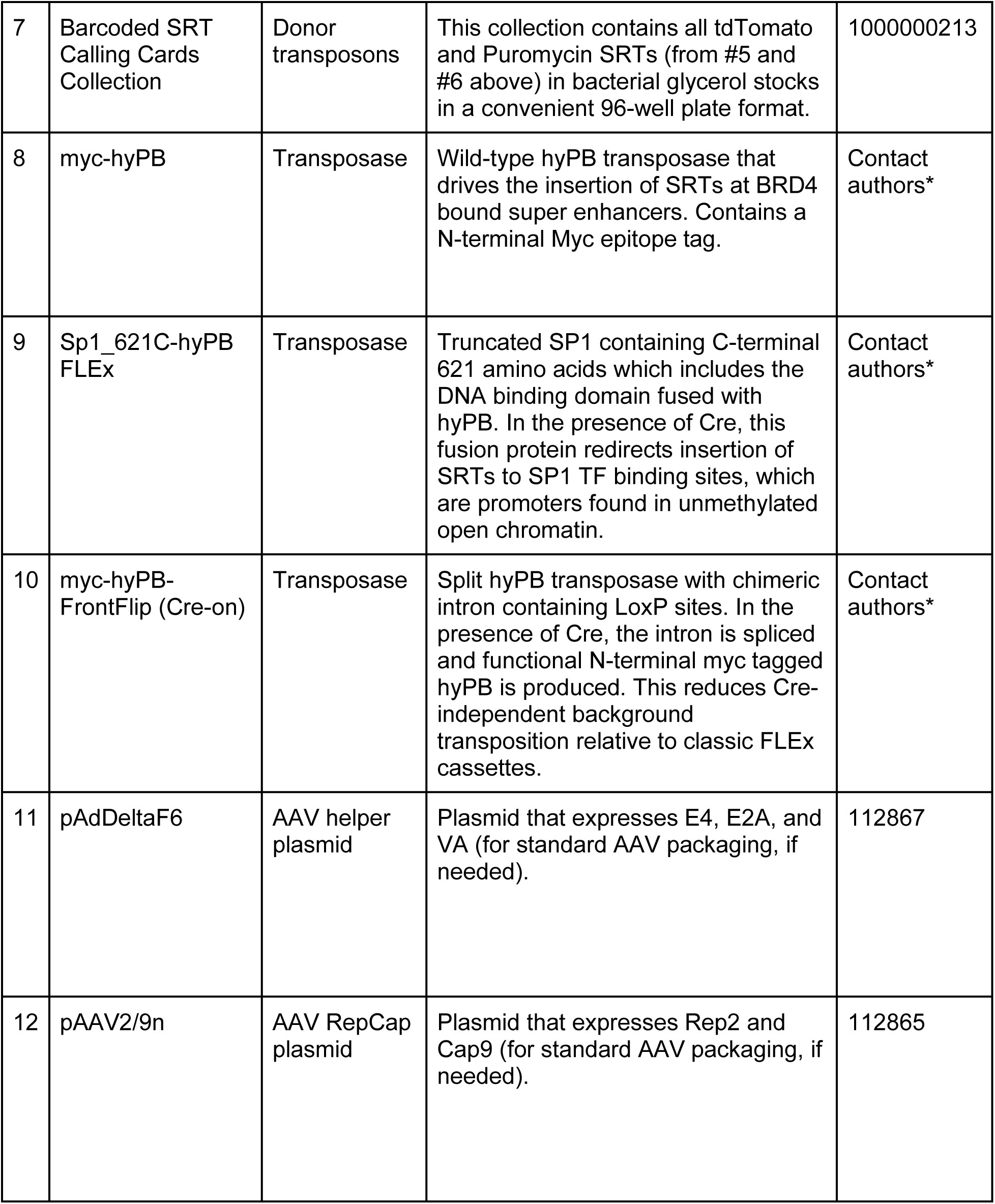
Calling Cards plasmids and AAV constructs *NOTE: piggyBac donor transposon plasmids and AAV packaging plasmids are distributed through Addgene’s website (https://www.addgene.org/). Contact Addgene or authors for information about access to plasmids.

*NOTE: it is highly recommended to validate each plasmid by long-read sequencing. Fully sequencing the entire plasmid is advantageous because it can reveal unexpected products, deletions, recombinations, and concatemers. Additionally, it can sequence through repetitive elements such as ITRs, which can form secondary structures and are typically troublesome for Sanger sequencing reactions. Confirmation of intact ITRs is crucial as mutations in this region can affect packaging efficiency.*

Next, we want to create a balanced plasmid pool of all the barcoded self-reporting transposons (plasmid collection 7 in Table 1).

1. Quantify the concentration of each of the purified plasmids by nanodrop or Qubit. *NOTE: the Qubit assay is recommended for sample concentrations that are <100ng/ul*.
2. Pool equal masses (e.g. 100ng) of each plasmid into a clean 1.5ml microcentrifuge tube.
3. Pulse vortex to mix and briefly centrifuge.
4. Quantify the concentration of the pool by nanodrop or Qubit.
5. Transform plasmid pool into NEB Stable Competent *E. coli* cells or equivalent bacterial strain according to manufacturer’s instructions. *NOTE: use of strains that are optimized for plasmids containing repeat elements with reduced recombination and endonuclease activity are recommended.*
6. After 1 hr of outgrowth, directly inoculate a liquid overnight culture and incubate in a shaking incubator for 12-18 hr at 37°C or 24 hr at 30°C. *NOTE: the standard step of plating on selection plates is skipped to streamline amplification of the plasmid pool and recovery of all barcoded elements. The outgrowth can be plated; however, the whole plate would need to be scraped then inoculated into a liquid overnight culture.*
7. The next morning, harvest bacteria and purify plasmid DNA. *NOTE: many commercial kits offer endotoxin-removal steps. It is highly recommended to perform this step*.
8. Quantify concentration by nanodrop or Qubit.
9. Sequence to verify important plasmid features (e.g, AAV ITRs, tdTomato transgene, and PB LTR) by Sanger sequencing or the entire plasmid by long-read sequencing.

This is a safe stopping point and the plasmid pool can be stored at −20°C, or proceed to the next step.

10. (Optional). Assess endotoxin levels to ensure virtually undetectable levels of <2.5 endotoxin units [EU]/ml. *NOTE: depending on the injection route and tissue, trace amounts of endotoxin can trigger strong adaptive immune responses and lead to inflammatory response, especially when injected intravenously.*

It is highly recommended to assess barcode distribution of the plasmid pool by next-generation sequencing prior to AAV packaging. This procedure is described in Support Protocol 1.

### Delivery of Calling Card reagents *in vitro* (timing: variable)

There are numerous methods and commercial reagents available to transfect plasmids into cells. Polyethyleneimine (PEI) is a robust and cost-effective reagent that has been optimized for high transgene expression in a variety of cell lines. Refer to this document for a generalized transfection protocol (https://currentprotocols.onlinelibrary.wiley.com/doi/10.1002/cpch.25).

### Intracerebroventricular injection (timing: 1 hr)

All animal studies were approved by and were performed in accordance with the guidelines of the Animal Care and Use Committee of Washington University in Saint Louis, School of Medicine and conform to NIH guidelines of the care and use of laboratory animals.

Delivery of AAVs into the ventricle of P0-2 mouse pups is an easy and efficient route to label cells in the developing cortex (Cammack et al., 2020). A key benefit of delivery at this age is that it allows for efficient CNS labeling yet is considered a non-surgical injection since the skull bone has not yet hardened. Additionally, the procedure for a whole litter is more rapid than adult stereotactic injection and the survival rate is very high. Investigators and end users should be trained according to institutional guidelines prior to using viral vectors in the laboratory.

See **Supplemental Figure 5** for an example of the setup.

*NOTE: if available in the animal facility, work with viral vectors and animals in a biosafety cabinet.*

11. Remove dam and stud from home cage and place in clean cage away from pups to avoid stressing the dam.
12. Set a heating pad to “low” (warm to the touch) and place the home cage on top.
13. Prepare an appropriate amount of AAV cocktail by thawing viral aliquots on ice (6ul per animal). Dilute each AAV with sterile PBS to 1.0×10^13^ vg/ml and mix the transposase and donor transposon AAVs 1:1. Keep the AAV cocktail on ice. (*Optional):* Add 0.05% Fast Green FCF dye to the AAV cocktail to facilitate visualization of the injection into the ventricles.
14. Fill a beaker with freshly made 0.5% sodium hypochlorite (10% Clorox) solution to decontaminate any material or liquids that come in contact with AAVs.
15. Prepare and clean a Hamilton syringe with 5 full volume washes with sterile deionized water, 5 washes with 80% ethanol, followed by 5 washes with sterile deionized water.
16. Load the AAV cocktail into a Hamilton syringe. Avoid excessive pipetting and handling to prevent bubbles as injection of air into the ventricle is often fatal.
17. Place a wet paper towel on a bed of ice. Anesthetize pups by placing them on the ice for 5-8 mins (maximum of 15 mins) to induce hypothermia. The paper towel ensures that their skin is not making direct contact with the ice. When unresponsive to physical stimuli, the pups are sufficiently anesthetized.
18. Working quickly but carefully, inject 1ul of virus per site at 3 sites (noted as x’s in the figure to the right) per hemisphere at a rate of 1ul per second. After each injection, wait 5 seconds before withdrawing the needle to minimize backflow. Repeat for the opposite hemisphere. (See **Supplemental Figure 5C** for details)
19. Place pup in home cage on heating pad for recovery.
20. Repeat injection steps 18 and 19 for the remaining pups.
21. Leave pups on the heating pad for 5-10 mins until they have returned to normal body temperature, indicated by movement, and return to pink color.
22. Carefully return pups to home cage. *NOTE: rub gloves through dirty home cage bedding prior to handling the pups to mask the foreign scent of nitrile gloves. This should decrease the chance of cannibalization.*
23. Dispose of any leftover virus or unwanted contaminated material that came into contact with AAV into the beaker with bleach. After 5 minutes, decontaminated unwanted materials can be disposed of properly.
24. Depending on experimental design, allow time for animals to age and AAVs to express. Collection can occur as early as 2 days, or as late as adulthood.

## BASIC PROTOCOL 2: SAMPLE PREPARATION

### Materials

TRIzol Reagent (ThermoFisher 15596026)
Chloroform (Sigma-Aldrich C2432)
RNA kit
TURBO DNase
High Sensitivity RNA ScreenTape (Agilent 5067-5579) and Sample Buffer (Agilent 5067-5580)
Qubit RNA Assay Kits (ThermoFisher Q32852, Q10211, or Q33224)
Refrigerated centrifuge (Eppendorf 5430R)
Rotor for 1.5ml and 2ml tubes (Eppendorf FA-45-30-11)
Nanodrop Spectrophotometer (ThermoFisher ND-2000)
Qubit 3.0 Fluorometer (ThermoFisher Q33216)
Tapestation 4200 System (Agilent G2991BA) (often available in genomics cores)

#### Harvesting *in vitro* cultures

There are numerous methods and commercial kits to purify total RNA. The steps outlined below describe a generic procedure to lyse cells and purify RNA using a phenol-chloroform extraction method. Specific cell lines or cultures may require some optimization or additional steps.

25. Carefully aspirate the growth medium.
26. Add 1ml cold TRIzol per 2.5×10^7^ cells directly onto cells and pipette up and down several times until cells have been lysed and homogenized. This can be scaled down or up as needed. *NOTE: washing cells with DBPS prior to addition of TRIzol can lead to mRNA degradation and is thus not recommended.*
27. Transfer to a clean microcentrifuge tube and incubate for 5 minutes at RT to allow for complete dissociation of nucleoprotein complexes. Samples can be frozen at −80C until further processing.
28. Add 0.2ml chloroform for every 1ml TRIzol used and shake vigorously for 30 sec.
29. Incubate for 7 mins at RT.
30. Centrifuge samples for 15 mins at 12,000 xg at 4°C.
31. Harvest the aqueous phase to a new tube and record the volume. *NOTE: Do not touch the interphase or organic phases when removing the aqueous phase! The interphase and organic phases can be saved and frozen at −80°C if desired for DNA and protein recovery.*
32. Proceed to RNA purification.

#### Harvesting brain tissue (timing: 20 min/mouse)

33. Deeply anesthetize a mouse with isoflurane.
34. Optional. Conduct a trans-cardial perfusion using ice-cold DPBS.
35. Harvest the brain and dissect specific regions if needed.
36. Transfer to a clean RNase-free tube. *NOTE: the tissue can be snap-frozen with liquid nitrogen and stored at −80°C until further processing with minimal impact on the quality of final Calling Card libraries.*

The steps outlined below describe a general procedure to homogenize mouse brain tissue and purify total RNA using a phenol chloroform extraction method. Any RNA cleanup approach that produces high quality DNA-free total RNA should be compatible with Calling Cards. Different tissue types may require some optimization.

37. Fresh or frozen tissue can be directly homogenized in 1ml TRIzol for each 100mg tissue.
38. Incubate for 5 mins at RT. If processing multiple samples, homogenize all samples and start the timer after the last homogenization.
39. Add 0.2ml chloroform for every 1ml TRIzol used and shake vigorously for 30 sec.
40. Incubate for 7 mins at RT.
41. Centrifuge samples for 10 mins at 12,000 xg at 4°C.
42. Harvest the aqueous phase to a new tube and record the volume. *NOTE: Do not touch the interphase or organic phases when removing the aqueous phase! The interphase and organic phases can be saved and frozen at −80°C if desired*.
43. Proceed to RNA purification.

## BASIC PROTOCOL 3: SEQUENCING LIBRARY PREPARATION

### Materials

Standard desalted primers (custom synthesized by Integrated DNA Technologies (IDT); sequences are provided in **Table 2**).
IDTE pH 8.0, 0.2um filtered (Integrated DNA Technologies 11-05-01-13, or equivalent) RNaseZap (ThermoFisher AM9782, or equivalent) RNA Clean & Concentrator-5 (Zymo R1014), RNEasy Plus Mini Kit (Qiagen 74134), or equivalent *NOTE: both RNA purification kits have been tested and validated for RNA purification. Other cell or tissue types may require alternative RNA purification reagents/kits.*
High Sensitivity RNA ScreenTape (Agilent 5067-5579) and Sample Buffer (Agilent 5067-5580)
Molecular Biology Grade Water (Corning 46-000-CM, or equivalent)
DEPC-treated water (Midsci IB42200, or equivalent)
Maxima H Minus Reverse Transcriptase (ThermoFisher EP0752)
dNTP mix (10mM ea, PCR grade) (ThermoFisher 18427088)
RNaseOUT Recombinant Ribonuclease Inhibitor (ThermoFisher 10777019) or SUPERase-In RNase Inhibitor (20U/ul) (ThermoFisher AM2696). *NOTE: both RNase inhibitors have been tested and validated.*
RNase H (New England Biolabs M0297L or ThermoFisher 18021071)
QIAquick PCR Purification Kit (Qiagen 28106)
Buffer EB (Qiagen 19086)
PowerUp SYBR Green Master Mix (ThermoFisher A25743, or equivalent)
Qubit ssDNA Assay Kit (ThermoFisher Q10212)
KAPA HiFi HotStart ReadyMix (Roche KK2601)
AMPure XP Reagent (Beckman Coulter A63882) or Mag-Bind TotalPure NGS beads (Omega Biotek M1378-02) *NOTE: both magnetic beads have been tested and validated for clean-up and size selection of Calling Card libraries. If using the Mag-Bind beads, in-house validation of size selection ratios of each lot with a DNA ladder is recommended.*
Ethyl alcohol 200 Proof (ACS/USP Grade) (Pharmco-Aaper 11100020S, or equivalent)
Qubit dsDNA High Sensitivity Assay Kit (ThermoFisher Q32851)
High Sensitivity D5000 ScreenTape (Agilent 5067-5592)
High Sensitivity D5000 Ladder and Sample Buffer (Agilent 5067-5593)
Nextera XT DNA Library Preparation Kit (Illumina FC-131-1096)
Cole-Parmer Essentials PTFE Tissue Grinder, 5ml Vessel (Cole-Parmer UX-44468-14)
Serrated Tip Plunger, 15ml, 270mm L (Cole-Parmer UX-44468-10)
96 well metal cooling block (Argos Technologies 63615-04), or equivalent
QuantStudio 6 Flex Real-Time PCR System (ThermoFisher 4485691), or equivalent system for SYBR-green qRT-PCR (optional - for library density QC)
T100 Thermal Cycler (Biorad 1861096), or equivalent
Magnetic Separation Rack for PCR strip tubes (Permagen MSR812, or equivalent)
Tapestation 4200 System (Agilent G2991BA) (often available in genomics cores)
Qubit 3.0 Fluorometer (ThermoFisher Q33216)
Nanodrop Spectrophotometer (ThermoFisher ND-2000)

#### RNA purification (timing: 60 mins)

The Zymo RNA Clean & Concentrator-5 Kit and QIAGEN RNEasy Plus Mini Kit have been tested and validated for brain tissue and *in vitro* cultured cells. Generally, any RNA cleanup and purification kit that yields high quality RNA can be used. Optimizations may be needed for other tissue types or kits. The following section describes the protocol using the Zymo RNA Clean & Concentrator-5 kit.

**Table 2.**
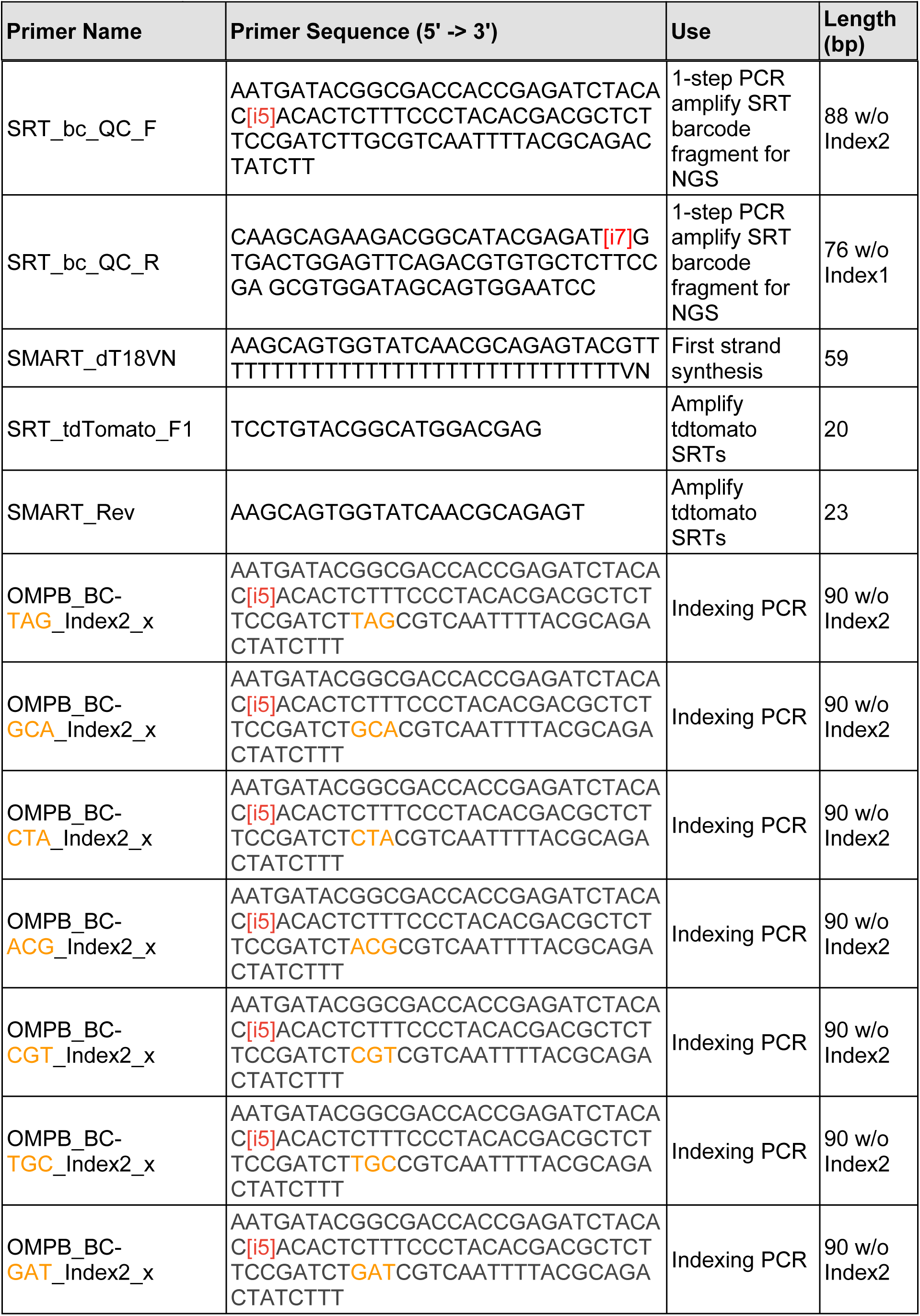
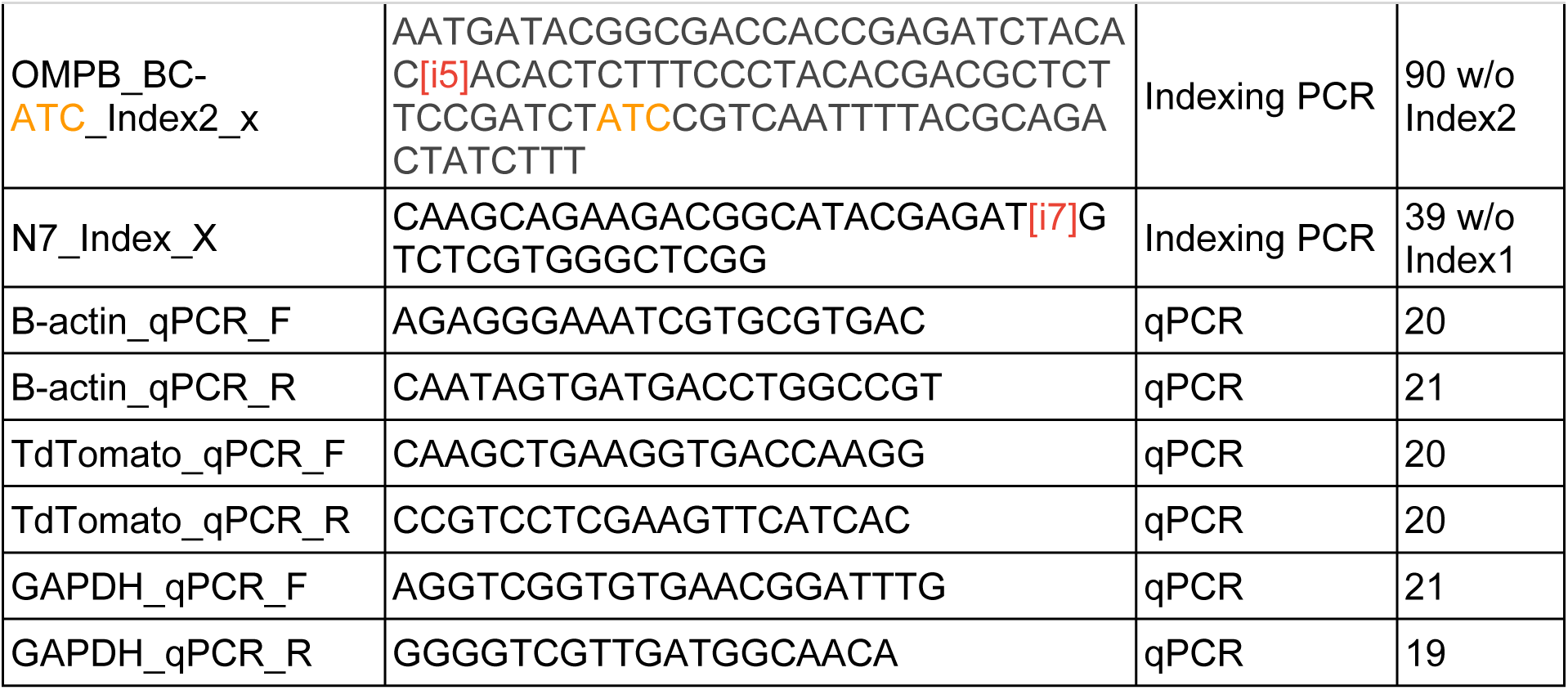
Primer sequences

*NOTE: Clean all working surfaces and pipettes with RNaseZap before setting up the workspace. Use sterile, RNase- and DNase-free, and filter barrier tips throughout the procedure.*

*NOTE: An on-column DNase treatment is described below. If it this does not sufficiently remove genomic DNA, a more effective DNase treatment in solution can be performed using TURBO DNA-free DNase Treatment and Removal Reagents (ThermoFisher AM1907), or equivalent.*

44. Add 2 volumes of aqueous phase of Zymo RNA Binding Buffer to each tube. *NOTE: For example, if you collected 500ul aqueous phase, add 1ml Binding Buffer*.
45. Add 1 volume of aqueous phase+Binding Buffer of 100% ethanol to each tube. Mix thoroughly by pipetting. *NOTE: For example, if you have 1.5ml aqueous phase+Binding Buffer from the previous step, add 1.5ml ethanol*.
46. Transfer 700ul to a column at a time. If using a centrifuge, spin for 15 sec at 12,000 xg at RT to bind RNA. Discard flow-through and continue for the remainder of the solution. *NOTE: A RNase- and DNase-free vacuum manifold can also be used to process many samples in parallel*.
47. Add 400ul RNA Wash Buffer to the column and spin at 12,000 xg for 1 min at RT. Discard flow through and move to a new collection tube.
48. Prepare the DNase mixture.

49. Add 40ul DNase mix directly to the membrane of each column. Incubate for 15 min at RT.
50. Add 400ul RNA Prep Buffer and spin at 12,000 xg for 15 sec at RT. Discard flow-through.
51. Add 700ul RNA Wash Buffer and spin at 12,000 xg for 30 sec at RT. Discard flow-through.
52. Add 400ul RNA Wash Buffer and spin at 12,000 xg for 1 min at RT. Discard flow-through.
53. Spin empty column at 12,000 xg for 1 min at RT to remove residual Wash Buffer.
54. Transfer the column to a pre-labeled RNase-free tube. Add 30-100ul nuclease-free water directly to the silica bed. Incubate at RT for 1 min, then spin at 12,000 xg for 2 min. *NOTE: the amount of water will depend on tissue/cell type, expected RNA yield, and desired concentration*.
55. Use 1ul to measure A_260_/A_280_ and estimate concentration using a Nanodrop. *NOTE: Values should be near 1.95-2.00*.
56. Assess quality, integrity, and concentration of RNA using a High Sensitivity RNA ScreenTape or Bioanalyzer RNA 6000 Pico Assay. *NOTE: Alternatively, the Qubit RNA High Sensitivity Assay and Qubit RNA IQ Assay can be used*.

| Component<br>Add in order | 1 rxn (ul) | 8 rxn + 10% (ul) |
| --- | --- | --- |
| DNase Digestion Buffer | 35 | 283.5 |
| DNase I | 5 | 40.5 |
| *Total* | *40* | *324* |

This is a safe stopping point and the RNA can be stored at −80°C, or proceed to first strand cDNA synthesis.

#### First strand cDNA synthesis (timing: 2 hrs)

The goal of this section of the protocol is to reverse transcribe polyA mRNAs using an oligo(dT) primer that also adds on a common primer sequence (Picelli et al., 2014). After first strand synthesis, RNase H is used to degrade the RNA strand within RNA/DNA duplexes. The cDNA products are then column purified as carryover SMART_dT18VN primer was found to increase nonspecific PCR products in the subsequent steps.

Use a thermocycler with a heated lid set to 105°C for all incubations throughout this protocol.

57. Set up the following PCR program on a thermocycler and preheat to 65°C.

58. Add the following components into individual nuclease-free tubes on ice. A master mix containing RT primer and dNTPs can be made.

| Temperature | Time | Cycles |
| --- | --- | --- |
| 65°C | 5 min | 1 |
| 4°C | Hold | 1 |
| 50°C | 60 min | 1 |
| 85°C | 10 min | 1 |
| 4°C | Hold | 1 |

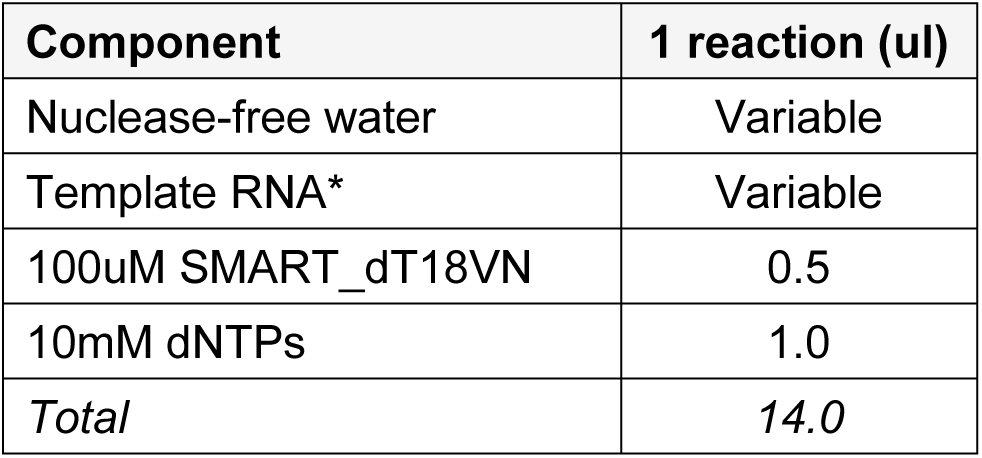 *Application note: Up to 5ug total RNA can be used for cDNA synthesis. It is recommended to maximize the starting RNA amounts to maximize recovery of Calling Cards insertions.
59. Mix by pipetting and centrifuge briefly.
60. Transfer tubes to preheated thermocycler and incubate at 65°C for 5 min.
61. After incubation, place tubes on ice and preheat the thermocycler to the next phase of the program.
62. Add the following components into each tube on ice.

63. Mix by pipetting and centrifuge briefly.
64. Transfer tubes to preheated thermocycler and resume the program to incubate at 50°C for 60 mins, 85°C for 10 mins, and 4°C hold.
65. Once first strand cDNA synthesis is complete, add 2U RNase H to each reaction. Mix gently and centrifuge briefly.
66. Incubate in a thermocycler at 37°C for 20 mins.
67. Clean up PCR reactions with QIAquick PCR Purification Kit (Qiagen) according to manufacturer’s instructions.

| Component | 1 reaction (ul) |
| --- | --- |
| 5X RT Buffer | 4 |
| 100uM RNaseOUT RNase Inhibitor | 1 |
| Maxima H Minus Reverse Transcriptase (200U/ul) | 1 |
| *Total* | 6 |

a. Add 105ul (5 volumes) Buffer PB to each reaction and mix by pipetting.
b. Transfer to QIAquick column and spin at 10,000 xg for 1 min.
c. Add 700ul Buffer PE to wash the column and spin at 10,000 xg for 1 min.
d. Remove residual wash buffer by spinning at 10,000 xg for 1 min.
e. Transfer column to a clean RNase- and DNase-free tube.
f. Elute cDNA by adding 30ul Buffer EB directly to the membrane and incubate for 1 min at RT.
g. Centrifuge at 10,000 xg for 1 min.
68. Quantify single-stranded cDNA concentration and yield using the Qubit ssDNA assay.

This is a safe stopping point and the cDNA can be stored at 4°C for 48 hours or −20°C for weeks, or proceed to the next step.

#### Library density qPCR (timing: 1.5 hrs) (optional quality control)

A library density qPCR assay can be performed to assess relative expression of tdTomato containing SRTs as a preliminary quality control checkpoint. Results can inform a go/no-go decision for individual samples to prevent unnecessary labor and use of reagents. Example qPCR results are shown in **Figure 7.**

69. Depending on the single-stranded cDNA concentration determined above, use 1-10ng cDNA per reaction.
70. Prepare the appropriate amount of reagents based on the number of reactions according to the table below. At least 3 replicates of each reaction are recommended. Include no template control reactions to identify PCR contamination.

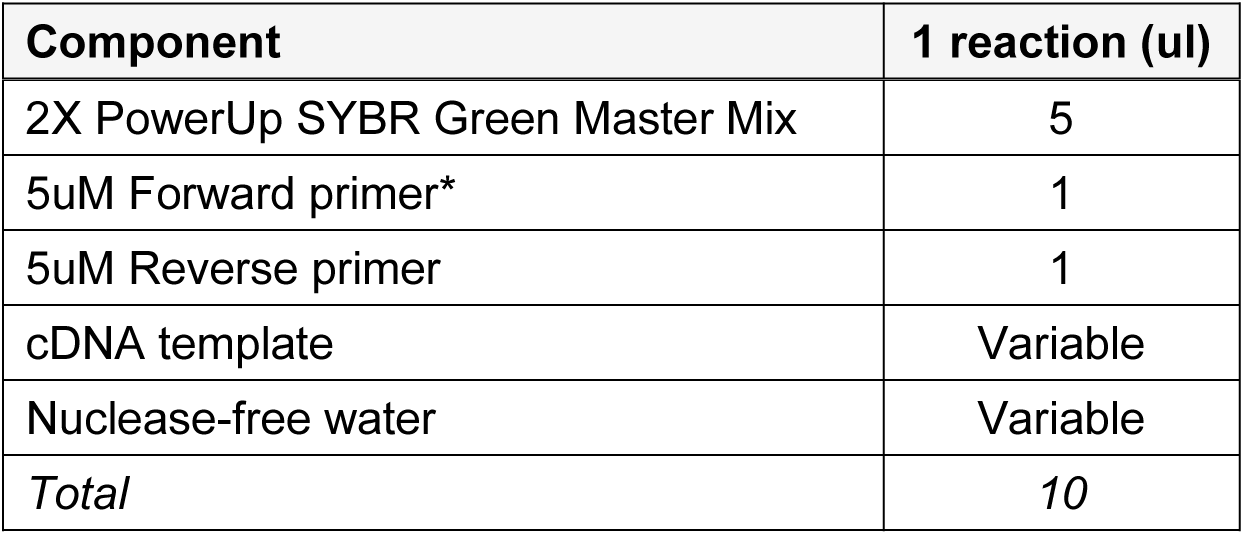 *tdTomato, B-actin, and Gapdh primer sequences can be found in **Table 2**.
71. Transfer the reaction mixes to each well of a 96 or 384 well plate according to the plate design and layout.
72. Seal the plate with an optical adhesive cover and centrifuge at 100 xg for 1 min to remove any air bubbles and ensure all liquid is at the bottom of each well.
73. Place the plate in the qPCR machine and set up the instrument to run using the following cycling conditions followed by a melt curve.

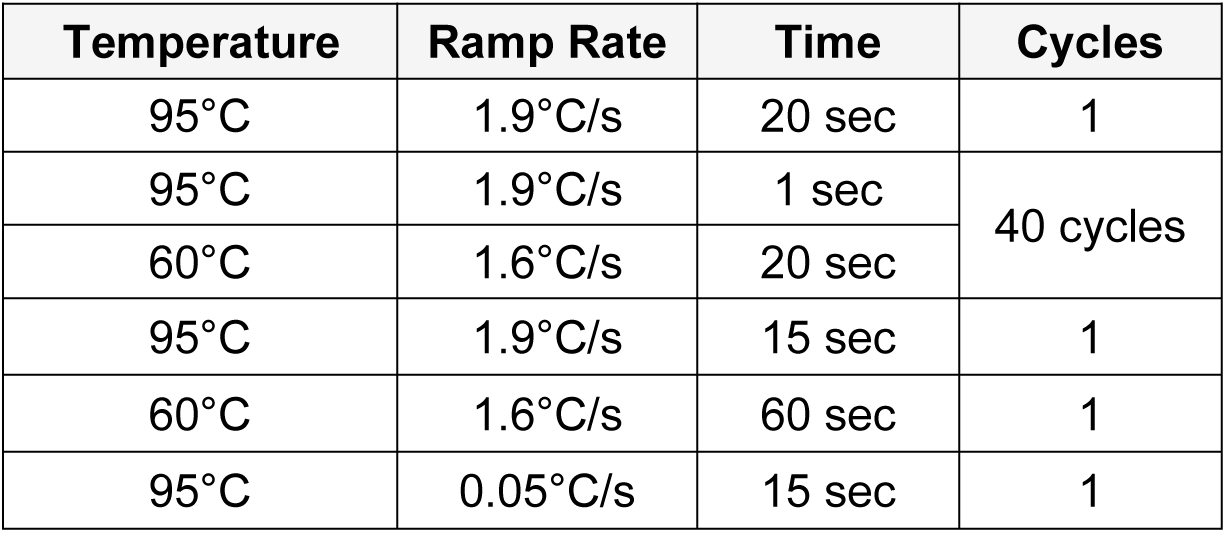
74. Once the run is finished, calculate the relative expression of tdTomato to B-actin (or other housekeeping gene). *NOTE: Samples with no or extremely low tdTomato expression should be omitted from subsequent library preparation steps. Thresholds for determining failed samples will need to be determined empirically, but generally C_T_ values greater than 30 cycles will likely not yield libraries. Examples are shown in* ***Figure 7C***.

**Figure 7.**
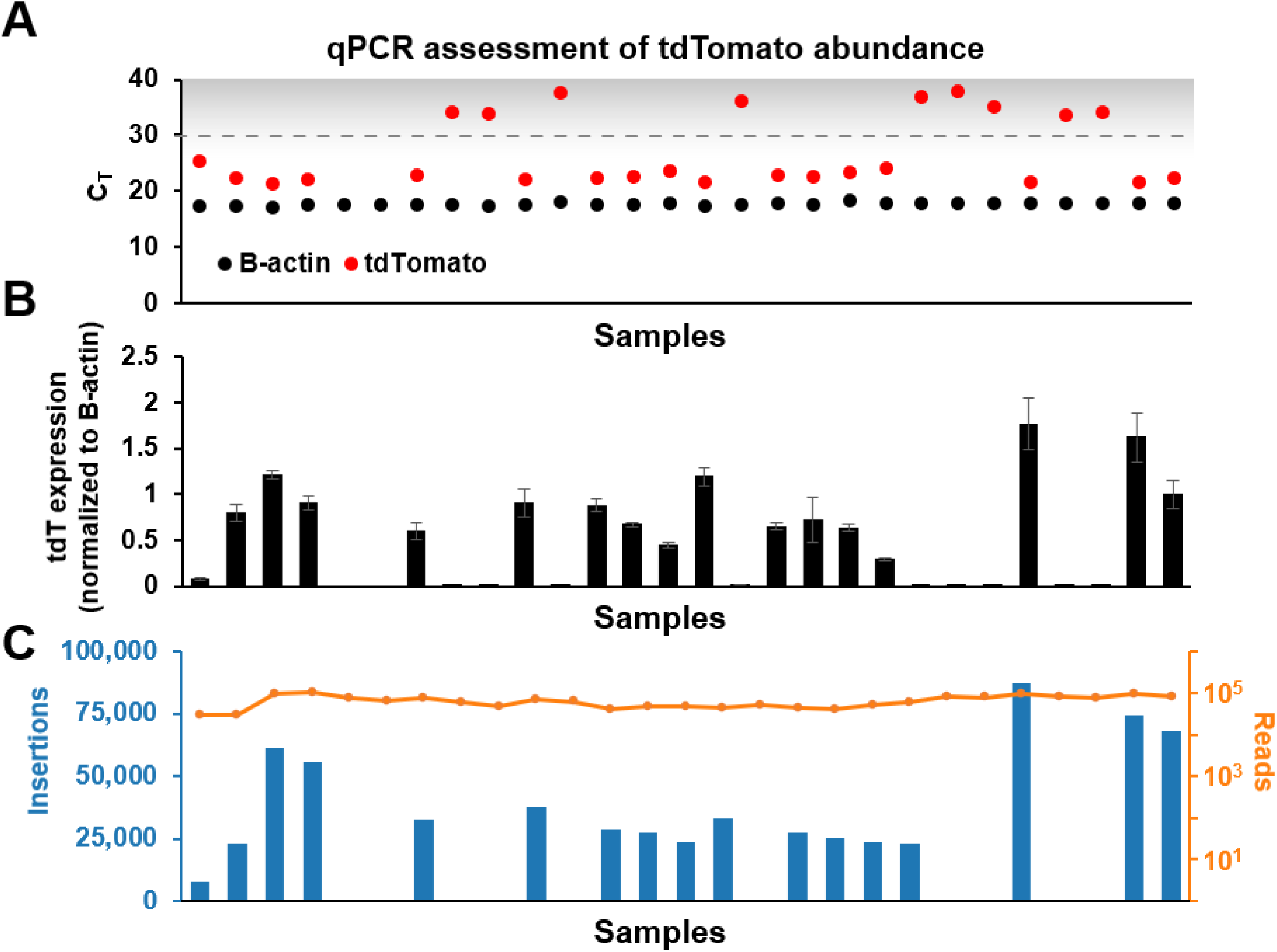
Representative quality control of cDNA and library complexity qPCR. **(A)** Example data from a qPCR library density assay using samples with a range of Calling Cards expression. Samples with a C_T_>30 have minimal number of tdTomato transcripts and can be omitted from downstream processing. **(B)** Plot showing tdTomato expression normalized to B-actin expression. **(C)** Plot showing the number of recovered insertions (blue) with number of shallow sequencing reads (orange) per sample. In this example, insertions were recovered from samples with a tdTomato C_T_<30.

#### Amplification of self-reporting transcripts (timing: 3 hrs, 20 mins hands-on)

This PCR step will amplify cDNA that contain SRTs for downstream tagmentation, indexing, and sequencing.

75. Set up and preheat a thermocycler with the following PCR program.

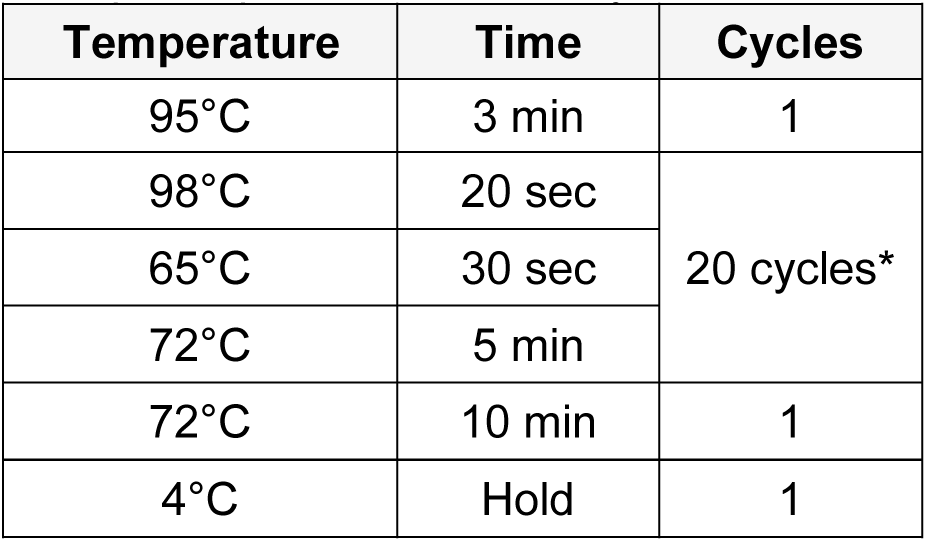 *\*NOTE: the number of cycles will need to be optimized per tissue/cell type to generate enough PCR product for downstream tagmentation. It is important to avoid overamplification, which can introduce amplification bias and PCR duplicates*.
76. Add the following components into individual nuclease-free tubes on ice.

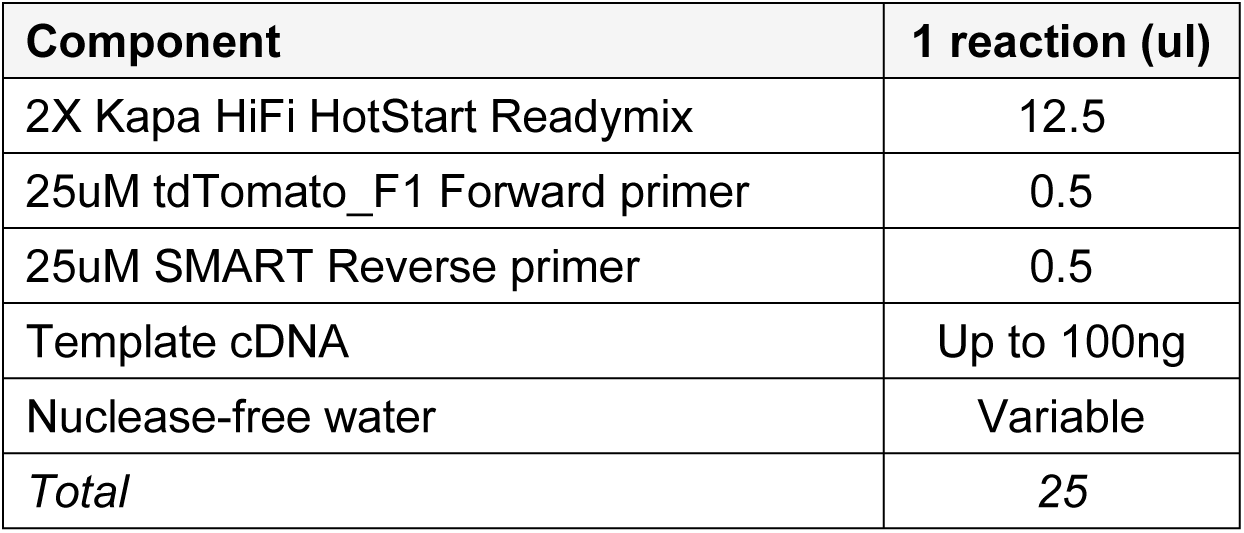 *NOTE: It is recommended to maximize the amount of template cDNA into this PCR step to maximize recovery of Calling Card insertions (****Figure 8***).
77. Transfer tubes to the preheated thermocycler and start the program.

**Figure 8.**
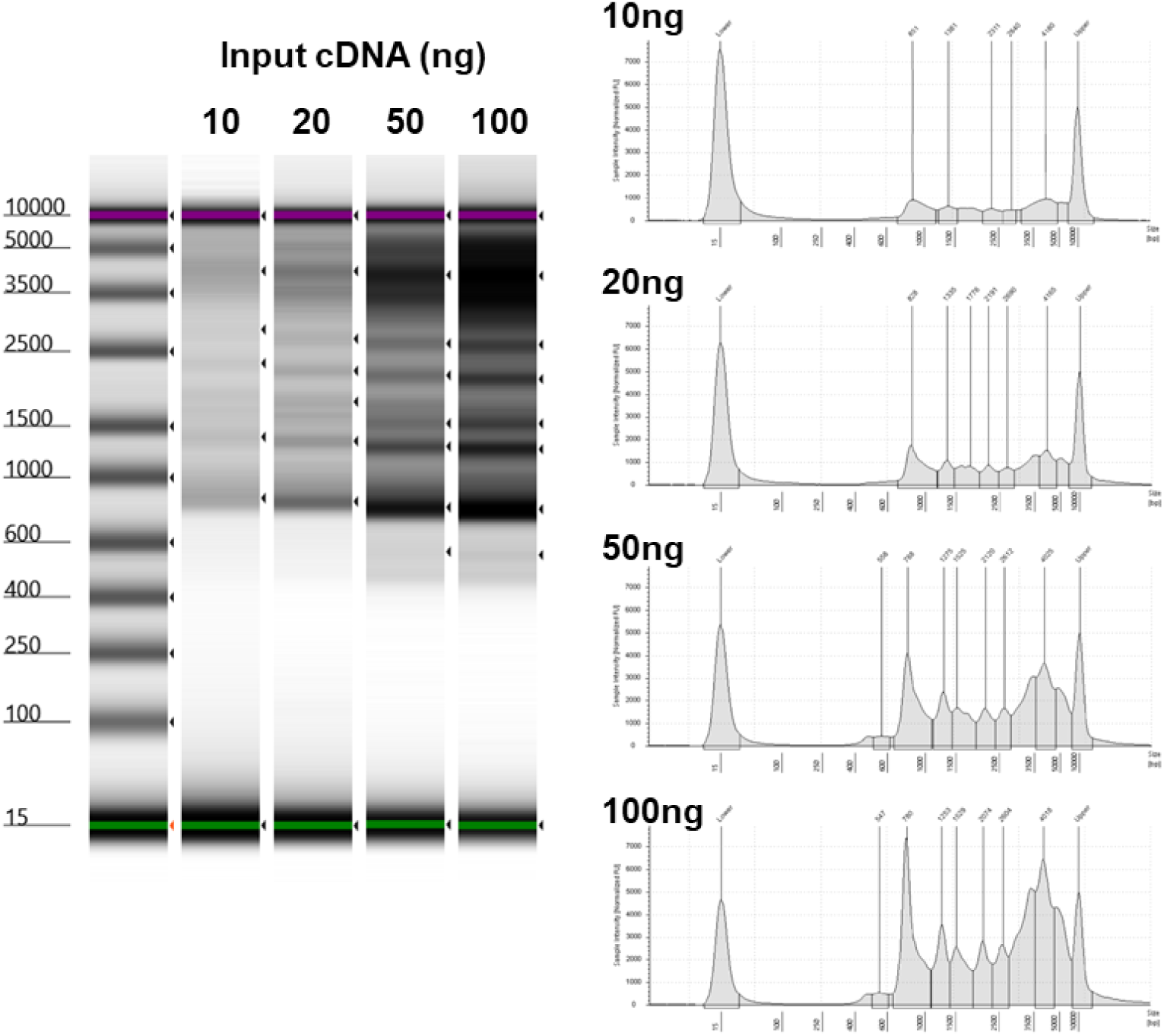
Titration of input cDNA into PCR to amplify SRTs. Example Tapestation gel images and electropherogram traces demonstrating that larger amounts of starting material (up to 100ng) can be used to increase yield of SRTs.

This is a safe stopping point and the DNA can be stored at 4°C for 48 hours or −20°C for weeks, or proceed to the next step.

#### Bead cleanup of PCR products and QC (timing: 30-45 mins)

Magnetic beads are used to clean up the SRT PCR products. If this is your first time working with magnetic beads, it is recommended to consult the manufacturer’s documentation for best practices and tips. For all wash steps, use freshly made 80% ethanol.

*NOTE: AMPureXP beads (Beckman Coulter) and Mag-Bind TotalPure NGS (Omega BioTek) magnetic beads have both been tested and validated*.

78. Bring AMPure XP beads to room temperature for at least 30 mins. *NOTE: vortex to completely resuspend beads immediately prior to use*.
79. Add 25ul of nuclease-free water to each PCR reaction to bring the volume up to 50ul.
80. Add 30ul AMPure XP beads (0.6X) to each sample and mix thoroughly by pipetting or vortexing.
81. Incubate on bench for 5 min.
82. Place on the magnetic rack for 1 min or until the solution clears. Without disturbing the beads, aspirate and discard the supernatant.
83. Add 200ul 80% ethanol, making sure not to disturb the bead pellet. Incubate for 30 sec.
84. Aspirate supernatant and discard.
85. Repeat wash by adding 200ul 80% ethanol to each sample. Incubate for 30 sec.
86. Aspirate supernatant and discard.
87. Pulse-spin the strip tubes to collect any remaining ethanol. Place on the magnetic rack and aspirate and discard residual ethanol.
88. Air dry the pellet for 1 min or until the beads become matte and lose their shine. *NOTE: do not over dry the beads (they will appear cracked) as this will decrease elution efficiency!*
89. Remove the strip tubes from the magnetic rack. Add 11ul Buffer EB to elute PCR products. Mix thoroughly by pipetting.
90. Incubate on bench for 2 min.
91. Place on a magnetic rack for 1 min, or until supernatant is clear.
92. Transfer 11ul supernatant of each sample to a new strip tube. *NOTE: it is important to ensure that there is no bead carryover as this can affect downstream steps. The supernatant should be completely clear*.
93. Quantify the concentration and visualize PCR products by running a 1:10 diluted sample on an Agilent High Sensitivity D5000 ScreenTape device or Bioanalyzer High Sensitivity DNA kit. Example gel images and traces are shown in **Figure 9A,B**. *NOTE: Alternatively, the Qubit dsDNA High Sensitivity kit can be used to quantify concentration of SRTs*.

**Figure 9.**
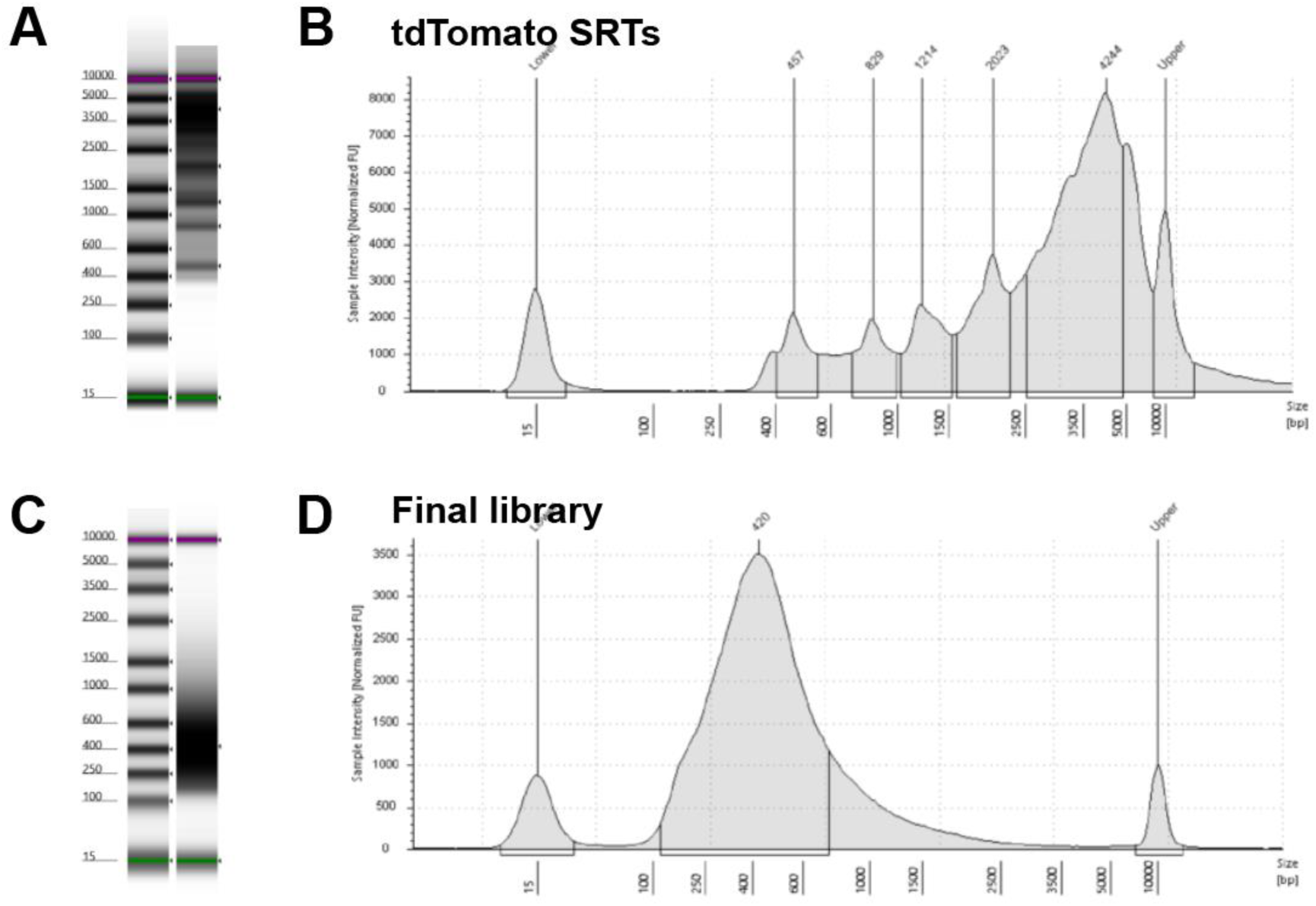
Representative quality control of SRTs and final library. Representative gel images and electropherogram traces for tdTomato SRTs after bead cleanup **(A,B)** and final sequencing library **(C,D)**.

This is a safe stopping point and the DNA can be stored at 4°C for 48 hours or −20°C for weeks, or proceed to the next step.

#### Tagmentation and indexing PCR (timing: 1-1.5 hrs)

In this tagmentation reaction, the Nextera XT transposome will enzymatically cleave the cDNA into smaller fragments and tag them with a Nextera overhang. The subsequent indexing will add a unique identifier to each library and enables multiple libraries to be pooled and sequenced together. Proper tagmentation and indexing of the library to a final size of 200-1000bp is optimal for efficient clustering on the sequencer flow cell.

94. Set up and preheat a thermocycler with the following PCR program.

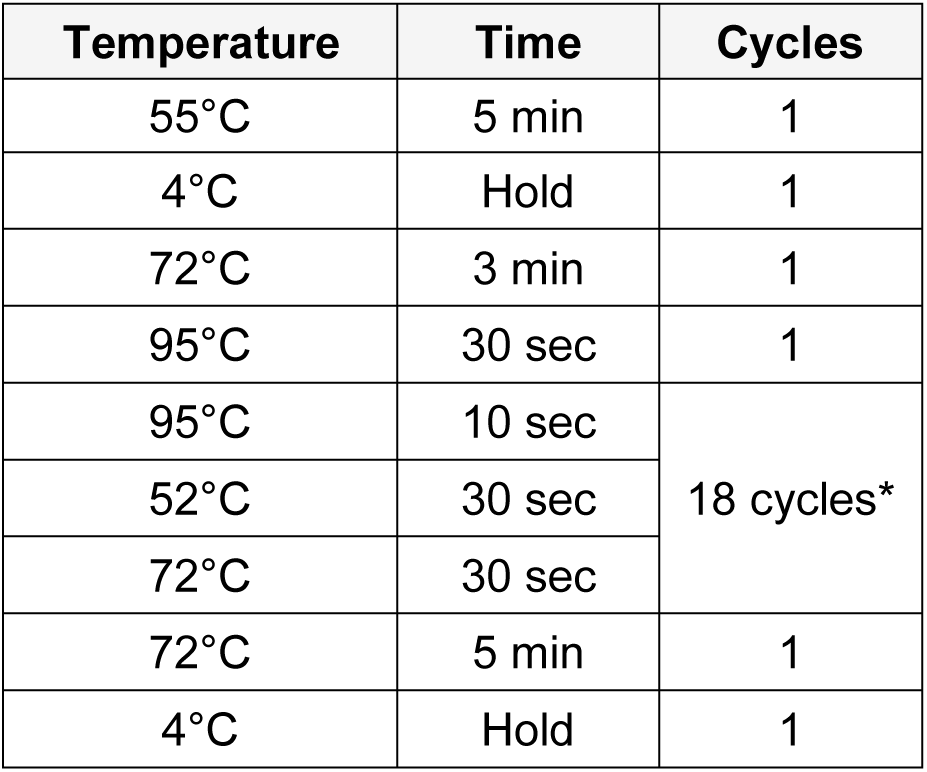 *\*NOTE: The number of cycles will need to be optimized per tissue/cell type to generate enough PCR product for downstream sequencing*.
95. Dilute amplified SRT PCR products to 300pg/ul with nuclease-free water.
96. Add the following components into individual nuclease-free tubes on ice.

97. Pipette to mix and briefly spin down. Transfer to the thermocycler preheated to 55°C and incubate for 5 min.
98. After the incubation, add 5ul Neutralization Tagment (NT) Buffer to stop the tagmentation reaction. Pipette to mix and briefly spin down. Incubate at room temperature for 5 mins.
99. Add the following to each PCR tube.

95. Dilute amplified SRT PCR products to 300pg/ul with nuclease-free water. 96. Add the following components into individual nuclease-free tubes on ice.
| Component | 1 reaction (ul) |
| --- | --- |
| Nextera Tagment DNA (TD) Buffer | 10 |
| Amplicon Tagment Mix (ATM) | 5 |
| Nuclease-free water | 3 |
| 300pg/ul SRT DNA | 2 |
| *Total* | *20* |

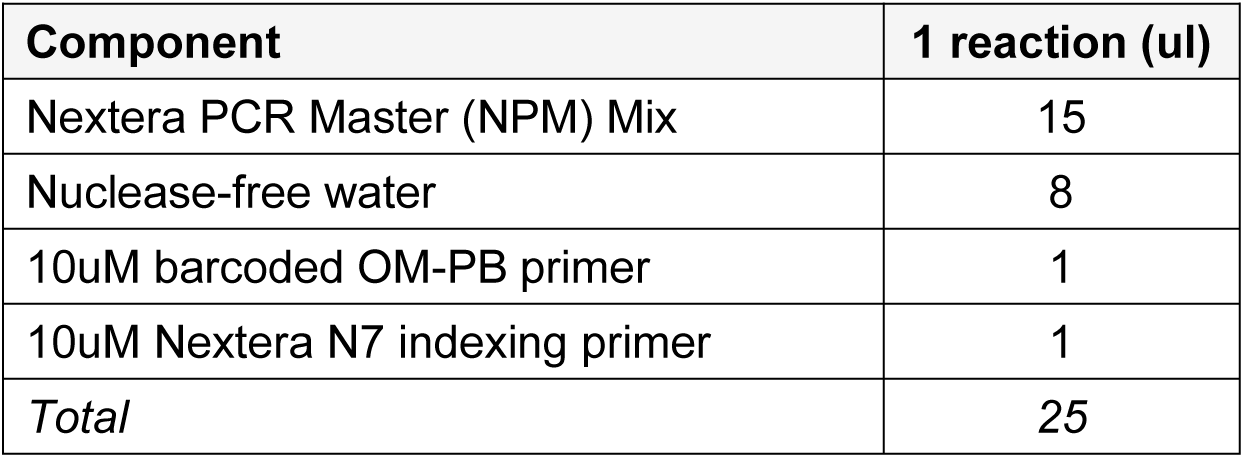 *NOTE: each replicate should receive a unique barcode-index combination (see “Distinguishing between biological and technical replicates” for an example barcode-index strategy)*.
100. Pipette to mix and briefly spin down. Transfer to the thermocycler preheated to 95°C and start the program.

This is a safe stopping point and the tagmented DNA can be stored at 4°C for 48 hours or −20°C for weeks, or proceed to the next step.

#### Bead cleanup of PCR products and QC (timing: 30-45 mins)

Magnetic beads are used to size-select and clean up the tagmented and indexed PCR products. Repeat steps 80-92 above from the previous “Bead cleanup of PCR products and QC” section.

101. Quantify the concentration and visualize PCR products by running a 1:10 diluted sample on an Agilent High Sensitivity D5000 ScreenTape device or Bioanalyzer High Sensitivity DNA kit. Libraries should be smoothly distributed between 200-1000bp. Example gel images and traces are shown in **Figure 9C,D**.

This is a safe stopping point and the libraries can be stored at 4°C for 48 hours or −20°C for weeks, or proceed to the next step.

## BASIC PROTOCOL 4: LIBRARY POOLING AND SEQUENCING

Libraries that were generated using the same protocol, have similar distributions of fragment sizes, and are uniquely indexed can be pooled and sequenced together. Balancing and pooling libraries is relatively simple, but accurate quantification of concentration is essential for cluster generation on the Illumina flow cell. Calling Card libraries are typically low yield libraries and thus bead-based normalization methods are not recommended. Normalizing by qPCR is a sensitive method to quantify adaptor-ligated templates and is the recommended method, however it is relatively time consuming and requires more reagents than other methods. An alternative is to pool samples equimolar based on Tapestation or Bioanalyzer values. The final combined library concentration should ideally be at least 5nM, however low molarity pools ∼1nM have been successfully sequenced without any loss of quality.

### Materials

Kapa Library Quantification Kit (Roche KK4824, or formulation compatible with your qPCR machine), or NEBNext Library Quant Kit for Illumina (NEB E7630)
QuantStudio 6 Flex Real-Time PCR System (ThermoFisher 4485691), or equivalent system for SYBR-green qRT-PCR
96 well metal cooling block (Argos Technologies 63615-04), or equivalent

#### Library quantification by qPCR (timing: 1.5 hrs)

This protocol is only needed if libraries are being quantified by qPCR. If libraries are being pooled based on Tapestation or Bioanalyzer molarity values, skip to the next section “Library pooling by Tapestation or Analyzer”.

102. Prepare samples and plate according to kit manufacturer’s instructions. *NOTE: For the Kapa kit, be sure to obtain the kit with the compatible passive reference dye formulation for your qPCR instrument. Similarly, for the NEBNext kit, only add an appropriate volume of ROX according to qPCR instrument requirements. Consult the qPCR instrument technical manual to verify ROX parameters*.
103. Run the qPCR assay according to the manufacturer’s instructions.
104. *Optional:* run a melt curve analysis to detect adaptor dimers.
105. Determine the concentration of the library samples using the standard curve generated by the DNA standards.
106. For each library, use the sample’s average fragment size to determine molarity.
107. Normalize each sample to the sample with the lowest molarity and pool samples equimolar for sequencing. *NOTE: samples with a molarity <1nM should not be included in the pool. The final pool should ideally be at least 5nM*.
108. Submit for Illumina next-generation sequencing.

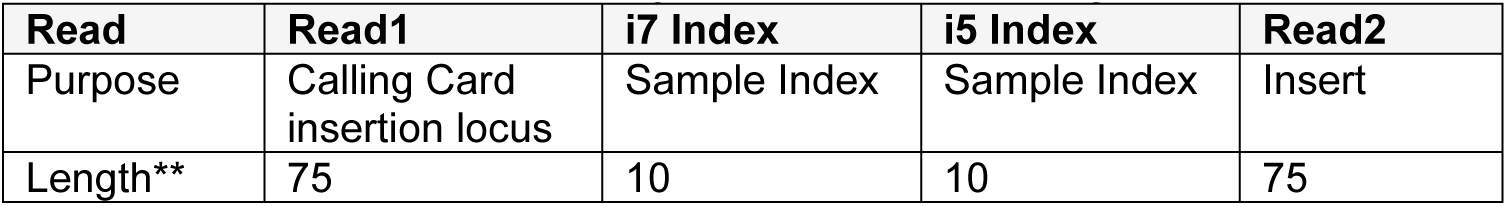 *NOTE: a read length of 75bp is the minimum recommended read length. Shorter reads may result in reduced alignment rates.*

#### Library pooling by Tapestation or Bioanalyzer (timing: 15 mins)

This protocol is only needed if libraries are not quantified by qPCR.

109. Normalize each sample to the sample with the lowest concentration and pool samples equimolar for sequencing. *NOTE: samples with a molarity <1nM should not be included in the pool. The final pool should ideally be at least 5nM*.
110. Submit for sequencing.

#### Sequencing (timing: variable)

Sequencing uses standard Illumina NGS sequencers, typically provided by a local genomics core facility, MGI@GTAC, or commercial sequencing vendor. If it is available and time efficient with your local genomics core, first-pass shallow sequencing at a depth of at least 100k single or paired end 150bp reads per sample within the pooled library is recommended to perform preliminary sequencing QC. This step ensures high quality library preparations prior to committing and investing in resources for deep sequencing. Metrics such as sequencing read counts, base call qualities, adapter content, alignment scores, and overrepresented sequences are representative of the whole pool even at low sequencing depth.

The full sequencing depth needed to recover all Calling Card insertions in a sample is variable depending on the complexity of the library and number of insertions. In our experience, 10-25M reads per sample is sufficient to reach ∼90% saturation for most samples (**Figure 10A**). To estimate sequencing saturation, unique reads and alignments within .bam files can be downsampled to simulate less sequencing. These points can be plotted, and a logarithmic growth curve can be fitted to the points to estimate the number of reads needed to reach >90% sequencing saturation.

**Figure 10.**
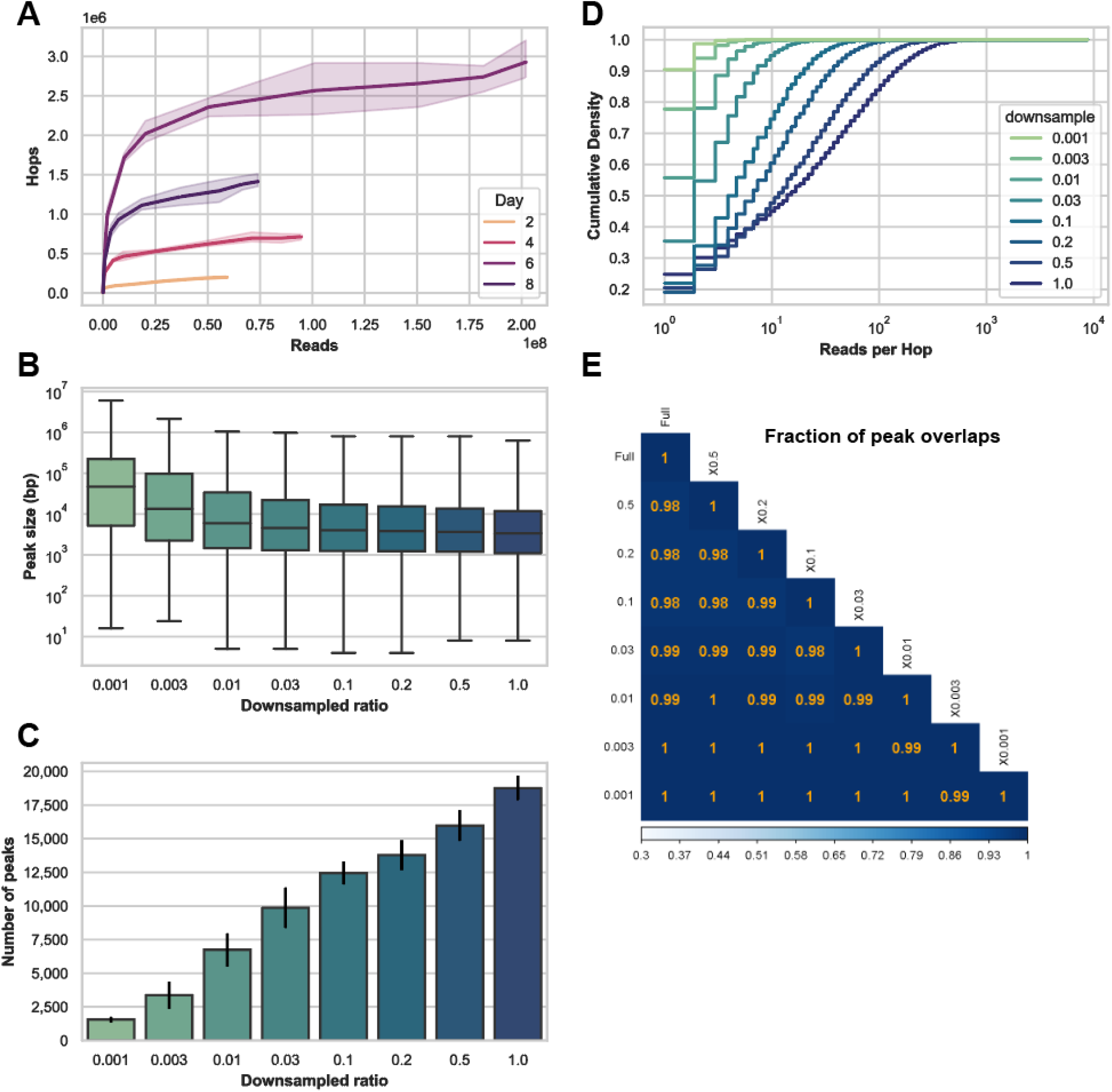
Sequencing saturation and called peaks. **(A)** Plot of insertion densities at various downsampled read depths for samples collected 2-8 days after injection with Calling Cards reagents (see Figure 5 for experimental design). Panels B-E are based on the deeply sequenced Day 8 sample in A. The BAM file was downsampled at set ratios to simulate a range of sequencing depths. **(B)** Box plots showing the distribution of the sizes of called peaks. **(C)** Bar plots demonstrating that the number of called peaks increases with deeper sequencing. **(D)** Cumulative density plots of the number of reads per Calling Card insertion. **(E)** Heatmap showing that the called peak regions are virtually identical with different sequencing depths. Taken all together, shallower sequencing will lead to few broad peaks while deeper sequencing will increase the resolution and result in more peaks that are narrower.

**Figure 11.**
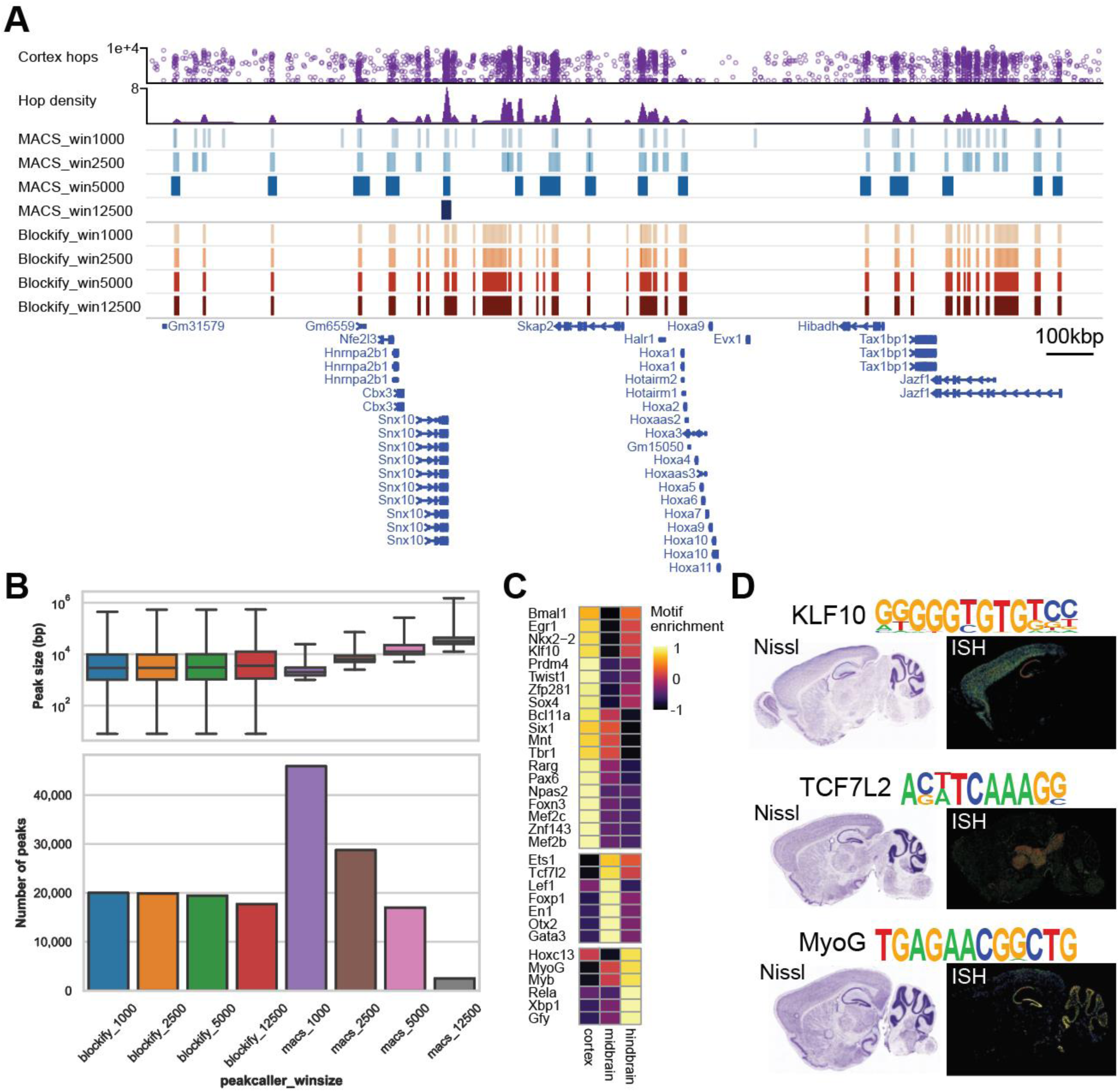
Comparison of peak callers and determining optimal parameters. **(A)** Representative genome browser tracks of raw Calling Cards insertions and computed density. A range of window sizes are used with a custom peak caller based on the MACS algorithm (blue) and Blockify (red), which is based on the Bayesian blocks algorithm. Both were tested with a range of ‘window size’ settings (win1000-win12500). MACS_win2500 was chosen for downstream analysis because it captured major peak regions without overcalling peaks. **(B)** Box and bar plots showing the distribution of peak sizes and number of peaks as a function of peak caller and the window size parameter. **(C)** Heatmap showing the enrichment scores of TF motifs found specific to the cortex, midbrain, and hindbrain regions. **(D)** Representative Nissl and ISH images taken from the Allen Brain Atlas ISH database (https://mouse.brain-map.org/) confirm brain region specific expression of TFs corresponding to the Calling Cards identified genomic regionally enriched motifs.

The MiSeq, MiniSeq, NextSeq550, and NovaSeq6000 platforms have been successfully used to sequence Calling Card libraries. The Genome Technology Access Center at the McDonnell Genome Institute (GTAC@MGI; https://gtac.wustl.edu/) is familiar with and offers a service to sequence Calling Card libraries on the NovaSeq6000 platform, available to any laboratory. General guidelines for in-house sequencing, or to share to your sequencing provider, are provided below.

Due to the low complexity of bases in the first 38 bases of R1, we recommend that Calling Card libraries are not sequenced by themselves on their own lane, but rather multiplexed with other library types. Otherwise, 50% co-loading with a complex sequencing library (e.g. PhiX DNA) is required for MiniSeq, NextSeq550, and NovaSeq6000 platforms. For the MiSeq, a cluster density of 750k/mm^2^ is targeted and libraries are loaded at a concentration of ∼30nM with 10-15% PhiX. For the MiniSeq and NextSeq550 platforms, pooled libraries are loaded at a concentration of ∼3nM to aim for a cluster density of 150 k/mm^2^. For the NovaSeq6000 platform, libraries were loaded at 1.5nM and sequenced on a S4 flow cell using the Xp workflow and a 151×10×10×151 sequencing recipe according to the manufacturer protocol. Reads were demultiplexed by sample index (i5/i7) using bcl2fastq allowing for 1 mismatch without adapter trimming (see official documentation for details; https://support.illumina.com/sequencing/sequencing_software/bcl2fastq-conversion-software.html).

111. Transfer raw `fastq` files from source location to your cluster or compute environment using research data management software (e.g. Globus) or command-line tools (e.g. `rsync`).

## BASIC PROTOCOL 5: DATA ANALYSIS (TIMING: VARIABLE)

The bioinformatics analysis pipeline has been built as a `nextflow` workflow, which containerizes each process (Ewels et al., 2020). This not only greatly simplifies the maintenance of software dependencies, but also enables the deployment across a variety of different compute environments. A high-level overview is provided here and complete documentation can be found on the `nf-core/callingcards` main page (https://nf-co.re/callingcards).

*NOTE: this pipeline is being actively developed and there is a chance that some code written here may be different than the latest version. This guide is written based on the first stable release v0.0.1. See* https://nf-co.re/callingcards *for the most up to date documentation*.

### Materials

POSIX compatible system (e.g. Linux, macOS, etc.)
8-core Intel or AMD processor (16 cores recommended)
64GB RAM
500GB free disk space
Bash 3.2 or later

If running in cluster mode:

Shared filesystem
Batch scheduling system (e.g. SLURM, SGE, LSF, etc.)

#### Install and configure nextflow (timing: variable)

112. Install `nextflow 22.10.4` or later on your system. Details can be found on the official documentation (https://www.nextflow.io/docs/latest/getstarted.html).
113. Install a container engine of choice (e.g. `Docker`, `Singularity`, `Shifter`, `Podman`, or `Charliecloud`). For this guide, we will be using `Singularity`.
114. Test the installation and configuration with a minimal data set. This only tests the proper configuration of the pipeline and local compute environment. It should complete without errors in a few minutes and the output data can be disregarded. Note that some config profiles may be needed to instruct `nextflow` how to fetch the required software. In this example, test and singularity profiles are chained. The path to a custom `local_config` file is specified in the `–c` parameter. It is recommended to check if a config file is already available for your cluster from `nf-core/configs` (https://nf-co.re/usage/configuration). If one is not available, a tutorial for writing one can be found here (https://nf-co.re/docs/usage/tutorials/step_by_step_institutional_profile).

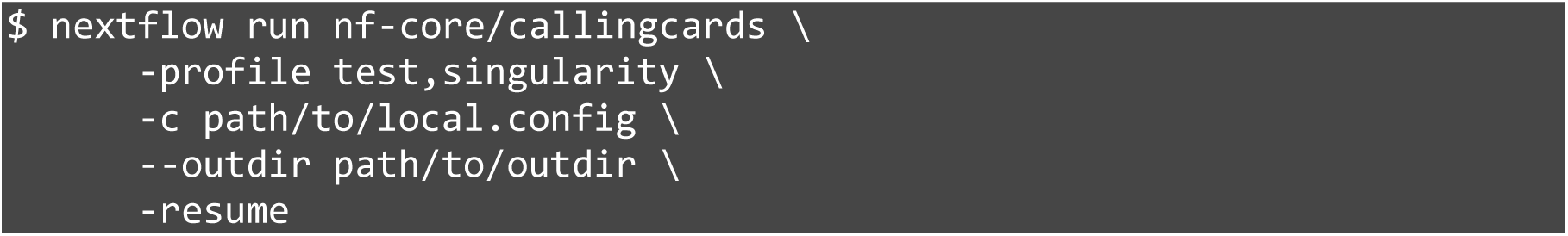 If the pipeline successfully completes, `nextflow` is correctly configured and loaded. The pipeline should create the following relative paths and files within the top-level project directory.

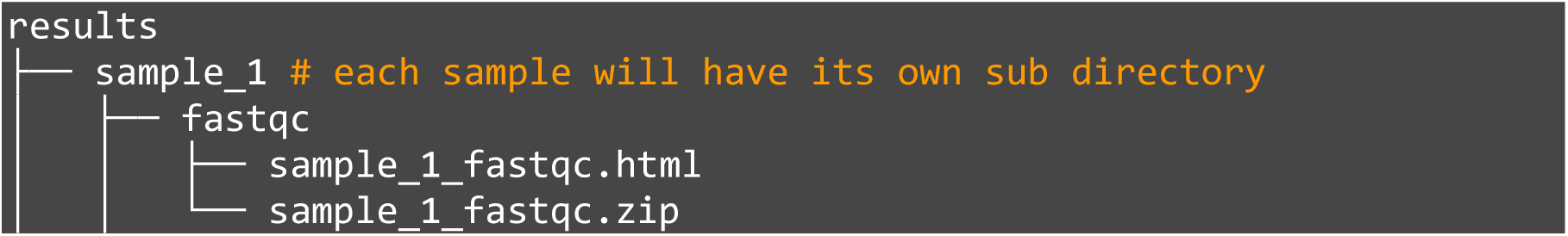
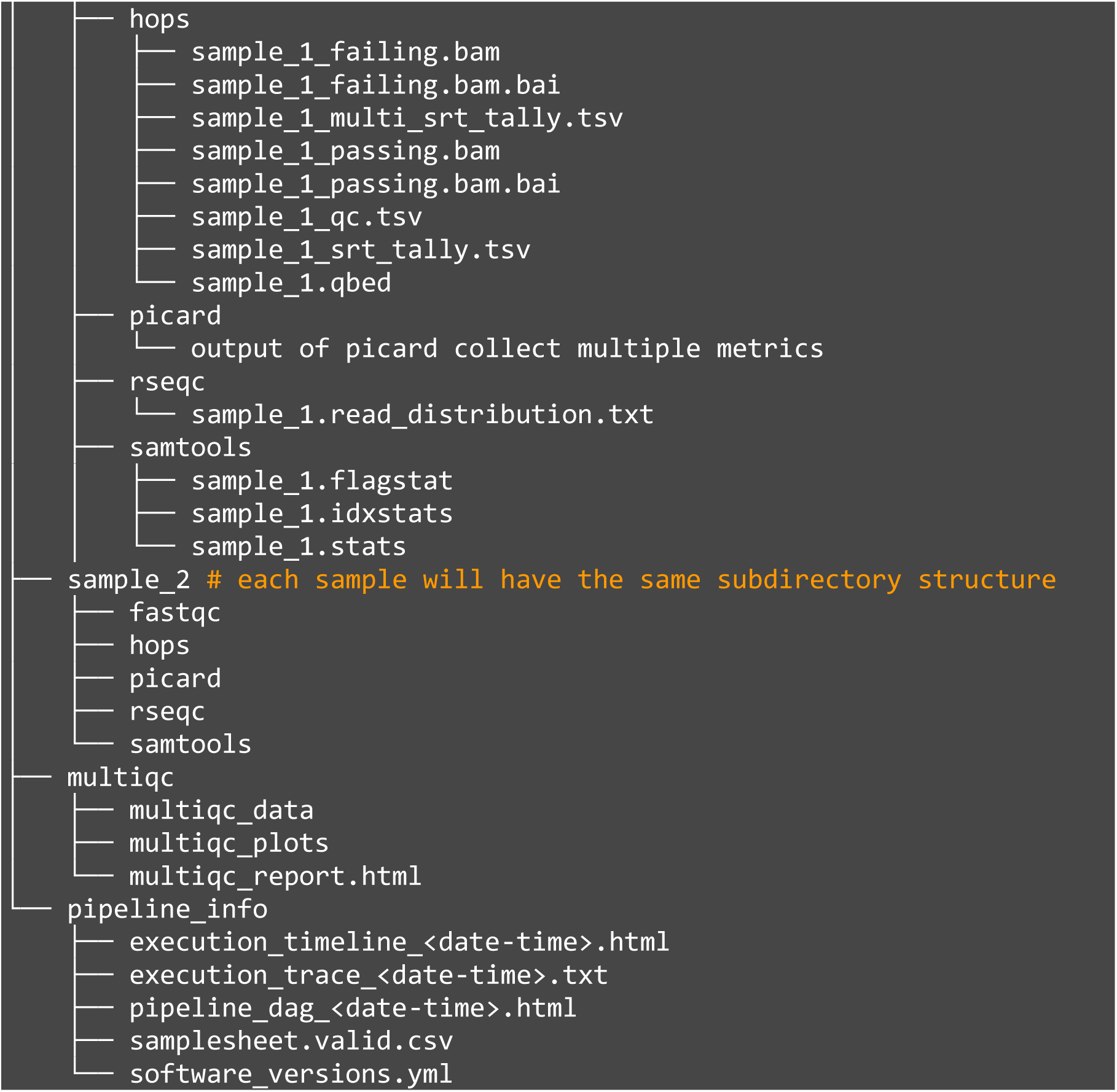

#### Run nf-core/callingcards on your own data (timing: variable)

115. Using a text editor, create a `samplesheet.csv` containing the information in the format below with each row being an independent sample to be analyzed.

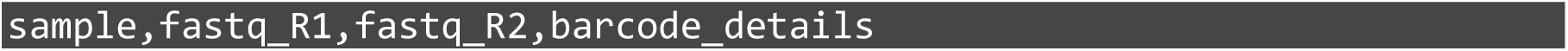

116. Create a `barcode_details.json` that provides SRT barcode information. An example json for a standard experiment using all available SRT barcodes is provided below. More complex experimental designs such as multiplexing transcription factors with subsets of SRT barcodes will require editing the SRT barcode array within the components object. Note that the 3bp OM-PB primer barcode will need to be specified in the `r1_pb` object.

| Fields | Description |
| --- | --- |
| sample | Custom sample name. This entry will be identical for multiple sequencing libraries/runs from the same sample. Spaces in sample names are automatically converted to underscores (_). |
| fastq_R1 | Full path to FastQ file for Illumina short reads 1. File has to be gzipped with the extension **.fastq.gz** or **.fq.gz**. |
| fastq_R2 | (Optional). Full path to FastQ file for Illumina short reads 2. File has to be gzipped with the extension **.fastq.gz** or **.fq.gz**. |
| barcode_details | Full path to the barcode details json file for a given sample. |

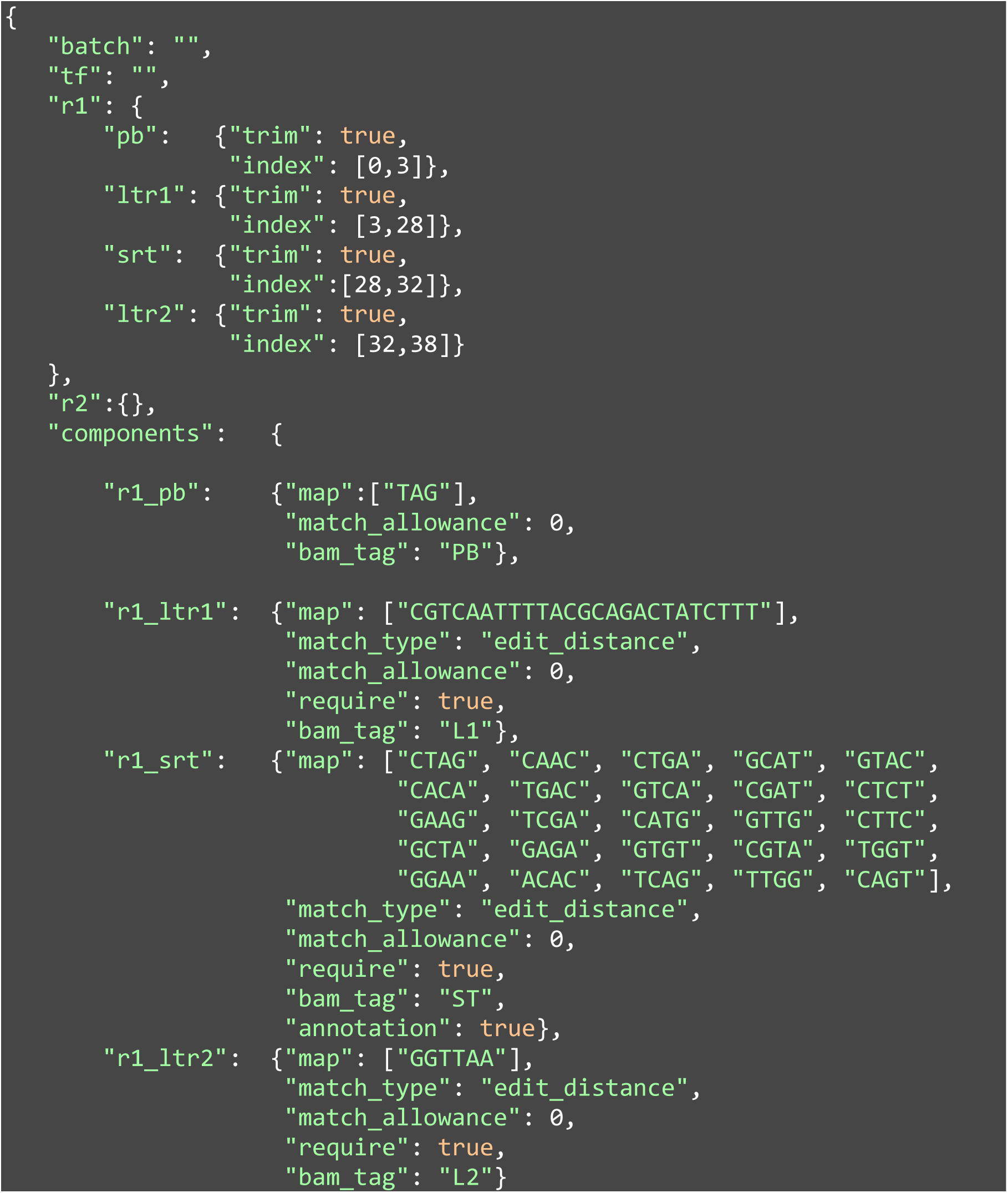
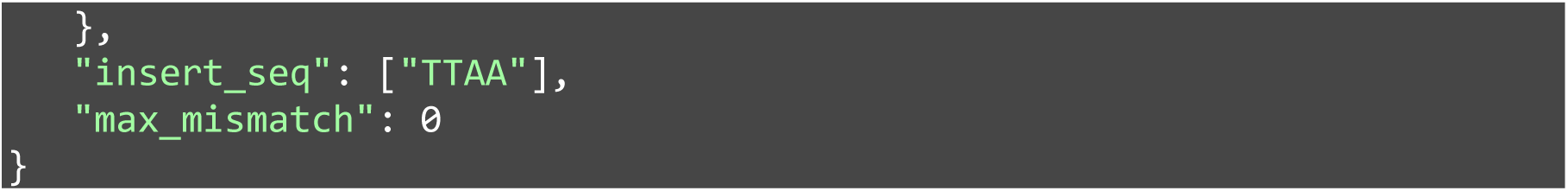

117. Create a `params.json` file which will save the input parameters in json format. An example is provided below using the `bwamem2` aligner.

| Fields | Description |
| --- | --- |
| batch | (Optional). Name of the experiment, e.g. `experiment_1234`. |
| tf | (Optional). Either ID or symbol of the TF. |
| r1 | Keys correspond to barcode component names. Each component is a map with keys trim which is boolean. Set to `true` to trim off this portion of the read (NOTE: not used in mammals pipeline. This functionality is instead controlled by UMITools). Index specifies where this component occurs in the read. |
| r2 | same as r1. |
| components | Each key corresponds to a component in `r1` or `r2` and must be preceded by `r1` or `r2` as appropriate. Each value is a map where the key map is required and lists the expected sequence(s) for that component. The key `r1_pb` holds the value of the 3bp OM-PB primer barcode. Optional additional keys are `match_allowance` to allow mismatches `>0`, `bam_tag` which is used to add the sequence extracted from the read to the aligned read, `match_type` which controls how the sequences are matched (default is `edit_distance` and appropriate in most circumstances), and `require` is a boolean which, when set to `false`, allows any number of mismatches in the component without penalty. |
| insert_seq | A list of sequences expected to be present on the 5' end of the aligned read. |
| max_mismatch | The maximum number of mismatches allowed in a barcode. By default this is equal to the sum of the mismatches allowed on each component. Set this to less than that sum to fail barcodes which pass component checks but have more than `x` number of mismatches overall. |

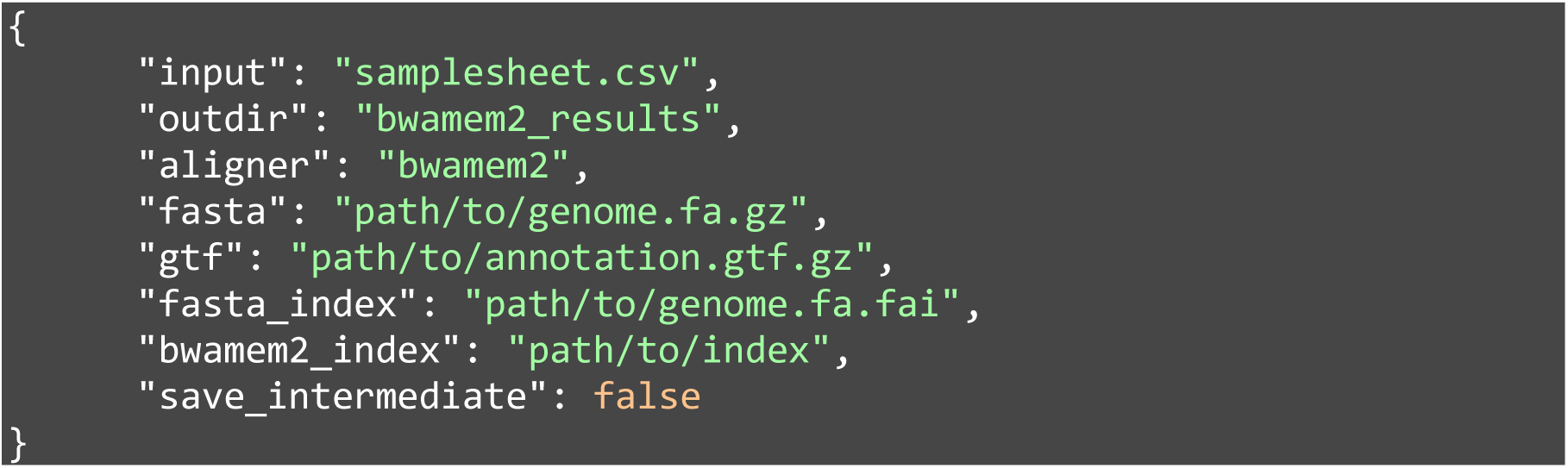
118. To run the pipeline with default parameters, use the following command (see https://github.com/nf-core/callingcards/blob/master/conf/default_mammals.config for more details).

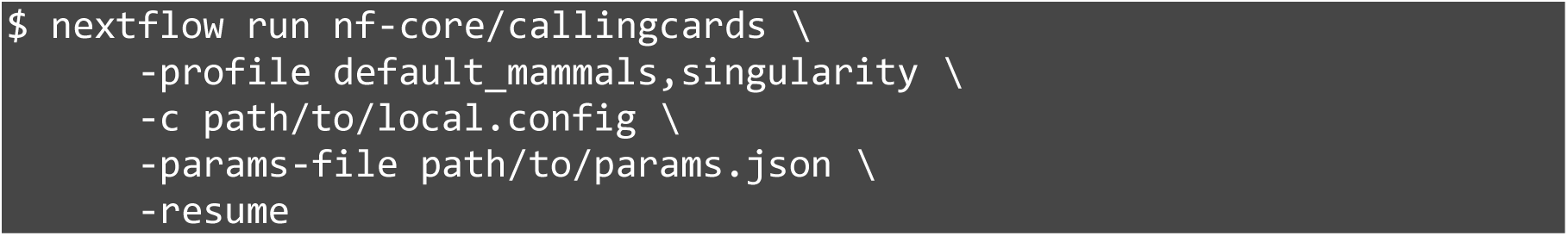 The `nextflow` pipeline checks the input `samplesheet.csv`, `params.json`, indexes the reference genome if not already done and in the working directory, prepares the reads, extracts SRT barcodes, trims adapter sequences, aligns the reads to a reference genome, processes the aligned reads to map the insertion locus, and outputs a `qBED` file (**Figure 2**). This file is then used to call peaks, calculate significance, motif enrichment, and other downstream analyses.
119. Examine the first 10 lines of the output `qBED`. Each line of the `qBED` file corresponds to a unique insertion. The columns represent chromosome, start coordinate, end coordinate, sequencing depth of that insertion, strand, and SRT barcode.

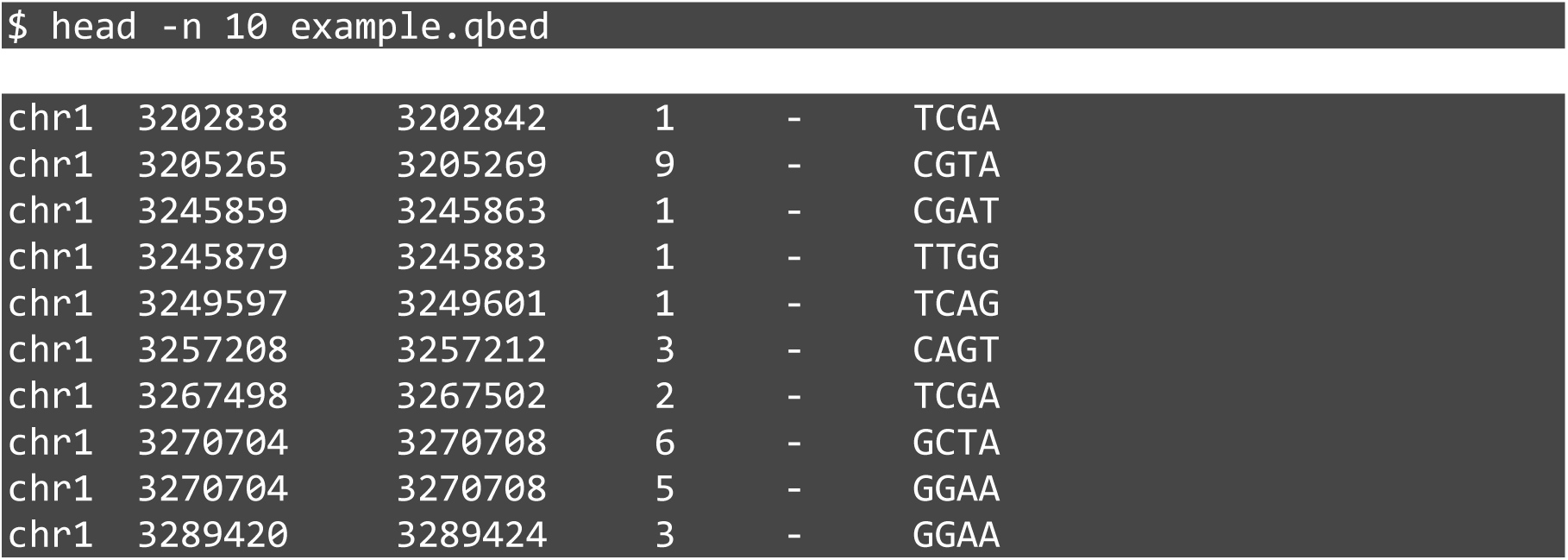

#### Visualization of Calling Cards data (timing: variable)

The WashU Epigenome Browser (http://epigenomegateway.wustl.edu) (Li et al., 2019) provides a streamlined platform to visualize Calling Card data using the `qBED` format, which reports transposon insertions as discrete points along the genome (x-axis) and the number of reads associated with that insertion (y-axis) (Moudgil et al., 2020a). This creates a scatterplot-like depiction that can be used to visualize densities of Calling Card insertions across the genome (Figure 1A). To load `qBED` tracks into the WashU Epigenome Browser, the file must first be sorted, compressed, and indexed.

120. Install a stable release of HTSlib (Bonfield et al., 2021) (https://github.com/samtools/htslib).
121. If not already sorted, sort the `qBED` file.

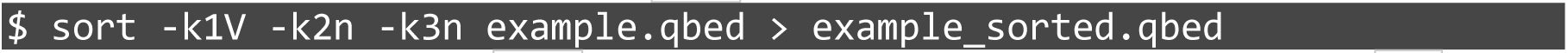
122. Block compress the `qBED` file. The output file will be a new file with the `.gz` extension and the original will be removed.

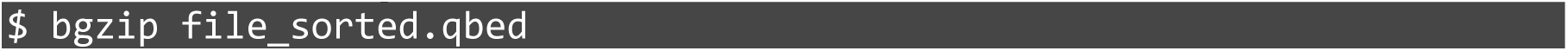
123. Index the compressed file in bed format. This will create `file_sorted.qbed.gz.tbi`

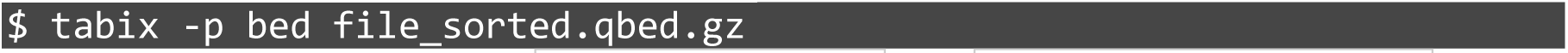
124. Store the compressed `qBED (*.qbed.gz)` and `index (*.qbed.gz.tbi)` together locally or in a web-accessible directory.
125. On the browser, select the desired reference genome (ie. `mm10, hg38`, etc.).
126. Upload the compressed `qBED` and index pair together to the browser. Select `qBED` as the track file type. If you are using local files, select `Tracks > Local Tracks`. Additional tracks and track types (ie. `BED`, `bedGraph`, `bigWig`, etc.), can also be compressed, indexed, and simultaneously loaded in a similar way by repeating steps 105-107).
127. Track parameters such as color, height, scale, opacity, and marker size can be customized from within the browser interactively. Publication quality `SVG` or `PDF` images can be captured and saved using the `Screenshot` tool (`Apps menu > Screenshot`). To load many tracks or to ensure reproducibility between sessions, parameters can be edited and loaded in JSON format. An example to load a `qBED`, `bedGraph`, and `BED` file for a single sample is shown below). All options can be found at https://eg.readthedocs.io/en/latest/usage.html#track-customization.

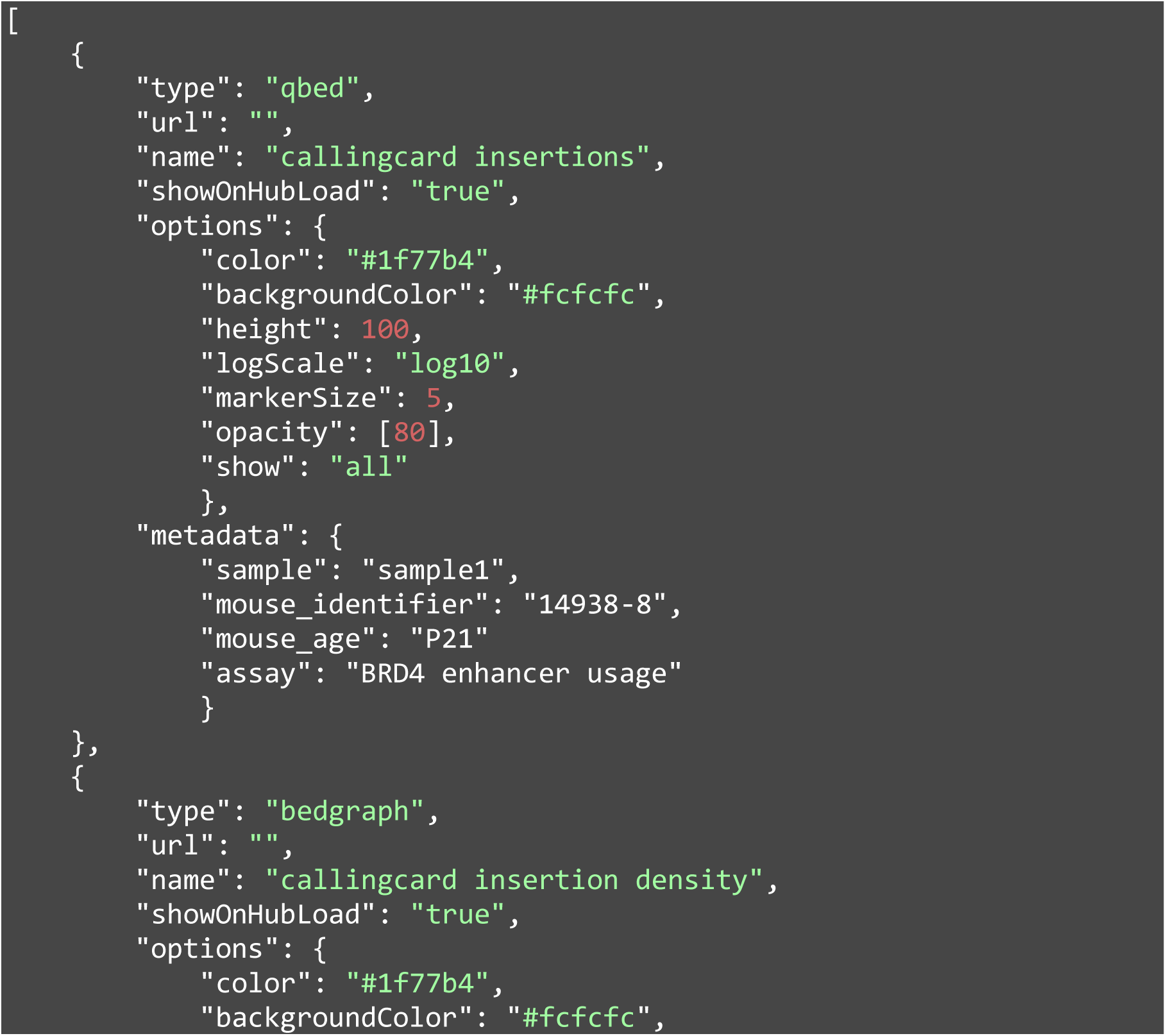
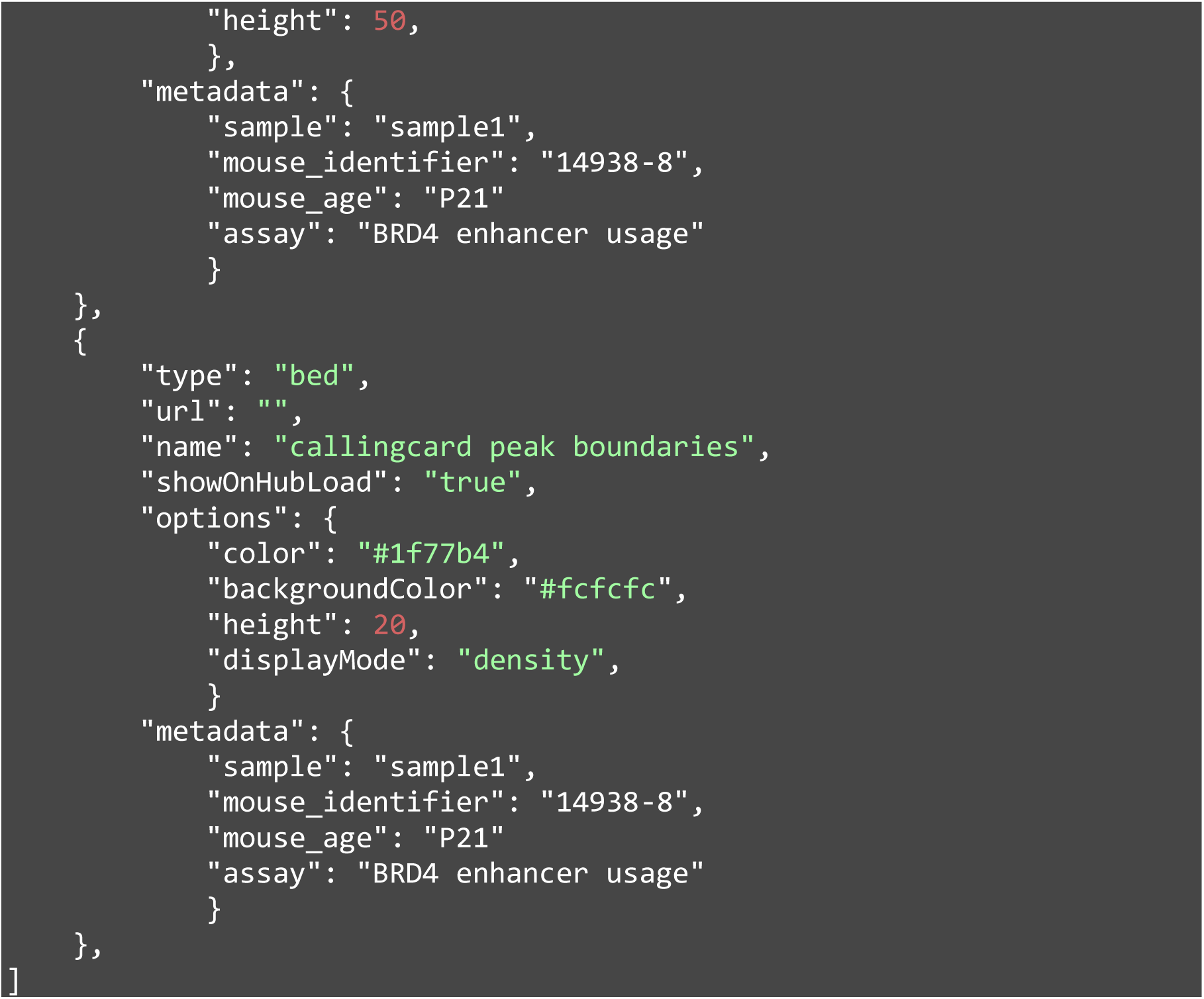

## SUPPORT PROTOCOL 1: NGS QUANTIFICATION OF BARCODE DISTRIBUTION WITHIN SRT PLASMID POOL AND AAV GENOME

Since the barcoded SRT plasmid pool now contains multiple elements, attention is needed to ensure that all elements are represented evenly, amplified, and that none are lost. Having an evenly distributed plasmid pool is optimal for transformation, outgrowth, and ultimately for generating an evenly distributed pool of packaged AAV particles. While Sanger sequencing provides the consensus sequence of the pool, next-generation sequencing can reveal the actual representation of the library. To do so, a fragment of the plasmid containing the SRT barcodes will be amplified by PCR and standard Illumina sequencing adapters will be ligated. These products will then be purified, quantified, and submitted for sequencing. Similarly, the AAV genomes from the packaged viral particles can be harvested and analyzed for the actual distribution of barcodes that were packaged.

### Materials

Q5 Hot Start High-Fidelity Master Mix (NEB M0494)
AMPure XP Reagent (Beckman Coulter A63882) or Mag-Bind TotalPure NGS beads (Omega Biotek M1378-02)

*NOTE: both magnetic beads have been tested and validated for clean-up and size selection of Calling Card libraries. If using the Mag-Bind beads, in-house validation of size selection ratios of each lot with a DNA ladder is recommended*.

T100 Thermal Cycler (Biorad 1861096), or equivalent
Magnetic Separation Rack for PCR strip tubes (Permagen MSR812, or equivalent)

#### Isolation of viral genome from AAV particles (timing: 3.5-4 hr)

The protocol is adapted from the ‘Digest virus particles to release DNA’ section of Support Protocol 1: Determination of rAAV Titers by the Dot-Blot Assay (ref).

128. Add 100ul DNase digestion buffer to clean 1.5ml microcentrifuge tubes. Pulse vortex concentrated virus and centrifuge to collect liquid.
129. Add 2ul virus into each 1.5ml tube. Pulse vortex and centrifuge to collect liquid. Incubate for 1 hr at 37°C.
130. Add 5ul of 0.5M EDTA to each tube to inactivate the DNase. Pulse vortex and centrifuge to collect liquid. Incubate for 10 min at 70°C.
131. Prepare fresh proteinase K solution and add 120ul to each tube. Pulse vortex and centrifuge to collect liquid. Incubate for 2 hr at 50°C.
132. Inactivate the proteinase K by incubating for 10 mins at 95°C.
133. Allow the tubes to cool and clean up reactions using QIAquick PCR Purification kit.

#### Library preparation (timing: 1h)

134. Prepare a 1-step sequencing library to assess barcode distribution of the pool.

135. Transfer tubes to a PCR machine and begin thermocycling with the following parameters.

| Component | 1 reaction (ul) |
| --- | --- |
| Q5 HF 2X Master Mix | 5 |
| 10uM SRT_bc_QC_F primer | 0.5 |
| 10uM SRT_bc_QC_R primer | 0.5 |
| 5ng plasmid pool | Variable |
| Nuclease-free water | Variable |
| *Total* | *10* |

135. Transfer tubes to a PCR machine and begin thermocycling with the following parameters.
| Temperature | Time | Cycles |
| --- | --- | --- |
| 98°C | 30 sec | 1 |
| 98°C | 5 sec | 10 cycles |
| 55°C | 10 sec |  |
| 72°C | 30 sec |  |
| 4°C | Hold | 1 |

This is a safe stopping point and the PCR amplicons can be stored at −20°C, or proceed to the next step.

#### Bead cleanup and quantification (timing: 30min)

136. Bring AMPure XP beads to room temperature for at least 30 mins. Vortex completely to resuspend beads immediately prior to use.
137. Add 40ul of nuclease-free water to the PCR reaction to bring the volume up to 50ul.
138. Add 50ul AMPure XP beads (1X) to the sample and mix thoroughly by pipetting or vortexing.
139. Incubate on bench for 5 min.
140. Place on the magnetic rack for 2 min or until the solution clears. Without disturbing the beads, aspirate and discard the supernatant.
141. Add 200ul 80% ethanol, making sure not to disturb the bead pellet. Incubate for 30 sec.
142. Aspirate supernatant and discard.
143. Repeat wash by adding 200ul 80% ethanol to each sample. Incubate for 30 sec.
144. Aspirate supernatant and discard.
145. Air dry pellet for 1 min or until the beads become matte and lose their shine. *NOTE: do not over dry the beads (they will appear cracked) as this will decrease elution efficiency!*
146. Pulse-spin the strip tubes to collect any remaining ethanol. Place on the magnetic rack and aspirate and discard residual ethanol.
147. Remove the strip tubes from the magnetic rack. Add 20ul Buffer EB to elute PCR products. Mix thoroughly by pipetting.
148. Incubate on bench for 2 min.
149. Place on a magnetic rack for 1 min, or until supernatant is clear.
150. Transfer 20ul supernatant to a new tube.
151. *NOTE: it is important to ensure that there is no bead carryover as this can affect downstream steps. The supernatant should be completely clear*.
152. Quantify the concentration and visualize PCR products by running a 1:10 diluted sample on an Agilent D1000 ScreenTape device.
153. Submit for shallow low depth sequencing. *NOTE: >10k reads is sufficient to quantify plasmid distribution within the pool*.

#### Data analysis (timing: variable)

154. Search within `input.fastq.gz` for string CTTTNNNNGGTTAA and count the number of unique instances, where the NNNN represents the SRT barcode. The output is saved in a tab-delimited file named `output.txt`

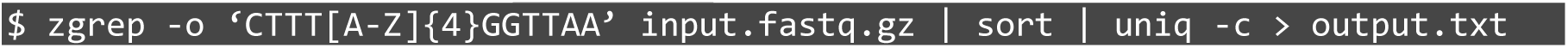

### COMMENTARY

#### Historical development of the protocol

Calling Cards was first developed and used to study TF binding in *Saccaromyces cerevisiae*. This yeast method uses a Ty5 integrase and engineered TF-Sir4 protein fusions to leave permanent “Calling Cards” where the engineered Sir4 has bound to the genome (Wang et al., 2008). To expand upon this technology, the transposase system was switched to the piggyBac transposon/transposase system due to its high transposition efficiency and activity across species (yeast, zebrafish, insects, mouse, and human), broadening the technology’s applicability (Wang et al., 2012). In these early protocols, to recover Calling Card insertions, genomic DNA was cleaved with restriction endonucleases, fragments were circularized, and amplified using inverse PCR from the transposon, and sequenced to capture transposon-genome junctions. The genomic sequences were then mapped to identify the location of the insertions. However, when Calling Cards began to be ported to complex systems like the mammalian brain, the small number of insertions per cell (<30) and the scale of the mammalian genome (∼3 Gb), made recovery of sufficient numbers of transposons challenging.

The development of self-reporting transposons (SRTs) solved this problem by having each insertion produce multiple copies of its own transposon-genome junction by transcribing RNA from within the transposon. Thus, insertions could be mapped from RNA with base-pair resolution using specialized RNA sequencing (RNA-seq). In parallel, and with the same starting material, standard RNA-seq can provide a simultaneous readout of gene expression from the same cells, in a manner compatible with single-cell sequencing technologies.

The SRT is a piggyBac transposon that contains an EF1α promoter driving a tdTomato reporter without a polyadenylation signal. When inserted in the genome, RNA polymerase II will transcribe the SRT (tdTomato) into the flanking genomic region until it reaches a cryptic polyadenylation signal or genomic polyadenine stretch. It was found that RNA-based libraries were much more sensitive than the DNA-based method and recovered nearly 50% more insertions (Moudgil et al., 2020b). Since each insertion produces multiple RNA copies of itself, one limitation of using this technology (outside of single-cell applications) was that one cannot differentiate 1) multiple independent insertions at the same locus across cells from 2) one insertion at the same locus whose RNA was sequenced multiple times. Initially this was resolved by preparing multiple independent aliquots of adjacent tissue for RNA extraction, followed by barcoding each sample by PCR (Cammack et al., 2020), thus exponentially amplifying the amount of sample handling. However, we more recently resolved this issue by introducing transcribed barcodes into the SRT itself. This feature dramatically reduces the required number of replicates, decreases cost and labor requirements, and increases the sensitivity of detecting multiple insertions at the same locus (Lalli et al., 2022). Finally, we have found that higher expression and copy number of the donor transposon is beneficial to maximize the number of insertions per cell, as transposons are more limiting than transposase activity (Yusa et al., 2011) (**Supplemental Figure 2A**), and now can recommend optimized ratios in this protocol.

The enhanced sensitivity of the SRTs relative to inverse PCR also enabled the application of Calling Cards to address questions in complex tissues, like the mammalian brain. Thus, to direct Calling Cards to genetically defined cell populations, we developed Cre-dependent hyPB transpose systems. We initially adopted a FLEx (or “<u>fl</u>ip-<u>ex</u>cision”) switch vector design so that a cassette encoding hyPB is in the antisense orientation and should not produce functional protein. In the presence of Cre (e.g., when delivered by AAV into a Cre-expressing mouse brain), recombination occurs, the sequence is flipped into the sense orientation, driving hyPB expression. However, with FLEx Calling Cards, we had noticed a low level of hyPB activity in Cre negative animals, suggesting some transcription of hyPB from the antisense strand of the AAV. To eliminate this background transposition, we redesigned the FLEx cassette to insert an intron into the middle of hyPB and position key LoxP sites within it. The vector thus contains the front half of the hyPB protein antisense to the second half of the protein. Thus, without Cre, both sense or antisense strands would not produce functional protein. This “FrontFlip-hyPB” construct was shown to effectively eliminate background transposition (Cammack et al., 2020), and the general design may be helpful in suppressing anti-sense leakiness in other Cre-dependent enzyme systems.

Thus, Calling Cards using barcoded SRTs, optionally combined with cell-type specific designs, are the latest reagents that can be used to record BRD4-bound enhancer usage or TF-DNA binding events in specific cell types in complex systems.

#### Applications of the method

Calling Cards has been successfully used to record BRD4-bound enhancer usage with unfused hyPB or transcription factor binding with custom TF-hyPB fusions *in vivo* and *in vitro* (Moudgil et al., 2020b; Wang et al., 2008, 2012; Cammack et al., 2020; Kfoury et al., 2021). Using this method, it was found that sex-biased BRD4-bound enhancer activity in glioblastoma (GBM) underlies sex differences in stem cell function and tumorigenicity in GBM. Pharmacological or genetic inhibition of BRD4 in male and female patient-derived GBM cell lines revealed that male GBM cells are more sensitive to BET inhibition and has important clinical implications (Kfoury et al., 2021). In addition, we have included a vignette here showing how Calling Cards can be used to record regional differences in enhancer usage across brain development. Additionally, the development of Cre recombinase dependent constructs with reduced background (ie. FrontFlip-hyPB) or use of cell type-specific promoters enables targeted recording in genetically defined populations (Cammack et al., 2020). While this protocol derives from our experience mainly on mouse neural tissue and cell types, tailoring Calling Cards for a particular cell type, tissue, or model organism may require optimization of reagent delivery, RNA isolation, and some PCR steps during library preparation. We also note that the tdTomato reporter within the SRT is useful to not only visualize cells but can also be used to enrich cell populations with FACS. We have developed a collection of modular Calling Card reagents that can be adapted to be used across a wide range of applications (see **Table 1** for list of available reagents). Calling Cards is positioned to generate unique datasets that enable the analysis of observed current cell states with historical molecular and epigenetic states.

#### Comparison with other methods

There is a myriad of genomic methods to investigate the epigenome, transcriptome, and chromatin state, and the assay of choice should be determined by the biological question (**Table 3)**. While many methods produce snapshots of states at tissue harvest, Calling Cards is one of two methods we know of that produces data that represents a cumulative catalog of protein-DNA interactions over time, using a genetically encoded system. The other method, DamID (Vogel et al., 2007), identifies binding sites of a protein by using a DNA methyltransferase (DAM) fusion protein, which methylates an adenine within a GATC sequence in close proximity to the binding site. Since adenine methylation does not occur naturally in eukaryotes, mapping the location of the methylated adenine nucleotides is then interpreted as a proxy for the binding site. The cumulative view of activity can be an appealing method to obtain the ground truth of TF binding or enhancer usage as this is recorded over time and is not sensitive to collection time. Further, since it is genetically encoded, it can be delivered to specific cell types. Since Calling Cards starts from an RNA sample, the mRNA transcriptome can be analyzed in parallel from the same cells and input material, allowing a measure of the final transcriptional state in parallel to the recorded TF binding profile. This approach is thus unique from DamID datasets, which cannot provide quantitative measures of transcription in addition to the record of DAM fusion protein-DNA interactions. RNA Pol II can be fused with DAM to profile transcription start sites and gene expression, but the ability to simultaneously profile a TF is lost.

**Table 3.**
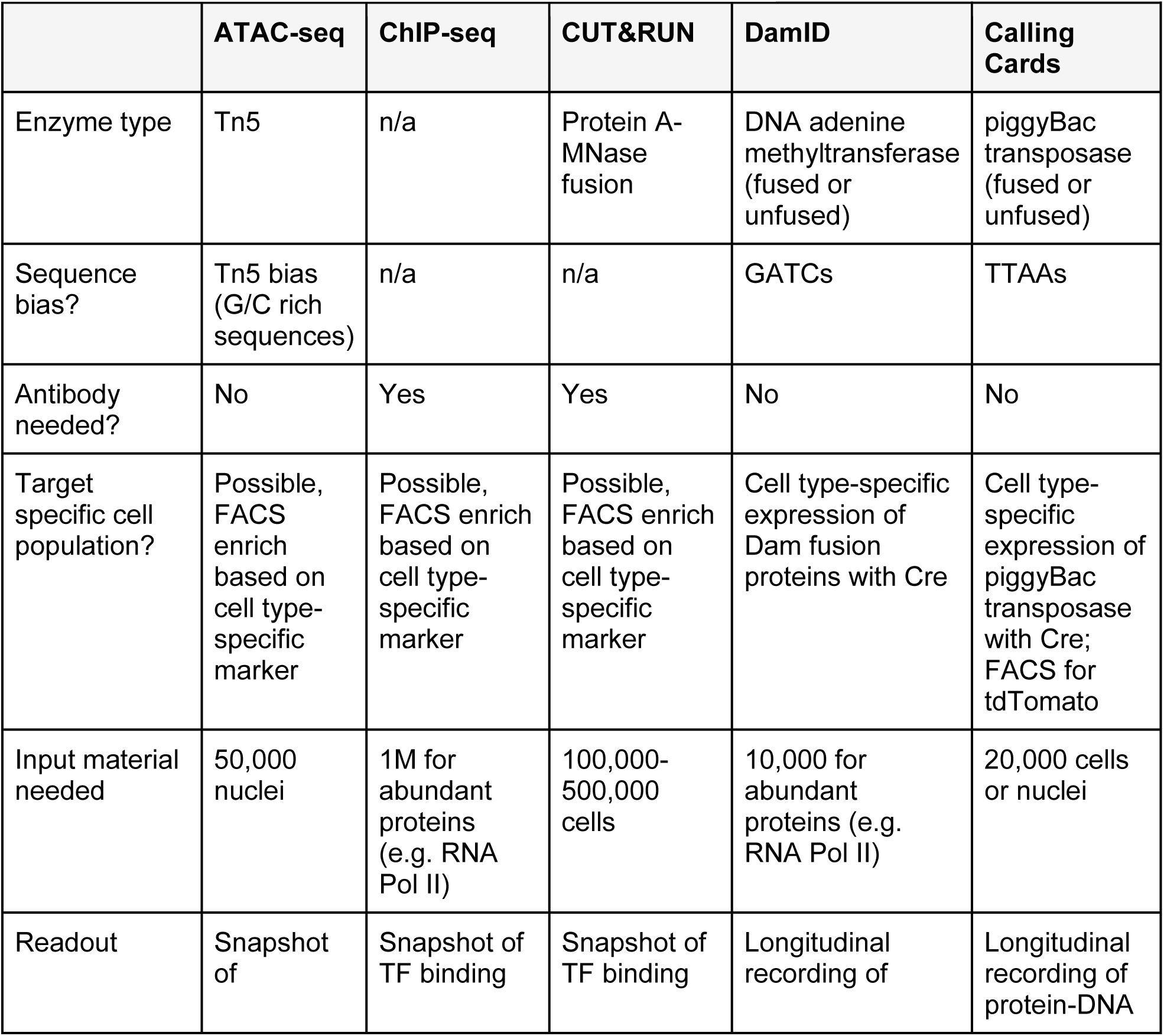
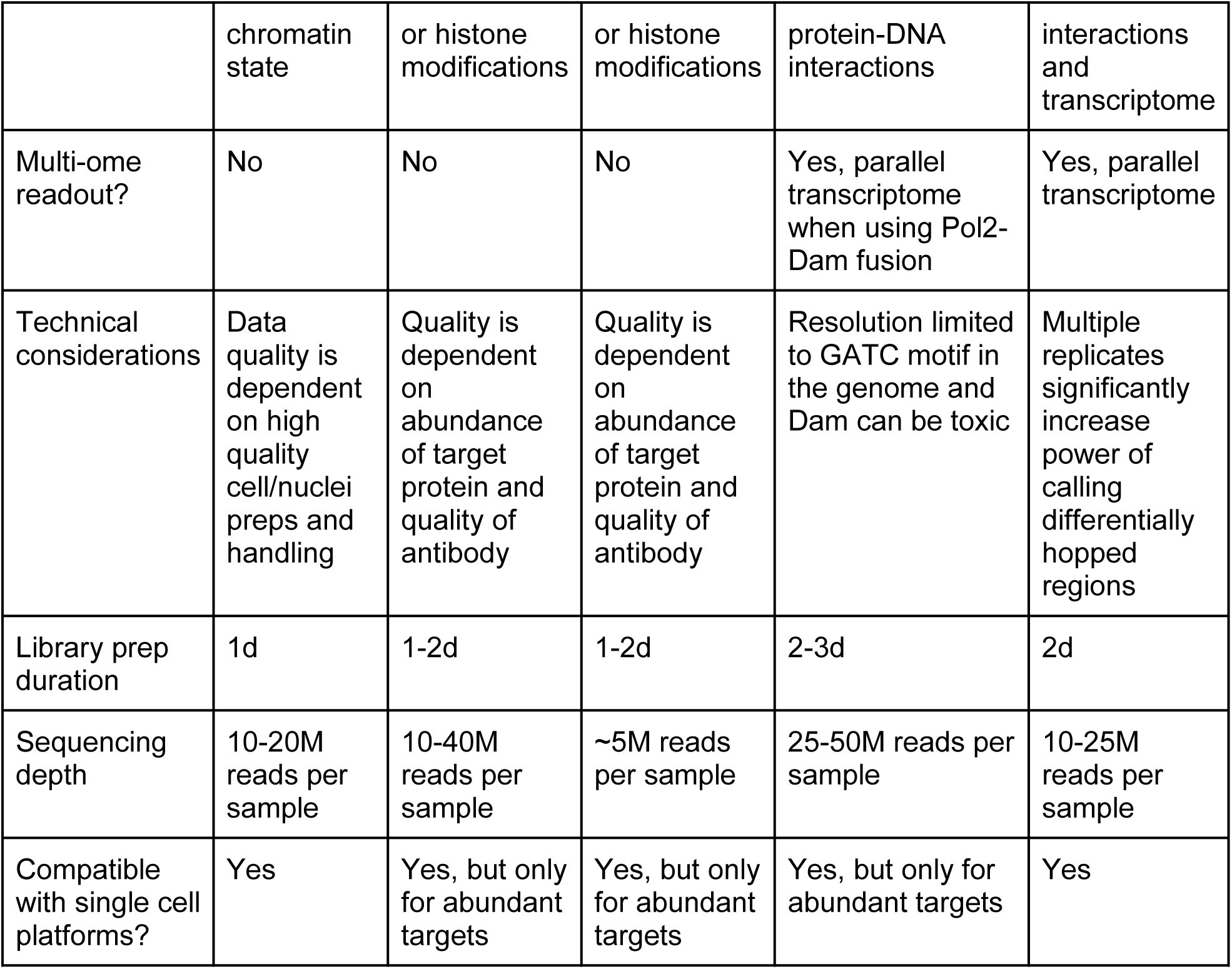
Comparison with other genomic methodologies

In comparison to non-recording methods of assessing DNA-protein interactions, in test cases examined so far, Calling Cards data is concordant with ChIP-seq data (Moudgil et al., 2020b). Thus, Calling Cards offers a powerful complementary approach to ChIP-seq/CUT&RUN and together, the resulting datasets can provide a more complete picture of TF biology and gene regulatory elements. Furthermore, experiments that require screening multiple time points, multiple TFs, or association of historical molecular events with eventual cell states are applications where one might prefer Calling Cards over ChIP-seq. However, for systems where genetic delivery of reagents is not an option (e.g., human brain), non-Calling Card methods are required. Finally, Calling Cards has been demonstrated to work both *in vitro* and *in vivo* and is a versatile tool that can be broadly used in models to study development and diseases that accumulate changes over time.

#### Troubleshooting

| Problem | Possible reason | Solution |
| --- | --- | --- |
| RNA is degraded or low quality (low RIN, low A260/A280 ratio) | Starting material was improperly handled/stored | Snap freeze tissue in liquid nitrogen. If starting with a frozen sample, do not let the sample thaw prior to RNA cleanup. If starting with fresh tissue, dissect quickly and minimize time to homogenization. |
|  | RNase contamination | Clean all surfaces, equipment, and gloves with RNase removing detergents. |
| Genomic DNA contamination | DNase treatment was insufficient | Repeat DNase treatment or switch from an on-column to an in-tube DNase I treatment method. |
| Low RNA yield | Overloaded homogenization | Homogenize samples in 1ml TRIzol per 100mg tissue. Scale volume up if needed. |
|  | Residual organic solvent contamination | Precipitate RNA and increase the number of ethanol washes. |
|  | Insufficient homogenization or lysis | Ensure no visible cell aggregates or tissue chunks remain during homogenization. |
| Low cDNA yield | Degraded RNA | Check RNA quality by BioAnalyzer or TapeStation. |
|  | Residual contamination from RNA purification steps | Precipitate RNA and wash with ethanol to remove EDTA, guanidinium, and proteases. |
|  | Low amount of RNA input | Increase the amount of starting RNA by combining biological replicates. |
| Low SRT yield | Low expression of Calling Card constructs | Increase the amount of plasmid or virus used to deliver Calling Card reagents. |
| Distinct banding in SRT QC | Low complexity library | Increase the amount of plasmid or virus used to deliver Calling Card reagents. |
| No product after bead cleanup | Failed bead cleanup | Use fresh 80% ethanol and do not overdry the beads. Do not disturb bead pellet during washes. |
|  | Insufficient PCR amplification | Increase the number of cycles for the indexing PCR step. |
| Low concentration (<1nM) of final library | Failed bead cleanup | Use fresh 80% ethanol and do not overdry the beads. Do not disturb bead pellet during washes. |
| Final library fragment size distribution not 200-1000bp | Failed tagmentation | Undertagmentation yielding larger products is a result of too much input material. Overtagmentation leads to smaller fragments and is a result of too little input material. Verify 600pg DNA input using a fluorometric quantification method. |
| Sequencing reads are the OM-PB primer sequence followed by G's | OM-PB indexing primer carryover | Beware of a ~100bp band in the electropherogram QC of the final library. Repeat bead cleanup and QC. |
| Low proportion of reads with adapter | PCR mispriming due to low library density | Combine biological replicates to increase signal (SRT fragments) to background. |
| Presence of overrepresented sequences (>0.1% of library) that is not biologically relevant | PCR mispriming (may correspond to prominent bands in SRT QC), genomic DNA contamination, primer carryover | Ensure proper cleanup of DNA with silica column or magnetic beads. |
| Low number of recovered insertions | Low expression of Calling Card constructs | Increase the amount of plasmid or virus used to deliver Calling Card reagents. |
|  | Low library complexity | If starting from a rare cell population, purify the target population by FACS or enrich their RNA by a method like TRAP (Heiman et al., 2014). Otherwise add more biological replicates |
| Poor sequencing coverage of each insertion (<5 reads/unique insertion) | Insufficient sequencing depth | Sequence deeper. |

#### Understanding Results

##### Interpretation of quality control checkpoints

Considering the protocol length and cost of reagents, careful quality control throughout is recommended to ensure high quality datasets and efficient use of reagents and time. During the development and optimization of the protocol, we have identified key parameters that are important for the successful generation of high quality sequencing libraries and data. These include the amount of starting material, amount of DNA input into each PCR step, amount of final library, and the number of recovered insertions. Interestingly, we found relatively permissive limits of variation for QC steps, meaning that the protocol is robust at recovering insertions given proper library preparation steps. We generally found that maximization of input material at each step that allows a range of input leads to the most recovered insertions. For exceptionally complex libraries, multiple library preparations from the same starting material may be necessary to capture their full diversity.

The first QC checkpoint is quantifying the integrity of the purified total RNA. A minimum RIN of 8 with minimal signs of degradation is recommended for most cases, however this can vary depending on the type of sample. *In vitro* cell samples can be as high as 10, whereas FFPE samples will typically be low. RINs lower than 7 generally do not sequence well, however purification/enrichment kits can be used prior to the library preparation. In addition to RIN, genomic DNA contamination can be seen as an additional peak above the 28S band or as any unexpected signal between the rRNA peaks. The genomic DNA can not only cause inaccurate RNA quantification, but can also interfere with downstream steps, so an additional DNase I treatment (in addition to the DNase I treatment during RNA cleanup) is recommended prior to proceeding further.

The next QC step is the concentration quantification of single stranded cDNA following the first strand synthesis. The presence of the SMART_dT18VN RT primer during the subsequent PCR amplification of SRTs has been found to increase nonspecific products likely due to priming to endogenous polyA stretches. Thus, complete removal of primer using a simple spin column-based PCR purification kit is an important step prior to PCR amplification of SRTs. Library density assessment of tdTomato containing cDNA fragments using quantitative RT-PCR is an optional but valuable QC step. Here, the samples can be initially screened for the abundance of tdTomato and determined if certain samples not passing a threshold should be omitted from subsequent library preparation steps (**Figure 7A-C**). In our experience, C_T,tdTomato_>30 and tdTomato expression values normalized to B-actin that are <0.1 will still make libraries, however, the number of recovered insertions is minimal. The specific values will likely vary based on sample type and transfection/transduction efficiency and will require some initial empirical testing, but the potential benefit of saving labor and reagent costs is worthwhile.

Accurate quantification of ssDNA using a fluorometric method is highly recommended to maximize the amount of input material (up to 100ng) that goes into the next PCR reaction to amplify tdTomato containing SRTs (**Figure 8**).

After bead cleanup of SRTs, the electropherogram of the amplified SRTs should resemble **Figure 9A,B**. A mostly smooth distribution of products with a bias towards larger products should be expected from 300-5000bp. Some banding has been observed to occur and has not been found to affect library quality, however, strong and distinct bands without larger PCR products likely indicates a low number or diversity of SRTs, or can be artifacts resulting from PCR overamplification (**Supplemental Figure 6**).

The final QC step is to quantify the library concentration and fragment size distribution. The tagmented library should be a smooth distribution between 200-1000bp (Figure 9C,D). Presence of a ∼100bp band indicates insufficient removal of the OM-PB primer and another round of bead cleanup is recommended to remove the primer as this can have a detrimental effect on sequencing (**Supplemental Figure 6**).

#### Computational analysis of Calling Card data

Analysis can be broken down into three main stages: 1) generation of the qBED files, 2) peak calling and insertion counting, and 3) downstream analysis. The workflow and steps involved for the first two stages are diagrammed in **Figure 2B**. Further downstream analysis (e.g. differential peaks between samples, motif enrichment, gene ontology analysis) is not covered in this protocol. First, the raw reads are prepared by filtering for reads containing SRT barcodes with `UMItools`, trimming adapter sequences with `trimmomatic`, and standard quality control such as per base sequence quality scores and per base sequence content with `FastQC`. The reads are then aligned and the SRT insertions are mapped to the reference genome. The modularity of the `nextflow` pipeline enables the end user to easily select the desired alignment software. The default aligner is `bwamem2`. A full list of preconfigured `nf-core` modules can be found at https://nf-co.re/modules. In the `sam` file, the header of aligned reads is tagged with the OM-PB barcode, index1/i7, index2/i5, and SRT barcode, allowing us to identify the insertions’ sample of origin. The resulting bam file is indexed and parsed into a `qBED` file which lists the unique insertions. This file can be used as input into peak callers to call peaks for genomic regions that are enriched with Calling Card insertions.

#### Sequencing saturation

Following an initial sequencing of a sample, one can determine if deeper sequencing would be useful to recover more binding events. For Calling Cards data, sequencing saturation is a measure of how many times an insertion has been read. This provides a way to estimate how much new information (ie. unique insertions) is likely to be gained by sequencing deeper. Within each `qBED` file, the fourth column represents the number of reads associated with each insertion. To evaluate saturation, `bam` files were downsampled across set ratios (0.001, 0.003, 0.01, 0.03, 0.1, and 0.3 of the full library) using `samtools` to simulate shallower sequencing.

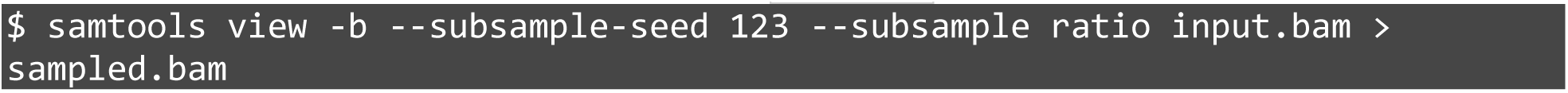

An example plot of downsampled samples with varying amounts of insertions is shown in **Figure 10A**. A final recommended sequencing depth can be estimated where the plots start to asymptote. Peaks were called on each of the downsampled files (**Figure 10B,C**). While the size of peaks was not as sensitive to the number of insertions, the number of called peaks is directly proportional to the number of insertions and reads, as expected. It is also evident that the number of reads per insertion also increases with the number of reads (**Figure 10D**). A left-skewed cumulative density indicates under-sequencing as there are many insertions that were only sequenced once. A shift to the right with a relatively low number of insertions with low coverage demonstrates that the sample is approaching sequencing saturation. With deeper sequencing, the resolution of the called peaks increases. This is evident based on the observation that the fraction of overlapping peaks does not change upon downsampling, despite the dramatic increase in the number of peaks (**Figure 10E**).

#### Assessing reproducibility of across replicates

To assess the reproducibility of Calling Cards data across biological and/or technical replicates, the `qBED` files for each experimental condition or genotype can be concatenated to create a file containing all insertions. Each insertion is tagged with unique index sequences and can be used to identify the sample of origin. Peaks are called on the aggregated insertions. The similarity between a pair of replicates can be analyzed by plotting the number of insertions (normalized to sequencing depth) found within each peak region from each of the replicates as a scatterplot, with values from one replicate on the x axis, and the other on the y axis. To assess multiple replicates, a scatterplot matrix can be plotted. Replicates with high similarity should have a symmetrical linear correlation (**Supplemental Figure 4**).

#### Peak calling and annotation

There are several algorithms available for peak calling on Calling Cards data and they are all contained in the py-callingcards package (reference, paper being written) available at https://github.com/The-Mitra-Lab/pycallingcards and detailed documentation with tutorials can be found https://pycallingcards.readthedocs.io/en/latest/. Briefly, all qBEDs from each sample are combined to call a joint set of peaks. Within the `nf-core/callingcards` pipeline, `CCcaller` is the default peak caller that first identifies pileups of insertions given the number of TTAA sites, then uses Poisson tests to identify regions that are enriched with insertions over the background. This is an optimized implementation of `CC_tools`, which is based on the `MACS` algorithm (Zhang et al., 2008). `Blockify` is another peak caller based on Bayesian blocks and is a noise-tolerant algorithm that segments the genome into non-overlapping regions with a constant number of insertions and similarly uses Poisson tests to identify significant peaks using a p value cutoff of 10^-62^ as described in (Moudgil et al., 2020b). Once the peaks are called, BEDtools is used to find the two closest genes to each peak region (Quinlan and Hall, 2010).

#### Differentially hopped regions

Differential peak analysis is performed using the `cc.tl.rank_peak_groups` function within py-callingcards (reference, paper being written). Briefly, two-sided Fisher’s exact test or the binomial test is used to test for peaks that are differentially hopped with significant log-fold changes across the compared groups. Differentially hopped regions (|LFC|>1, padj<0.01) can be identified and used for downstream analysis.

#### Multi-omics data integration

Using a TF-hyPB fusion transposase, Calling Cards records cumulative TF binding events. If homologous high-quality ChIP-seq datasets are available for the TF of interest, Calling Card data can be directly compared to assess the degree of overlap between the orthogonal methods. Depending on how similar the ChIP-seq experimental conditions are, one should expect a reasonable amount of intersection, keeping in mind that Calling Card data may yield more peaks given that it records over time. When using Calling Cards with the unfused piggyBac transposase, BRD4-bound enhancers are recorded and can be similarly validated with BRD4 ChIP-seq. Additionally, it can be informative to compare these putative enhancers with ATAC-seq, H3K27ac, and H3K4me1 ChIP-seq peaks which reveal open chromatin, active enhancers, and promoters (Cammack et al., 2020), as BRD4 ChIPseq can correspond somewhat to these. Generally, these Calling Cards peaks should be de-enriched in regions with repressive H3K27me3 marks. The combination of these datasets can be used to establish enhancer-gene regulatory interactions, which can be further confirmed with chromosome conformation capture technologies such as Hi-C or HiChIP.

Comparison to RNA-seq data is also sometimes informative especially when looking at promotor-localized peaks (and keeping in mind the usual caveats and challenges of linking enhancer regions to specific genes). Differentially expressed genes can be identified from bulk RNA-seq data using DEseq2 (Love et al., 2014) and correlated with TF binding or enhancer usage Calling Cards peaks. Associating these TF binding-gene pairs can be useful to those studying gene regulation. The `cc.tl.pair_peak_gene_bulk` function in py-callingcards identifies annotated genes associated with differential Calling Cards peaks are also significantly differentially expressed.

#### Files to submit for publication

Finally, with analysis completed, as with all high-throughput sequencing data, Calling Cards data should be uploaded to a publicly accessible repository such as Gene Expression Omnibus (GEO) concurrent with publication of corresponding manuscripts. To be maximally useful to the community, each submission should include a metadata spreadsheet, all raw `FASTQ` files prior to any processing (QC filters, adapter trimming, etc.), and processed data files. These should include insertions (`qBED`), genomic coordinates of called peaks (`BED`), density tracks of insertions (`bedGraph`), insertions per peak counts matrix for all samples in the study, and a list of differentially hopped regions. The `qBED`, `BED`, and `bedGraph` files are useful as they can then be used by any user to visualize the data on the WashU Epigenome Browser, or for comparison to their own datasets. We also recommend including a summary table in publications, reporting sequencing metrics, QC metrics, and the number of recovered insertions (example shown in **Table 5**).

**Table 4.** Software versions used in `nf-core/callingcards`.

| Process Name | Software | Version |
| --- | --- | --- |
| BWAMEM2_MEM | bwamem2 | 2.2.1 |
|  | samtools | 1.16.1 |
| CUSTOM_DUMPSOTWAREVERSIONS | python | 3.10.6 |
|  | yaml | 6.0 |
| FASTQC | fastqc | 0.11.9 |
| GTF2BED | perl | 5.26.2 |
| HOPS_MAMMALS | pycallingcards | 0.0.2 |
| PICARD_COLLECTMULTIPLEMETRICS | picard | 2.27.4-SNAPSHOT |
| SAMPLESHEET_CHECK | python | 3.8.3 |
| SAMTOOLS_FLAGSTAT | samtools | 1.16.1 |
| SAMTOOLS_IDXSTATS | samtools | 1.16.1 |
| SAMTOOLS_INDEX | samtools | 1.16.1 |
| SAMTOOLS_STATS | samtools | 1.16.1 |
| TRIMMOMATIC | trimmomatic | 0.39 |
| UMITOOLS_EXTRACT | umitools | 1.1.2 |
| Workflow | nextflow | 22.10.4 |
|  | nf-core/callingcards | 0.0.0-dev |

**Table 5.** Summary of bulk Calling Cards experiments

| Sample | Constructs | Biological replicates | Insertions | Reads | Mean coverage |
| --- | --- | --- | --- | --- | --- |
| Mouse cortex | AAV9-hyPB<br>AAV9-tdT-SRT_bc | 3 | 2,464,951 | 149,405,068 | 60.6 |
| Mouse midbrain | AAV9-hyPB<br>AAV9-tdT-SRT_bc | 3 | 1,453,903 | 123,980,679 | 85.3 |
| Mouse hindbrain | AAV9-hyPB<br>AAV9-tdT-SRT_bc | 3 | 645,633 | 84,644,681 | 131.1 |

#### Time Considerations

Basic Protocol 1: Preparation and delivery of Calling Cards reagents

Calling Cards plasmid preparation (timing: variable)
Delivery of Calling Cards reagents (timing: variable)
Intracerebroventricular injection (timing: 30 min-1 hr)
Basic Protocol 2: Sample preparation

Harvesting in vitro cultures (timing: variable)
Harvesting brain tissue (timing: 20 min/mouse)
Basic Protocol 3: Sequencing library preparation

RNA purification (timing: 60 min)
First strand cDNA synthesis (timing: 2 hr)
Library density qPCR (timing: 1.5 hr)
Amplification of self-reporting transcripts (timing: 3 hr)
Bead cleanup of PCR products and QC (timing: 30-45 min)
Tagmentation and indexing PCR (timing: 1-1.5 hr) Bead cleanup of PCR products and QC (timing: 30-45 min)
Basic Protocol 4: Library pooling and sequencing

Library quantification by qPCR (timing: 1.5 hr)
Library pooling (timing: 15 min) Sequencing (timing: variable
Basic Protocol 5: Data analysis

Install and configure nextflow (timing: variable)
Run nf-core/callingcards on your own data (timing: variable)
Visualization of Calling Cards data (timing: variable)
Support Protocol 1: NGS quantification of barcode distribution within plasmid poo

Isolation of viral genome from AAV particles (timing: 3.5-4 hr)
Library preparation (timing: 1h)
Bead cleanup and quantification (timing: 30 min)
Data analysis (timing: variable)

## DATA AVAILABILITY STATEMENT

The data that support the protocol are available in Gene Expression Omnibus (GEO) at https://www.ncbi.nlm.nih.gov/, reference number GSE223926.

## AUTHOR CONTRIBUTION STATEMENTS

*Conceptualization:* AY, CM, AM, AJC, RDM, and JDD; *Methodology:* AY, CM, JG, XC, AM, and AJC; *Software:* AY, CM, JG, AM, and RDM; *Validation:* AY, AM, and AJC; *Formal analysis:* AY, CM, SS, and MAG; *Investigation:* AY, SS, MAG, JH, and MC; *Resources:* MRB, RDM, and JDD; *Data curation:* AY, CM, and JG; *Writing-original draft:* AY and SS; *Writing-review and editing:* AY, CM, SS, MAG, JG, XC, JH, MC, MRB, RDM, and JDD; *Visualization:* AY, CM, SS, and MAG; *Supervision:* AY, RDM, and JDD; *Project administration:* AY, RDM, and JDD; *Funding acquisition:* AY, SS, MAG, MRB, RDM, and JDD.

## ACKNOWLEDGMENTS

We would like to thank the Genome Technology Access Center at the McDonnell Genome Institute (GTAC@MGI) for their sequencing expertise and services. This work was supported by the Hope Center Viral Vectors Core at Washington University School of Medicine. Imaging was performed in part through the use of the Washington University Center for Cellular Imaging (WUCCI). This work was supported by National Institute of Mental Health (RF1MH117070, RF1MH126723 to RDM and JDD), National Institute of General Medical Sciences (35GM141012 to MRB), National Human Genome Research Institute (T32HG000045 to AY), Autism Science Foundation (22-007 to SS), and National Institute of Neurological Disorders and Stroke (T32NS121881 to MAG).

## INTERNET RESOURCES

https://nf-co.re/callingcards: This is the official nf-core community page that hosts the bioinformatics pipeline. Complete documentation and release notes can be found here.

https://github.com/nf-core/callingcards/issues: Found a bug? Have a feature request? We welcome any submissions big or small through github.

https://nfcore.slack.com/channels/callingcards: This is the official slack channel that is monitored by the developers and authors. Feel free to drop in to ask questions or just say ‘hi’!

https://www.addgene.org/kits/mitra-barcoded-transposon/: This is a link to an Addgene plasmid kit that contains individual barcoded self-reporting transposons. These can be grown up and pooled into one large pool or multiple subpools.

