## Supplemental figures for "Calling Cards: a customizable platform to longitudinally record protein-DNA interactions over time in cells and tissues"

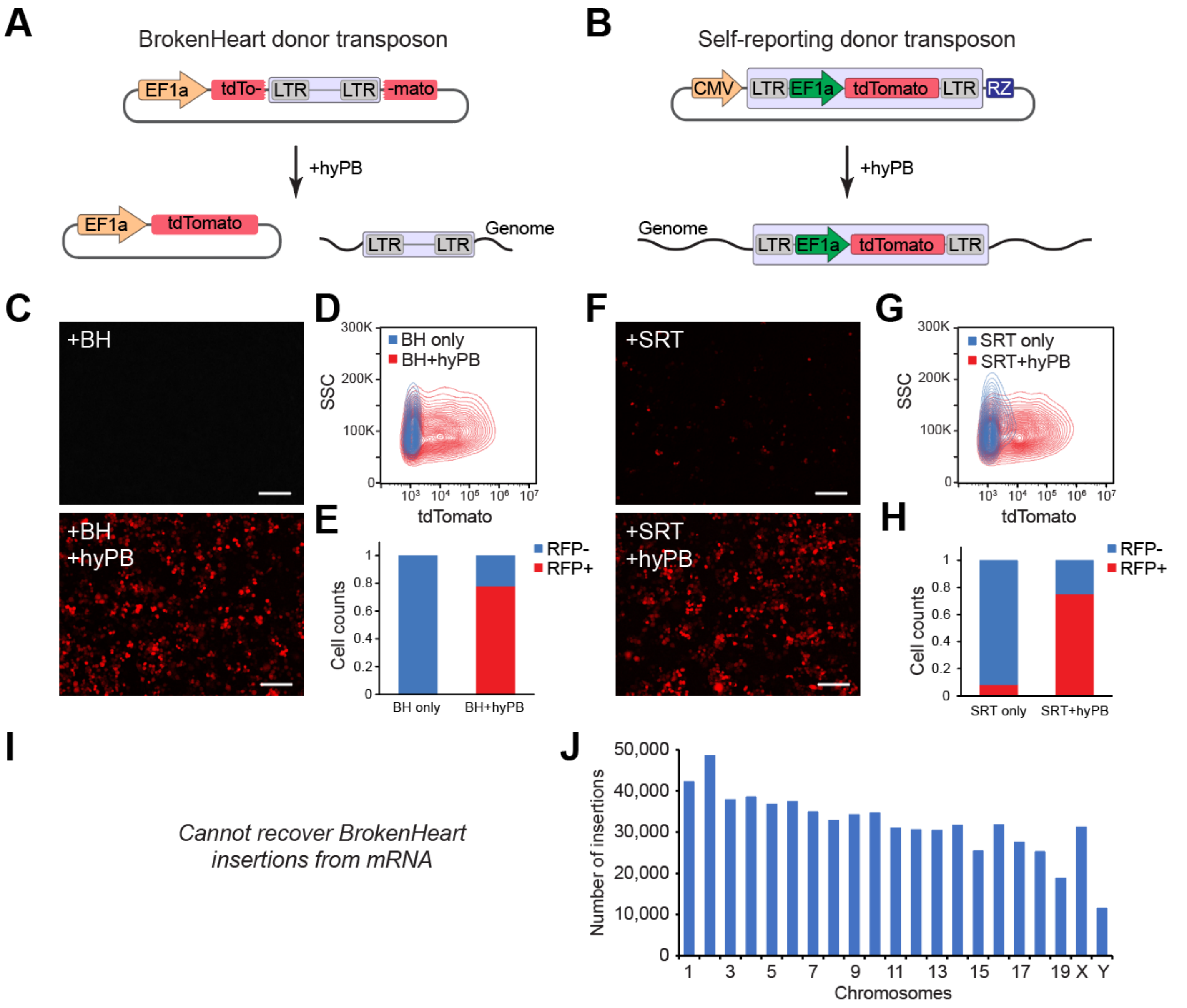


**Supplemental Figure 1. Comparison of BrokenHeart (BH) and self-reporting transposons (SRT). (A,B)** Schematics showing differences in BrokenHeart and SRT transposition mechanisms. TdTomato fluorescence is detectable only with transposase activity in BrokenHeart conditions, while some background is observed with SRTs. **(C)** Representative images of HEK293 cells transfected with BrokenHeart only or BrokenHeart+hyPB. Scale bar: 50um. **(D)** Contour plot showing virtually no tdTomato fluorescence in donor only control. **(E)** Quantification of cell proportions of RFP negative and RFP positive cells. **(F-H)** Similar analysis as **C-E**, but with SRT donor instead of BrokenHeart. Scale bar: 50um. **(I)** Sequencing libraries cannot be made from BrokenHeart Calling Card libraries, whereas libraries from SRT Calling Cards **(J)** can be prepared from mRNA.

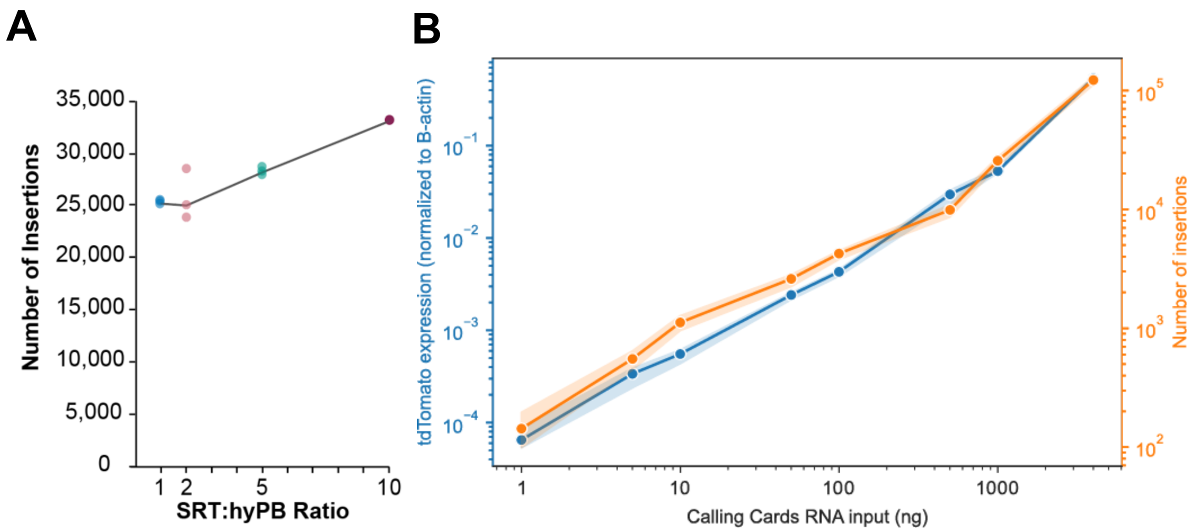


**Supplemental Figure 2 Optimization of experimental conditions to maximize Calling Cards insertions. (A)** Normalized insertions per 50k reads at four ratios of self-reporting transposon (SRT) to hyper piggyBac transposase (hyPB) (1:1, 2:1, 5:1, and 10:1) show that increasing transposon availability increases recovery of insertions in HEK293 cells. **(B)** TdTomato expression determined by quantitative RT-PCR (blue) and recovered insertions (orange) as a function of total RNA (4ug) spiked with a range of RNA containing Calling Cards insertions.

```
                         Index1                                 Primer barcode
5′- AATGATACGGCGACCACCGAGATCTACAC[i5]ACACTCTTTCCCTACACGACGCTCTTCCGATCTNNNCGTCAATTTTACGCAGACTATCTTT
        Illumina P5 adapter                     Truseq Read1                       piggyBac LTR
```

**Supplemental Figure 3. Sequence and structure of OM-PB primer.**

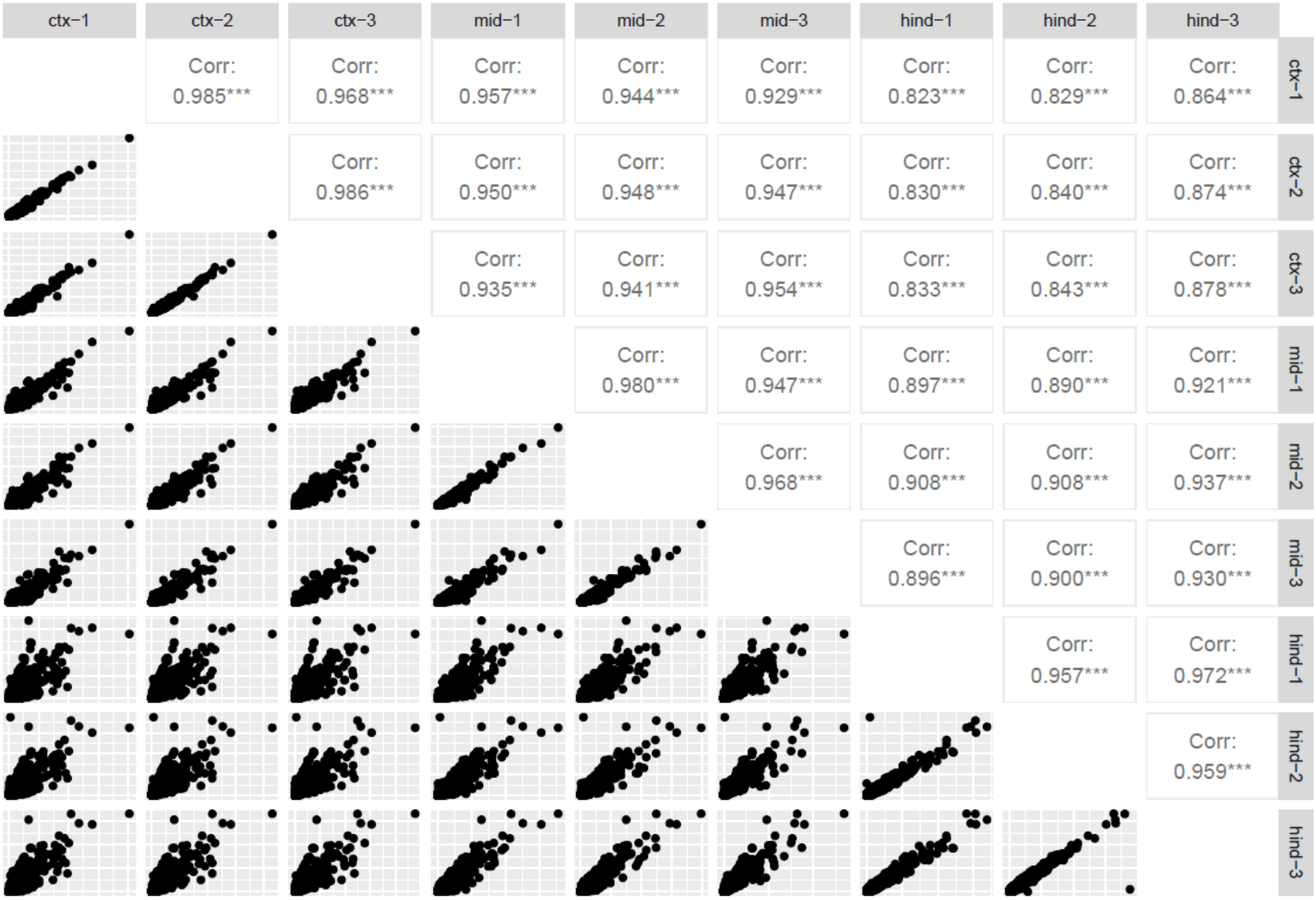


**Supplemental Figure 4. Scatterplot correlation matrix of biological replicates of cortex, midbrain, and hindbrain samples.**

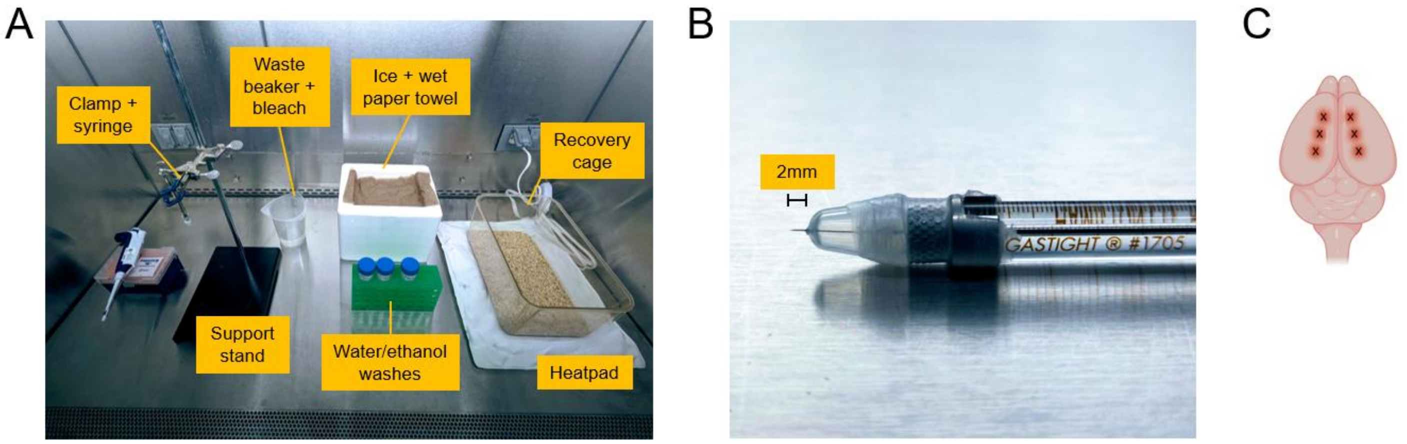


**Supplemental Figure 5. Setup for intracerebroventricular (ICV) injections. (A)** An example layout of materials and equipment that are needed for neonatal ICV injections of AAVs within a biosafety cabinet. **(B)** Close-up photograph of the hamilton syringe and custom needle guard to ensure a consistent injection depth of ~2mm. **(C)** Cartoon schematic depicting the approximate anatomical locations of the injections for a single animal.

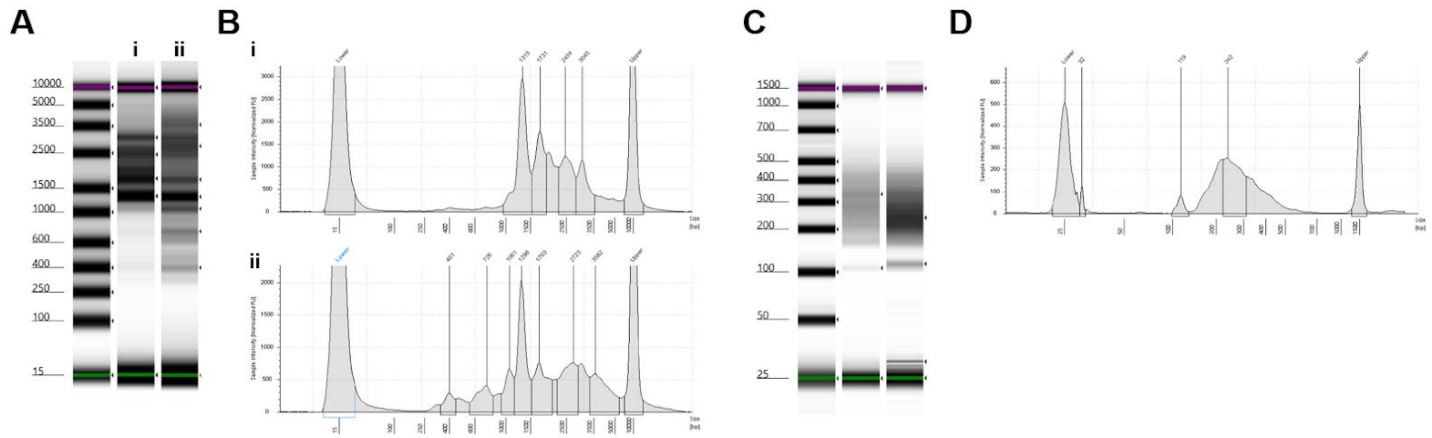

**Supplemental Figure 6. Tapestation traces of samples that do not pass QC.** Panel **(A)** shows the gel image of samples with abnormal SRTs. Panel **(B)** shows the electropherogram of the two samples. There is a lack of the distribution as seen in Figure 9A,B. Panel **(C)** shows the gel image of tagmented libraries where lane 2 does not pass QC due to the strong presence (>5% area of total region) of a ~120bp peak. Panel **(D)** shows the electropherogram of lane 2.
